# Benefits and Limits of Phasing Alleles for Network Inference of Allopolyploid Complexes

**DOI:** 10.1101/2021.05.04.442457

**Authors:** George P. Tiley, Andrew A. Crowl, Paul S. Manos, Emily B. Sessa, Claudia Solís-Lemus, Anne D. Yoder, J. Gordon Burleigh

## Abstract

Accurately reconstructing the reticulate histories of polyploids remains a central challenge for understanding plant evolution. Although phylogenetic networks can provide insights into relationships among polyploid lineages, inferring networks may be hindered by the complexities of homology determination in polyploid taxa. We use simulations to show that phasing alleles from allopolyploid individuals can improve phylogenetic network inference under the multispecies coalescent by obtaining the true network with fewer loci compared to haplotype consensus sequences or sequences with heterozygous bases represented as ambiguity codes. Phased allelic data can also improve divergence time estimates for networks, which is helpful for evaluating allopolyploid speciation hypotheses and proposing mechanisms of speciation. To achieve these outcomes in empirical data, we present a novel pipeline that leverages a recently developed phasing algorithm to reliably phase alleles from polyploids. This pipeline is especially appropriate for target enrichment data, where depth of coverage is typically high enough to phase entire loci. We provide an empirical example in the North American *Dryopteris* fern complex that demonstrates insights from phased data as well as the challenges of network inference. We establish that our pipeline (PATÉ: Phased Alleles from Target Enrichment data) is capable of recovering a high proportion of phased loci from both diploids and polyploids. These data may improve network estimates compared to using haplotype consensus assemblies by accurately inferring the direction of gene flow, but statistical non-identifiability of phylogenetic networks poses a barrier to inferring the evolutionary history of reticulate complexes.

## Introduction

Polyploidy, or whole-genome duplication, occurs throughout the eukaryotic tree of life. However, its evolutionary significance is especially evident in plants, with recent estimates suggesting up to 35% of vascular plant species are of recent polyploid origin (Wood et al. 2009; Barker et al. 2016). Despite advances in genomic data generation and numerous studies investigating the role of whole-genome duplication in plant speciation and local adaptation (reviewed in Soltis et al. 2014), polyploidy remains a central challenge for the field of phylogenetics. One persistent problem when analyzing sequence data from polyploid taxa, and especially allopolyploids, is identifying the alleles and divergent homeolog copies from parental lineages. Most bioinformatic tools for processing next generation sequence data were developed with diploids, or specifically humans, in mind. These approaches often collapse variable homeolog sequences into a single consensus sequence for *de novo* assemblies or assume the organism is diploid when performing genotyping and phasing for reference-based assembly. For polyploids, this can create chimeric sequences that may interfere with phylogenetic reconstruction and obscure evidence of polyploidy and its origins. Using allelic data that more accurately capture the complex genomic histories of polyploids should enable analyses to incorporate divergent signals from polyploid loci into phylogenomic inference, distinguish allopolyploidy from autopolyploidy, and identify parental taxa. However, few studies have examined the potential benefits of using phased versus unphased data to reconstruct polyploid histories (Kamneva et al. 2017; Oberprieler et al. 2017; Eriksson et al. 2018), and there are few formal methods and little guidance for phasing alleles from polyploid taxa. Here we explore the value of using phased data to reconstruct polyploid networks, leveraging fast polyploid phasing algorithms (Xie et al. 2016) to develop a bioinformatic pipeline that can phase alleles from polyploids using target enrichment sequence data.

Previous studies have suggested phasing alleles is crucial for accurate evolutionary reconstruction of reticulate complexes, at least when sampling relatively few loci (e.g., 4 to 10; Rothfels et al. 2017; Eriksson et al. 2018). However, it remains challenging to genotype (e.g. Blischak et al. 2018b) and phase next generation sequencing data from polyploids using short reads, in which individual variants may lack physical linkage information (e.g. Schrinner et al. 2020). Methods exist to genotype consensus loci from target enrichment data, but these have been either limited to diploids (Kates et al. 2018; Andermann et al. 2019) or used manual curation of variants with polyploids where both parental populations are available (Eriksson et al. 2018). Otherwise, obtaining phased sequence data for polyploids has largely depended on long-read sequencing to recover complete haplotype sequences (e.g., Rothfels et al. 2017) or cloning PCR products (e.g., Sessa et al. 2012; Oberprieler et al. 2017).

Target enrichment (or HybSeq), in which specific regions of the genome are isolated and sequenced (Albert et al. 2007; Gnirke et al. 2009; Faircloth et al. 2012; Lemmon et al. 2012), is an increasingly common method for collecting large-scale phylogenomic datasets, and these data can provide insights into the evolutionary history of reticulate complexes (e.g., Crowl et al. 2020; Karimi et al. 2020) and sources of gene tree discordance (e.g., Crowl et al. 2017; Morales-Briones et al. 2018; Stull et al. 2020). Probe kits for target enrichment have been developed in many land plant lineages (Wolf et al. 2018; Johnson et al. 2019; Liu et al. 2019; Breinholt et al. 2021), and there are bioinformatic pipelines available for custom probe design (e.g., Jantzen et al. 2020). The most common approach to assemble phylogenetic datasets from target enrichment data has been to use haplotype consensus assemblies from various pipelines (e.g. Faircloth 2016; Johnson et al. 2016; Andermann et al. 2018; Breinholt et al. 2018). The *de novo* assembly algorithms within these pipelines often consider only the most frequent nucleotide sequence and discard the alternatives (Bankevich et al. 2012; Iqbal et al. 2012; Luo et al. 2012). This results in loci in which variable positions are collapsed to a single base call (haplotype consensus loci), losing information related to heterozygosity. While this may be appropriate, or at least benign, for phylogenetic analyses of diploid taxa (Kates et al. 2018), it may pose substantial problems when attempting to investigate the evolutionary history of polyploid taxa or reticulate lineages. Some of the before-mentioned pipelines can recover heterozygosity information and phase sequences by mapping reads back to their haplotype consensus sequences (Andermann et al. 2018), but this ability is, at the time of writing, limited to diploids.

Recovering phased haplotype sequences from polyploid individuals, especially allopolyploids with distinct subgenomes, should be valuable when estimating phylogenetic networks that likely are more accurate representations of reticulate evolution than a bifurcating tree. Simulations and empirical analyses have suggested that phylogenetic networks can recover the reticulate histories of polyploid lineages with few loci, at least when gene tree discordance due to incomplete lineage sorting (ILS; Hudson et al. 1983; Pamilo and Nei 1988) is low (Oberprieler et al. 2017) using parsimony methods (Huber et al. 2006; Lott et al. 2009), or with moderate ILS when it is explicitly modeled (Jones et al. 2013). Contemporary phylogenetic network models and software packages jointly consider gene tree variation due to allele sampling error as described by the multispecies coalescent (MSC; Rannala and Yang 2003) and gene flow modeled as episodic introgression events (Solis-Lemus and Ané 2016; Wen et al. 2016; Zhang, Ogilvie et al. 2018, Flouri et al. 2020). Depending on the complexity and goals of the research question, these methods can search for networks with a constrained number of reticulation events using quartet-based maximum pseudolikelihood (Solis-Lemus and Ané 2016; Wen et al. 2018) or a full-likelihood Bayesian model where the number of reticulations is a parameter (Wen et al. 2018; Zhang et al. 2018). Also, it is possible to estimate model parameters (e.g., divergence times, population sizes, and the fraction of introgressed genes) on a fixed species network using a full-likelihood Bayesian model that allows efficient computation with large numbers of loci (Flouri et al. 2020). We refer to these network models from here on as the multispecies coalescent with introgression (MSci), consistent with Flouri et al. (2020), although other names have been used, such as the network multispecies coalescent (NMSC; Zhu and Degnan 2017) and multispecies network coalescent (MSNC; Wen et al. 2016). We emphasize that using network approaches to investigate polyploid complexes is not novel (e.g., Huber et al. 2006; Lott et al. 2009; Jones et al. 2013; Crowl et al. 2017), but the difficulty of collecting appropriate genomic data from polyploids for such analyses has limited their use.

To address the issues outlined above, we have developed a pipeline, PATÉ (Phased Alleles from Target Enrichment data), that can phase genotyping data for individuals of a known ploidy without the need for sampling their parental lineages. PATÉ was designed with scalability and population-level sampling in mind for target enrichment projects where deep coverage from paired-end Illumina data allows calling of high-quality variants. In this study, we first use simulations to explore the ability of network approaches to reconstruct the history of allopolyploidy in the presence of ILS, and whether phasing the data affects the accuracy of the reconstruction. We show that using phased allelic data can improve network estimation and divergence time estimation compared to using haplotype consensus sequences, but also highlight scenarios where phasing may not be necessary or beneficial. For an empirical example of how PATÉ output can be used in studies of reticulate evolution, we compared phased (either all alleles or only one) and unphased (either haplotype consensus or IUPAC ambiguity codes) data to infer the evolutionary history of the North American *Dryopteris* fern complex, a model system for reticulate polyploid evolution (Sessa et al. 2012a; Sessa et al.

2012b), using new targeted enrichment data. The system includes four diploid species, as well as one extinct diploid, that have formed five allopolyploids in which there is high confidence in the parent-progeny relationships (Fig. 1), although numerous sterile allopolyploid species have also been reported within the complex (Montgomery and Paulton 1981). The allopolyploid species have relatively ancient origins, with the best estimates placing hybridization events between six and 13 Ma (Sessa et al. 2012b). PATÉ is capable of recovering phased haploid sequences almost always the length of the original haplotype consensus assembly from polyploid individuals, although the switch errors among these haplotype sequences is unknown. Networks inferred from phased data more accurately represent some relationships within the North American *Dryopteris* complex than unphased strategies, but statistical identifiability problems confound all data types.

**Figure 1.**
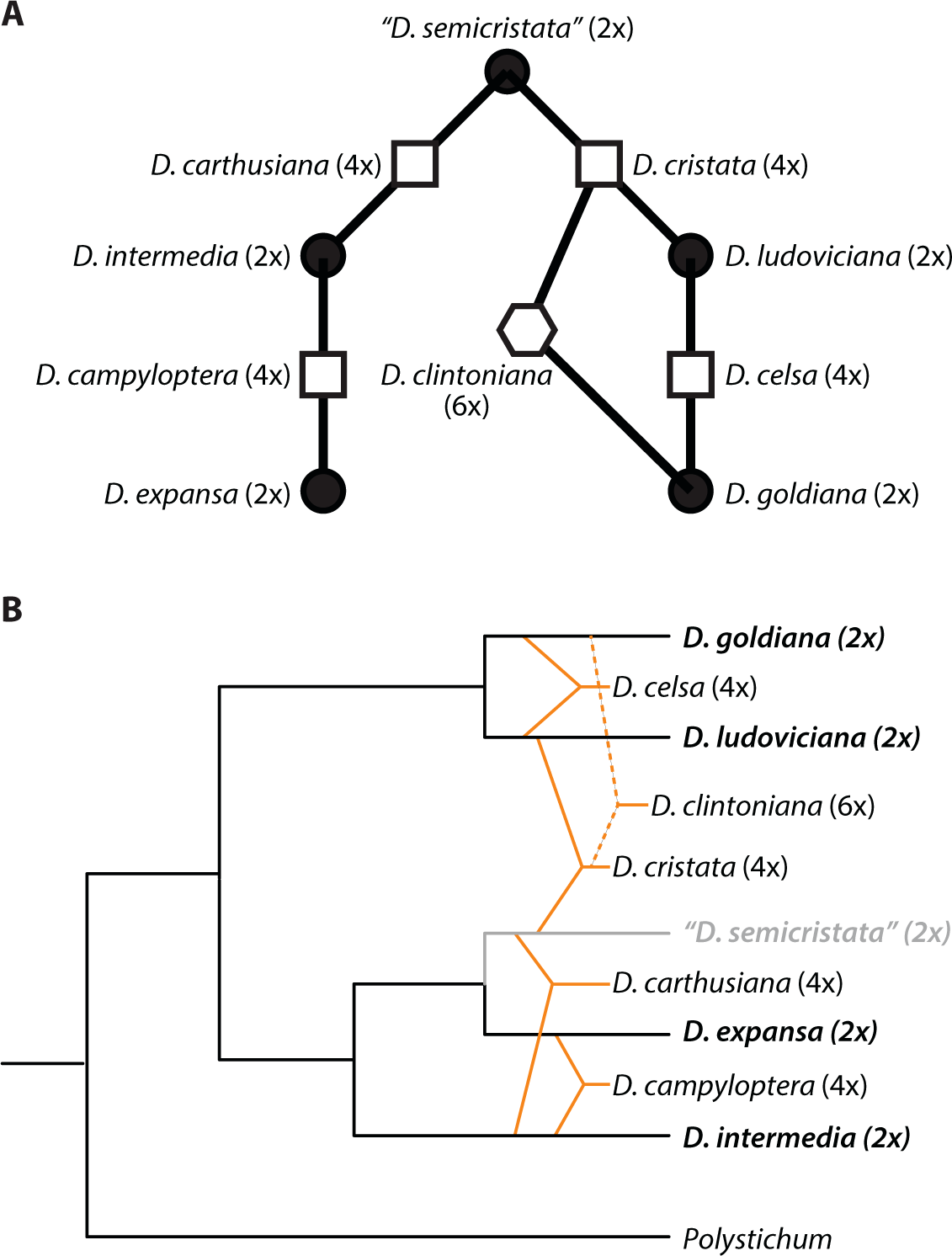
Hypothesized Relationships among North American *Dryopteris*. Synthesis of results from Sessa et al. 2012a and Sessa et al. 2012b. A) Links between shapes show the putative parents and their allopolyploid derivatives. Black circles are diploids, squares are tetraploids, and the hexagon is the one hexaploid species in the group. *Dryopteris semicristata* is presumed extinct. B) Placement of allopolyploids in the context of the backbone relationships among diploids. Tetraploids are indicated with solid orange lines and the hexaploid with dotted lines. The grey line denoting a sister relationship between *D. semicristata* and *D. expansa* reflects one possible placement for the extinct taxon based on previous analyses (Sessa et al., 2012b).

### Materials & Methods

### Testing the Effects of Phasing on Network Inference through Simulation

#### Simulating phased and unphased sequence data for an allopolyploid

We simulated gene trees and their nucleotide sequence data using the MSC model (Rannala and Yang 2003) with BPP v4.4.0 (Flouri et al. 2020) under a network with five tips, such that one taxon was a hybrid of the ancestors of two other tip taxa (Fig. 2). All simulations used the HKY85 (Hasegawa et al 1985) model of sequence evolution with a transition-transversion rate ratio of 3 and gamma-distributed among-site rate heterogeneity with a shape parameter of 0.6. Equilibrium frequencies were 0.3, 0.2, 0.3, and 0.2 for nucleotides A, C, G, and T. These parameters for the sequence evolution model were based on the observed means of parameter distributions across gene trees for our empirical *Dryopteris* data (see Materials and Methods: *Analyses of a Species Complex with Allopolyploidy*). The allopolyploid species *E* was treated as two lineages (*E* sister to *B* and *F* sister to *C*) whose alleles were pooled to form the hybrid species *E* at time τ_*h*_. This makes species *E* a tetraploid hybrid with the parents the ancestors of *B* and *C*. The mutation-scaled effective population size (θ) was constant among lineages and set at 0.01. Assuming a per-generation mutation rate (μ) of 1 × 10^–8^ and one year per generation yields an effective population size (*N_e_*) of 250,000 and hybridization age of 10 Ma (Fig. 2). We also simulated data such that θ was 0.001 (hybridization at 1 Ma) or 0.1 (hybridization at 100 Ma). Investigating such extreme values can generate general expectations for downstream estimation under a wide range of parameters, but it also has some biological salience as there is increasing evidence for reticulate evolution early in the diversification of plant orders and families (Sun et al. 2015; Stull et al. 2020; Liu et al. 2022).

**Figure 2.**
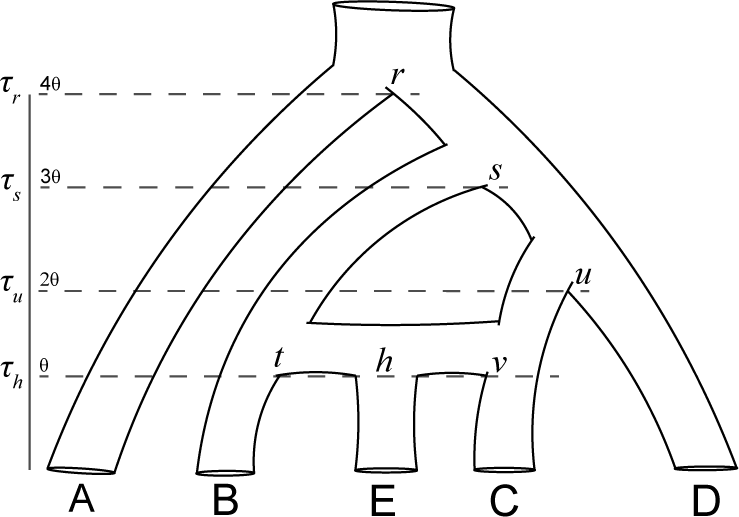
Species network used for simulation. The divergence times in expected substitutions per site are given for each node, and *h* is the hybrid node where two alleles enter from both *t* and *v*. E is an allotetraploid while other species are diploid. Absolute divergence times were changed by using values of θ at 0.001, 0.01, and 0.1. ILS was increased by multiplying θ by a factor of 2, 3, or 4 while keeping all τ_2_fixed to their initial values.

Because the distance between speciation nodes is 0.01 substitutions per site and θ = (Fig. 2), there are two coalescent units between nodes, 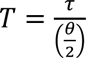 (Yang 2006), which yields a low probability (0.09) of gene tree discordance due to ILS (Hudson 1983). We incorporated more ILS into the simulation by keeping the node heights fixed but increasing θ to 0.02, 0.03, and 0.04. This increased the probability of a non-speciodentric split among all nodes to 0.25, 0.34, and 0.4, respectively. With three different starting values of θ and τ, this gave a total of 12 simulation conditions. While our simulations do not explore the limits of gene tree discordance, they allow us to learn about some general features of increasing ILS on network inference with different data types. We simulated 1000 gene trees and their sequences, 500 base pairs (bp) in length, under the MSC 100 times for each of the 12 conditions. We also explored the effects of sampling fewer genes on downstream analyses. For each replicate of 1000 gene trees and their sequences, we randomly sampled 400, 40, and four genes without replacement.

All simulations sampled haploid data. Two haploid sequences were sampled for each diploid species, and four haploid sequences were sampled for the allopolyploid species *E,* where two sequences came from each parental lineage. We then investigated unphased data in three ways. First, to generate unphased genotype sequences, the simulated haploid sequences for each species were collapsed into a single sequence in which heterozygous sites were represented by IUPAC ambiguity codes (genotype). The allopolyploid species was not restricted to only biallelic sites. Second, we generated haploid consensus sequences where for each variable site only one base was randomly chosen and retained (consensus). This could represent a case in which read coverage across a locus is highly uneven such that a haploid sequence is actually a chimera of two or more alleles. Finally, we simply picked one phased haploid sequence, which is possible when one parental haplotype has a majority of reads for a locus (pick one). This scenario where only one parent’s sequence would be recovered in the offspring could be anticipated in real data due to subgenome dominance (e.g., Buggs et al. 2014; Bird et al. 2018). In practice, we expect most *de novo* assemblers to generate output in between the haploid consensus data and pick one data.

#### Inferring species networks with phased and unphased simulated data

We estimated species networks with the SNaQ function (Solís-Lemus and Ané 2016) from PhyloNetworks v0.12.0 (Solís-Lemus et al. 2017) using Julia v.1.4.1 (Bezanson et al. 2015) from either the true gene trees used to simulate the data or gene trees estimated from the phased, pick one, genotype, and consensus sequence data. For estimated trees, we used IQTREE v1.6.10 (Nguyen et al. 2015) with the same substitution model used to simulate the data. Each SNaQ analysis used the species tree (A,((B,E),(C,D))) as the starting tree and allowed zero, one, or two reticulation events. Each analysis included ten independent optimizations of the pseudolikelihood score. We considered increasing numbers of reticulations if there was a pseudolikelihood score improvement of two or greater compared to the previous model with one fewer allowed hybrid edges. Note that pseudolikelihoods violate standard likelihood-based model selection criteria such as likelihood ratio tests or using AIC weights, and slope heuristics (Baudry et al. 2012) are often used instead. The choice of a cutoff here was necessary to streamline processing across thousands of simulations. We compared the estimated networks with one reticulation to the true network with the *hardwiredClusterDistance* function (Huson et al. 2010) in PhyloNetworks. This allowed us to score the number of replicates that 1) recovered the correct number of reticulations and 2) matched the true network when the number of reticulations was set to one. We estimated networks for samples of four, 40, 400, and 1000 gene trees for each of the 100 replicates from the 12 simulation conditions.

#### Effect of phasing on divergence time estimation

We also used our simulated phased and unphased data to estimate divergence times under the MSci model (Flouri et al. 2020) using BPP v4.4.0. Here, we estimate divergence times (τ), population sizes (θ), and the introgression probability, or proportion of immigrant loci in the receiving population, (φ) on the correct fixed species network. φ was originally represented by γ (Meng and Kubatko 2009) and referred to as an inheritance probability (Yu et al. 2011), but we use φ and the term introgression probability to be consistent with BPP (Flouri et al. 2020). Our MSci analyses used diffuse priors on τ_’_ and θ, with a mean on their simulated values and φ ∼ ϕ(1,1). Phased, pick one, and consensus sequences were treated as haploids while genotype sequences were treated as unphased diploids and used for analytical integration over phases (Gronau et al. 2011) implemented in BPP. Although this is not correct for the tetraploid, it is arguably more appropriate than treating all of the genotype sequences as haploids with ambiguous states. Each Markov chain Monte Carlo (MCMC) analysis collected 10,000 posterior samples, saving every 100 generations, while discarding the first 100,000 generations (i.e., 10% of the total run) as burnin. Scripts for simulation and subsequent analyses of simulated data are available in Dryad (https://doi.org/10.5061/dryad.5qfttdz53).

### A Phasing Pipeline for Polyploids

#### Target enrichment data

The success of empirical studies in which phasing was informative about hybridization or introgression events (e.g., Kamneva et al. 2017; Oberprieler et al. 2017; Eriksson et al. 2018) motivated us to use phased data to infer reticulate evolutionary histories of polyploids. We were aware of few instances of phasing genomic or phylogenomic data in polyploids, except in cases where chromosome-level whole-genome assemblies have characterized subgenomes in allopolyploid crops (Yang et al. 2017; Colle et al. 2019) or tree-based methods that are dependent on the sampling of parental lineages (Freyman et al. 2020; Nauheimer et al. 2020). We designed PATÉ for target enrichment data because of the availability of such data for many taxa, but it is applicable to other types of data with paired-end Illumina reads.

The end product of many *de novo* target enrichment assembly pipelines (such as HybPiper; Johnson et al. 2016) is a single consensus sequence for each locus for each individual. Allelic variation may be represented by ambiguous nucleotide codes within the single consensus sequence or lost when the pipeline outputs the haplotype consensus sequence where the majority vote from a collection of reads is used. We use these existing *de novo* assembly pipelines as a starting point to provide the reference sequence for each locus for each individual and leverage a recent phasing algorithm with high-quality variant calls to recover phased haplotype sequences for taxa with known ploidy levels. Ploidy levels are well established and relationships have previously been characterized for individuals in our *Dryopteris* analyses (Sessa 2012b).

#### Phasing alleles within loci

PATÉ (Supplementary Fig. S1) starts with assembled target enrichment loci, such as the supercontig files output from HybPiper (Johnson et al. 2016), that contain a single haplotype consensus sequence from each individual per locus. Reads for each individual are then realigned to their consensus locus using BWA v0.7.17 (Li and Durbin 2009). PCR duplicates are flagged with MarkDuplicates in Picard v2.9.2 (http://broadinstitute.github.io/picard), and variant calls are computed with HaplotypeCaller in GATK v.4.1.4 (McKenna et al. 2010). We applied the following hard filters with VariantFiltration in GATK: (1) the variant quality divided by the depth of the alternate allele (QD) should be two or greater; (2) a Fisher Exact test statistic based on a contingency table of read direction and whether that read has the reference or alternate allele (FS) should be 60 or less; (3) the root mean square of mapping quality scores (MQ) should be 40 or greater; (4) an approximation of the Rank Sum Test for variant position in a read should be negative eight or greater to avoid variants consistently towards the end of reads; (5) the fraction of reads with the alternate allele (AF) should be between 5% and 95%, and the total read depth of the site (DP) should be at least ten. The filter expressions are as follows: (1) QD < 2.0, (2) FS > 60.0, (3) MQ < 40.0, (4) ReadPosRankSum < −8.0, (5) AF < 0.05, AF > 0.95, (6) DP < 10. These loosely follow community recommendations on filters for germline variant discovery (DePristo et al. 2011). Notably, we do not perform quality score recalibration or filter on the mapping quality rank sum, as we anticipate allopolyploids could have a lower mapping quality associated with an alternate allele due to sequence divergence or structural variation among homeologous chromosomes. We also consider a very narrow window for filtering on allele frequency. Because increasing ploidy levels will generate smaller anticipated ratios of alternate to reference alleles, coupled with sequencing error and read stochasticity, we only aim to remove the most extreme cases. For example, if almost all reads support the alternate allele at a site, it is difficult to diagnose if the error lies in the consensus assembly or the read alignment. In these cases, only the reference site is retained, and the variant does not pass the allele frequency filter. However, the allele frequency filter could be removed if investigators are focused on organisms with higher ploidy levels. Additional options for pipeline configuration are available on GitHub (https://github.com/gtiley/Phasing).

Biallelic SNPs that pass filters are then phased with H-PoPG v.0.2.0 (Xie et al. 2016). H-PopG solves a heuristic phasing problem efficiently using dynamic programming. Although not guaranteed to be an optimal solution, H-PoPG has been shown to have high accuracy while also being fast (Xie et al. 2016; He et al. 2018; Moeinzadeh et al. 2020). Phasing variants in polyploids is difficult because for *n* variants and *k* ploidy, there are 2^*n*−1^)(*k* – 1)^*n*^ possible ways to link the sites together. H-PoPG evaluates possible solutions efficiently by grouping reads into *k* groups in such a way that differences within the groups are minimized. Focusing on target enrichment data also constrains the complexity of the phasing problem compared to whole-genome alignments because haplotype blocks are constrained to about 1000 bp. We then used the phased variants to create individual allele sequences, where invariable sites are filled in based on the reference sequence. In cases where there is no linkage information to phase across the entire locus, we retain the phasing only for the longest block. The variants for the shorter haplotype blocks can be collapsed into IUPAC ambiguity codes or treated as missing data based on the investigator’s preferences. PATÉ outputs analysis-ready fasta files with multiple alleles per species. Variants are only phased within loci; we do not attempt to assign loci to parental subgenomes. While this may complicate analyses of concatenated multi-locus datasets, it is ideal for the MSC that assumes free recombination between loci and can leverage multiple alleles per species for estimating θ. Assigning alleles to parental subgenomes should be possible with PATÉ outputs where both parental lineages are sampled and there is sufficient divergence for gene tree estimation.

### Analyses of a Species Complex with Allopolyploidy

#### The North American wood fern complex (Dryopteris)

We tested PATÉ using new target enrichment data from nine North American *Dryopteris* species, including both allotetraploid and allohexaploid taxa, with well-studied reticulate relationships (Sessa et al. 2012a, b), as well as two outgroup taxa from the sister genus *Polystichum*. All putative parental lineages are represented in our dataset, with the exception of a hypothesized extinct lineage (*D. semicristata*; Sessa et al. 2012b). We sampled two or three individuals for each *Dryopteris* taxon (Table S1). The target enrichment data were generated from the GoFlag 408 flagellate land plant probe set (Breinholt et al. 2021) at RAPID Genomics (Gainesville, FL). The target regions for this probe set are 408 exons found in 229 single or low-copy nuclear genes. We generated haplotype consensus assemblies for each with HybPiper (Johnson et al. 2016). The resulting supercontig sequences, which includes the exon region targeted by probes for enrichment and flanking intron regions, became our reference sequences for genotyping and phasing with PATÉ. We aligned both phased and haplotype consensus (i.e., the reference supercontig) sequences with MUSCLE v.3.8.31 with default settings (Edgar 2004). For comparison with simulation results, we generated a “pick one” dataset where only one phased haplotype per locus was randomly selected from each individual. A “genotype” dataset was generated by collapsing allelic variation into IUPAC codes for each individual as a locus.

#### Three species tests

We first explored the value of phasing data when estimating reticulate relationships among three species, where we sampled two extant parental lineages and their putative hybrid. We examined the three such extant relationships in the North American *Dryopteris* complex:

1. Two diploid parental lineages (*D. ludoviciana* and *D. goldiana*) and their putative allotetraploid descendent (*D. celsa*).
2. Two diploid parental lineages (*D. expansa* and *D. intermedia*) and their putative allotetraploid descendent (*D. campyloptera*).
3. One tetraploid parent (*D. cristata*), one diploid parent (*D. goldiana*), and their putative allohexaploid descendent (*D. clintoniana*).

These three scenarios represent different ages of parental divergence, with a common ancestor near 10 Ma for *D. ludoviciana* and *D. goldiana*, 15 Ma for *D. expansa* and *D. intermedia*, and 30 Ma for *D. cristata* and *D. goldiana* (Sessa et al. 2012b). We used both a full-likelihood Bayesian approach and a topology-based pseudolikelihood approach to estimate the correct species relationships from the different data types. First, using BPP v.4.4.0 (Flouri et al. 2020), we estimated log-marginal likelihoods (ln *mL*) with stepping-stone sampling (Xie et al. 2011) for the three possible rooted 3-taxon trees and network models that imply differences in the timing and direction of gene flow among species (Supplementary Fig. S2). Each ln *mL* estimate used 24 steps, and each step had a posterior sample of 10,000, saving every 30 generations after a 30,000 generation burnin (10% of the total run). The ln *mL* values were then used to calculate the six model probabilities following equation 1 using the bppr v.0.6.3 R package (https://github.com/dosreislab/bppr). A percentile-based 95% confidence interval was generated for each model probability by bootstrapping posterior distributions 1,000 times to investigate uncertainty in model selection.

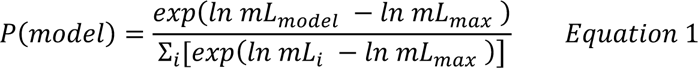

We performed four independent runs for each model for each data type prior to marginal likelihood estimation to evaluate mixing and diagnose potential convergence issues. The combined posteriors from the models that represent *a priori* hypotheses of relationships between allopolyploids and their parental lineages were used estimate all τ and θ for comparisons across data types. Notably, BPP can analytically integrate over possible phases of sequence data when given IUPAC codes representing heterozygous sites (Gronea et al. 2011), which is as accurate as perfectly phased data for diploids (Huang et al. 2022). In comparison, standard gene tree estimation under the Felsenstein likelihood (Felsenstein 1981) treat IUPAC codes as uncertainty and integrate over the possible tip states at sites independently.

For our topology-based method, we used the SNaQ function (Solís-Lemus and Ané 2016) from PhyloNetworks v0.12.0 (Solís-Lemus et al. 2017) to test the presence and placement of gene flow between the three species, while also including the two *Polystichum* outgroups. Gene trees were estimated for each data type across the three cases with IQ-TREE v1.6.10 (Nguyen et al. 2015), using ModelFinder (Kalyaanamoorthy et al. 2017) for model selection on the unpartitioned alignments. The starting tree was obtained with ASTRAL III v5.6.3 (Zhang et al. 2018) from the collection of ML gene trees. We tested the presence of zero, one, or two reticulations based on a two pseudolikelihood point cutoff for accepting an additional reticulation. Each analysis used ten independent optimizations. For SNaQ and ASTRAL analyses, all haplotypes were mapped to their respective species, as opposed to individuals.

#### Nine species tests

We investigated the differences in networks from SNaQ across data types for the nine-species complex. This complex involves multiple reticulation events on an edge, and thus the true network is not identifiable with topology-based methods (Solis-Lemus and Ané 2016). Although full-likelihood methods should not suffer from the some of the identifiability issues (Yu and Nakhleh 2015), they can impose a heavy computational burden (Wen et al. 2018; Zhang et al. 2018). Despite the inherent identifiability issues, it is worthwhile to examine the performance of scalable topology-based methods in such contexts to see if they will identify some of the correct reticulate relationships and if they also recover erroneous ones. Thus, we performed analyses of the nine-species complex and two outgroups across the four data types with SNaQ as described above, but we allowed up to six reticulation events. We used ASTRAL III v5.6.3 (Zhang et al. 2018) to generate the starting species tree for network estimation using gene trees inferred from IQ-TREE v1.6.10 (Nguyen et al. 2015) with the best model selected by ModelFinder (Kalyaanamoorthy et al. 2017). 100 standard bootstrap replicates were performed for each gene tree, which were used to evaluate gene tree incongruence and informativeness with respect to the ASTRAL species tree from phased data for the consensus, genotype, and pick one gene trees with phyparts (Smith et al. 2015). To assess node support for SNaQ, we performed 100 non-parametric bootstrap replicates, in which both loci and bootstrap trees per locus were sampled with replacement. Each bootstrap replicate was distributed independently to hasten the computation using a Perl script that accommodates multiple alleles available on GitHub (https://github.com/gtiley/bootSNaQ-MA).

## Results

### Simulation Shows Benefits of Phased Data

#### Network inference

In our simulation results, all data types performed well estimating the correct number of reticulations (Supplementary Fig. S3). The analyses typically converged to the true number of reticulations with 40 or more gene trees, except when using gene trees estimated from genotype data with a 25% (θ = 2 × τ_*h*_) chance of a discordant gene tree. However, using phased data provided more accurate estimates of the placement of the reticulation edge in comparison to the genotype, consensus, and pick one data (Fig. 3). When the true gene trees were used, which have information about the allopolyploid’s hybridization event (i.e., the allele sequences are sister to their respective parents in every tree), the correct network can be inferred with 40 loci regardless of the age of hybridization or degree of ILS. The gene trees estimated from phased data perform equally well, aside from requiring a few more gene trees for recent hybridization events with high ILS (θ = 4 × τ_*h*_). Analyses based on genotype, consensus, and pick one data almost always recover the true network when sampling 400 or more loci, but can miss the true network over half the time with 40 or fewer loci. However, the genotype data failed to converge to the true network with increasing loci for a hybridization age of 10 Ma with low to moderate amounts of ILS. Otherwise, varying degrees of ILS had a negligible effect on network estimation among data types, and estimated gene trees, aside from those from phased data, were less efficient when the hybridization age was more recent.

**Figure 3.**
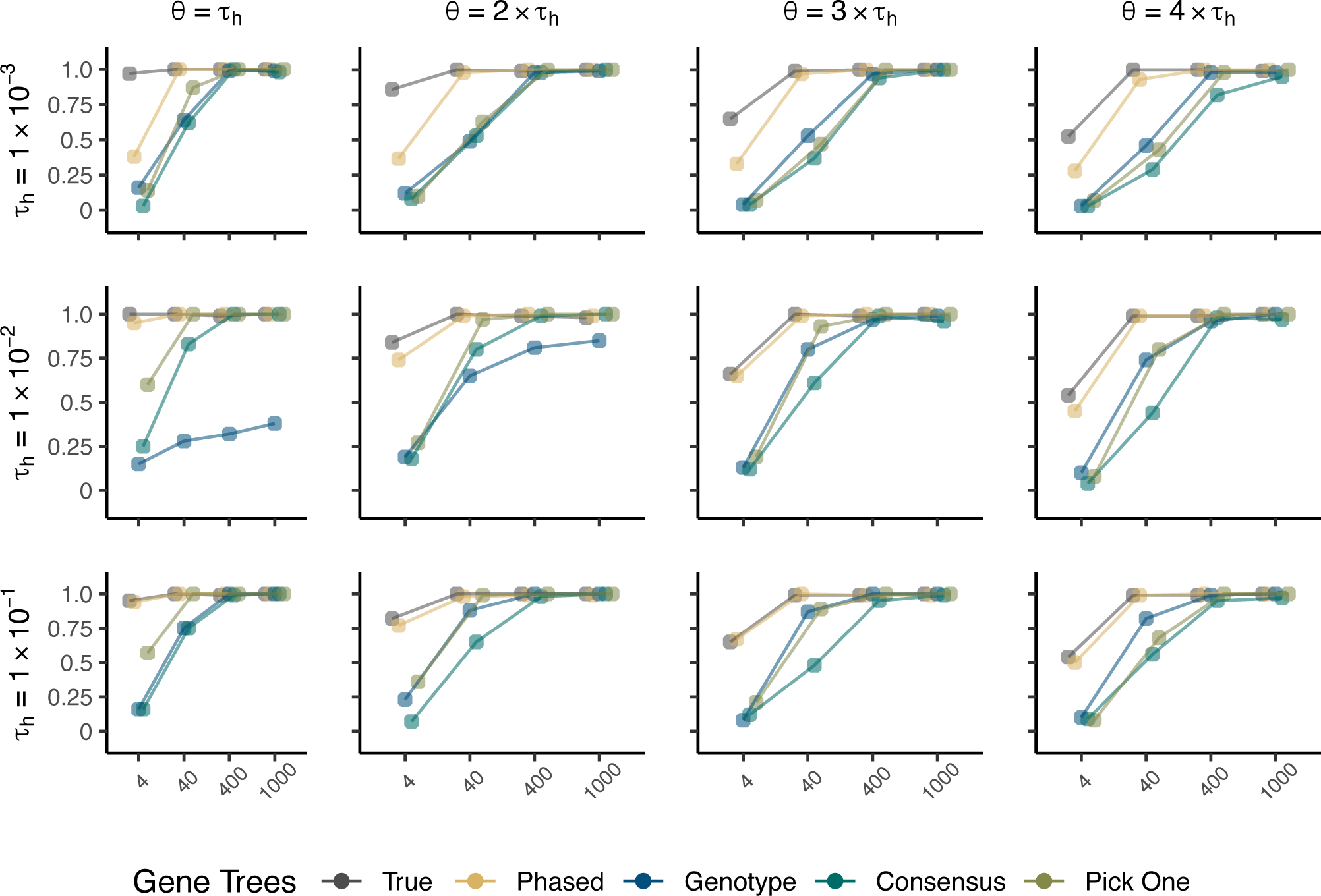
Proportion of simulation replicates that correctly identify the allopolyploid lineage using the SNaQ function of PhyloNetworks. The x-axes are the number of loci sampled for each simulation. The y-axes are the proportions of correct networks. Results are based on networks estimated with a single reticulation, even if that network was considered less optimal than networks with zero or two reticulations. We saved the true gene trees from the simulations, while we estimated gene trees with the phased, genotype, consensus, and pick one data types.

#### Divergence times

Using phased data, whether using all alleles or only one, typically resulted in accurate estimation of the hybridization age (Fig. 4). When divergence was low, there was a small amount of estimation error, such that the phased data underestimated and the pick one data overestimated the hybridization age, but the pattern suggests estimates could converge to the true values with more loci. Interestingly, convergence towards the true value appeared more efficient with higher ILS (Fig. 4). The phased and pick one data could otherwise accurately estimate the hybridization time with 40 or fewer loci across all other divergence and ILS scenarios. For analyses with genotype and consensus data, as the number of loci increased and uncertainty in the posterior was reduced, the posterior mean did not converge to the true estimate, and the simulated value was not within the highest posterior density (HPD) interval. There was always a small degree of estimation error with the genotype data despite using analytical phasing, and the consensus data overestimated the hybridization age between 1.5x and 2x when using 400 loci across all simulation scenarios. These biases from genotype or consensus data were also propagated for the parent nodes of the hybridization event across all levels of ILS for τ_*h*_ = 0.001 (Supplementary Figs. S4-S7), τ_*h*_ = 0.01 (Supplementary Figs. S8-S11), and τ_*h*_= 0.1 (Supplementary Figs. S12-S15). Nodes from speciation events could always be estimated accurately with high precision by the phased and pick one data types provided enough loci were sampled. The phased data was more efficient, always converging to the true value by 40 loci, while the pick one data required 400 to recover τ_2_ when the sequence divergence was low at τ_*h*_ = 0.001. There was a small degree of estimation error among speciation nodes for the genotype and consensus data, and the patterns suggested they might not converge to the true values with more loci. However, speciation times were overestimated by genotype and consensus data for τ_*h*_= 0.001 (Supplementary Figs. S4-S7) and τ_*h*_= 0.01 (Supplementary Figs. S8-S11), but underestimated by the genotype data and overestimated by the consensus data for τ_*h*_= 0.1 (Supplementary Figs. S12-S15). Across all simulations, there was little effect of increasing ILS aside from increasing HPD intervals, but the uncertainty was negligible when 400 loci were sampled.

**Figure 4.**
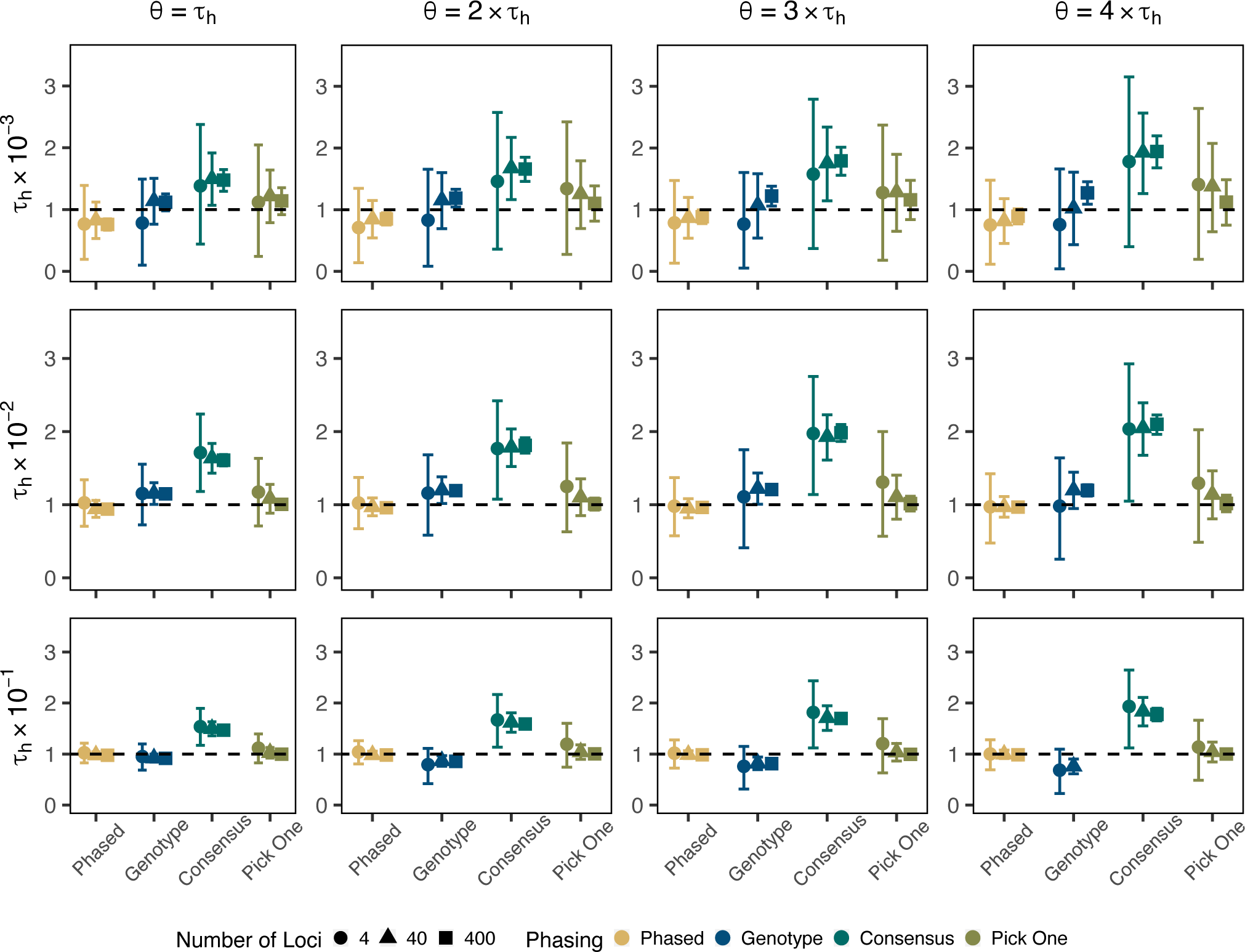
Estimating the Timing of Hybridization with the MSci Model. Divergence times are for node *h* in Figure 2. Divergence times are measured as the expected number of substitutions per site. The dashed line represents the true simulated values. Points are posterior means, and error bars are 95% HPD intervals, averaged across 100 replicates. Results are not shown for the genotype data when τ_*h*_= 0.1 and θ = 4τ_*h*_ because posterior means went towards infinity and numerically unstable values that cannot be displayed.

### Analyses of target enrichment data from Dryopteris

#### Recovery of Phased Loci

On average, 62% of loci sequenced for an individual were phased with eight variants passing filters per locus per individual (Supplementary Table S2). The ploidy level appears to be strongly associated with the number of phased loci. Among diploids, only 32% of loci were phased; the other 68% of diploid loci were either homozygous or had too few linked variants for phasing. For tetraploids and hexaploids, 87% and 94% of loci, respectively, were phased such that two or more phased haplotype sequences could be recovered. Among loci where phasing was possible, variants were almost always resolved as a single haplotype block, as opposed to being split into two or more blocks because not enough reads were available to physically link variants. For polyploids, the number of unique phased haplotype sequences most frequently matched the ploidy level, except in the case of a single *D. campyloptera* individual (B087-D08), which also had relatively few recovered loci. Phasing data only extended sequence alignment length by about two base pairs relative to the haplotype consensus sequences (Supplementary Table S3). However, phased data more than tripled the number of parsimony informative sites per alignment, similar to the increase in number of tips (Supplementary Table S3).

#### Placing a single reticulation event

For our three-species full-likelihood analyses with the MSci model, all data types decisively supported the putative reticulation hypothesis in the cases of the allotetraploids *D. celsa* or *D. campyloptera* (Table 1). However, only the genotyped data, coupled with the analytical phasing technique of BPP, chose the putatively correct model for the allohexaploid *D. clintoniana* (Table 1). The phased and pick one data types selected the network with the diploid *D. goldiana* as the hybrid descendent of the allohexaploid *D. clintoniana* and allotetraploid *D. cristata*, while the best model for consensus data was a binary tree with *D. clintoniana* sister to *D. cristata* (Table 1). However, model probability confidence intervals suggest a great deal of uncertainty underlies model selection results for the allohexaploid case, with the upper end of the interval for the putatively correct model at 0.603, 0.582, and 0.996 for the phased, consensus, and pick one data, respectively (Table 1). Model parameters typically converged across all analyses with the exception of the phased data for the *D. clintoniana* triplet (Supplementary Fig. S16), which was due to one MCMC run despite convergence of the lnL values.

**Table 1.** Model Probabilities for Possible Topological Hypotheses Using the MSci model.

| Hybrid | Model | Phased Model Probability | Genotype Model Probability | Consensus Model Probability | Pick One Model Probability |
| --- | --- | --- | --- | --- | --- |
| *D. celsa* | 1 | $8.02 \times 10^{-142}$ ($3.62 \times 10^{-146}$, $3.31 \times 10^{-137}$) | $1.95 \times 10^{-173}$ ($1.03 \times 10^{-177}$, $2.78 \times 10^{-169}$) | $5.79 \times 10^{-168}$ ($3.15 \times 10^{-172}$, $8.7 \times 10^{-164}$) | $4.32 \times 10^{-105}$ ($1.99 \times 10^{-109}$, $2.27 \times 10^{-101}$) |
| | 2 | $3.18 \times 10^{-126}$ ($4.89 \times 10^{-131}$, $1.34 \times 10^{-121}$) | $5.44 \times 10^{-136}$ ($2.17 \times 10^{-140}$, $7.99 \times 10^{-132}$) | $1.66 \times 10^{-113}$ ($9.18 \times 10^{-118}$, $2.83 \times 10^{-109}$) | $2.78 \times 10^{-84}$ ($1.24 \times 10^{-88}$, $2.14 \times 10^{-80}$) |
| | 3 | $3.33 \times 10^{-124}$ ($5.19 \times 10^{-129}$, $1.66 \times 10^{-119}$) | $5.77 \times 10^{-149}$ ($1.63 \times 10^{-153}$, $1.31 \times 10^{-144}$) | $5.66 \times 10^{-162}$ ($6.09 \times 10^{-166}$, $9.31 \times 10^{-158}$) | $1.44 \times 10^{-109}$ ($6.87 \times 10^{-114}$, $2.06 \times 10^{-105}$) |
| | 4 | $1.36 \times 10^{-146}$ ($5.71 \times 10^{-151}$, $5.14 \times 10^{-142}$) | $6.98 \times 10^{-149}$ ($2.24 \times 10^{-153}$, $1.62 \times 10^{-144}$) | $2.89 \times 10^{-152}$ ($1.19 \times 10^{-156}$, $4.33 \times 10^{-148}$) | $1.44 \times 10^{-109}$ ($8.54 \times 10^{-114}$, $8.43 \times 10^{-106}$) |
|  | **5** | **1 (1, 1)** | **1 (1, 1)** | **1 (1, 1)** | **1 (1, 1)** |
| | 6 | $3.69 \times 10^{-113}$ ($1.31 \times 10^{-117}$, $1.45 \times 10^{-108}$) | $1.59 \times 10^{-143}$ ($7.07 \times 10^{-148}$, $3.22 \times 10^{-139}$) | $1.97 \times 10^{-118}$ ($1.83 \times 10^{-122}$, $4.43 \times 10^{-114}$) | $1.46 \times 10^{-92}$ ($1.12 \times 10^{-96}$, $8.06 \times 10^{-89}$) |
| *D. campyloptera* | 1 | $1.42 \times 10^{-251}$ ($1.6 \times 10^{-256}$, $5.32 \times 10^{-247}$) | $3.77 \times 10^{-251}$ ($2.11 \times 10^{-255}$, $6.16 \times 10^{-247}$) | $3.15 \times 10^{-154}$ ($3.78 \times 10^{-158}$, $5.25 \times 10^{-150}$) | $1.52 \times 10^{-130}$ ($5.8 \times 10^{-135}$, $3.11 \times 10^{-126}$) |
| | 2 | $1.03 \times 10^{-210}$ ($1.12 \times 10^{-215}$, $8.01 \times 10^{-206}$) | $2.54 \times 10^{-226}$ ($1.33 \times 10^{-230}$, $2.41 \times 10^{-222}$) | $1.69 \times 10^{-110}$ ($9.98 \times 10^{-115}$, $1.72 \times 10^{-106}$) | $1.11 \times 10^{-90}$ ($5.02 \times 10^{-95}$, $2.23 \times 10^{-86}$) |
| | 3 | $2.16 \times 10^{-242}$ ($2.64 \times 10^{-247}$, $6.06 \times 10^{-238}$) | $1.53 \times 10^{-184}$ ($7.18 \times 10^{-189}$, $5.03 \times 10^{-180}$) | $1.24 \times 10^{-129}$ ($6.49 \times 10^{-134}$, $2.39 \times 10^{-125}$) | $5.72 \times 10^{-123}$ ($5.18 \times 10^{-127}$, $1.1 \times 10^{-118}$) |
| | 4 | $1.22 \times 10^{-133}$ ($7.33 \times 10^{-139}$, $7.63 \times 10^{-129}$) | $1.87 \times 10^{-193}$ ($6.47 \times 10^{-198}$, $2.12 \times 10^{-189}$) | $2.8 \times 10^{-134}$ ($1.98 \times 10^{-138}$, $7.92 \times 10^{-130}$) | $4.19 \times 10^{-114}$ ($2.78 \times 10^{-118}$, $6.67 \times 10^{-110}$) |
|  | **5** | **1 (1, 1)** | **1 (1, 1)** | **1 (1, 1)** | **1 (1, 1)** |
| | 6 | $4.96 \times 10^{-173}$ ($6.57 \times 10^{-178}$, $3.7 \times 10^{-168}$) | $1.88 \times 10^{-189}$ ($9.15 \times 10^{-194}$, $1.93 \times 10^{-185}$) | $5.11 \times 10^{-103}$ ($2.49 \times 10^{-107}$, $4.81 \times 10^{-99}$) | $5.44 \times 10^{-86}$ ($3.65 \times 10^{-90}$, $1.32 \times 10^{-81}$) |
| *D. clintoniana* | 1 | $3.44 \times 10^{-110}$ ($5.27 \times 10^{-115}$, $1.25 \times 10^{-105}$) | $6.01 \times 10^{-65}$ ($2.21 \times 10^{-69}$, $1.99 \times 10^{-60}$) | $2.62 \times 10^{-12}$ ($8.24 \times 10^{-17}$, $1.82 \times 10^{-8}$) | $1.52 \times 10^{-13}$ ($3.47 \times 10^{-18}$, $4.54 \times 10^{-10}$) |
| | 2 | $1.44 \times 10^{-04}$ ($1.46 \times 10^{-9}$, $0.826$) | $1.91 \times 10^{-47}$ ($4.69 \times 10^{-52}$, $3 \times 10^{-43}$) | $4.49 \times 10^{-9}$ ($1.83 \times 10^{-13}$, $4.18 \times 10^{-5}$) | $5.84 \times 10^{-6}$ ($9.84 \times 10^{-11}$, $7.03 \times 10^{-2}$) |
| | 3 | $4.6 \times 10^{-134}$ ($6.97 \times 10^{-139}$, $5.65 \times 10^{-130}$) | $2.31 \times 10^{-55}$ ($5.51 \times 10^{-60}$, $5.35 \times 10^{-51}$) | **1 (4.84 × 10<sup>-2</sup>, 1)** | $2.23 \times 10^{-12}$ ($5.82 \times 10^{-17}$, $1.38 \times 10^{-8}$) |
| | 4 | $1.23 \times 10^{-115}$ ($3.09 \times 10^{-120}$, $2.69 \times 10^{-111}$) | $2.18 \times 10^{-27}$ ($3.04 \times 10^{-32}$, $3.68 \times 10^{-23}$) | $1.88 \times 10^{-4}$ ($1.05 \times 10^{-8}$, $0.728$) | $2.02 \times 10^{-27}$ ($3.5 \times 10^{-32}$, $1.34 \times 10^{-23}$) |
| | **5** | **4.23 × 10<sup>-5</sup> (7.74 × 10<sup>-10</sup>, 0.603)** | **1 (1, 1)** | $1.73 \times 10^{-4}$ ($6.85 \times 10^{-09}$, $0.582$) | $5.7 \times 10^{-3}$ ($1.31 \times 10^{-7}$, $0.996$) |
| | 6 | **1 (3.81 × 10<sup>-2</sup>, 1)** | $2.48 \times 10^{-31}$ ($6.11 \times 10^{-36}$, $5.76 \times 10^{-27}$) | $1.04 \times 10^{-15}$ ($6.71 \times 10^{-20}$, $6.87 \times 10^{-12}$) | **0.994 (3.6 × 10<sup>-3</sup>, 1)** |

Note: Topologies of the six models that enumerate the possible binary and reticulate evolutionary histories are displayed in Figure S2. The best model for each data type and triplet is shown in bold. A percentile-based 95% confidence interval is given for each probability by bootstrapping the posteriors.

The simulation experiments established some expectations that the phased and pick one data would provide unbiased estimates of divergence times for nodes involved in the hybridization events, while the genotyped and consensus data would provide under- and overestimates, respectively. However, phased data generally produced younger mean τ estimates for the *D. celsa* triplet with reduced credible intervals relative to other data types, but much older mean τ estimates with more uncertainty for the *D. campyloptera* and *D. clintoniana* triplets (Fig. 5; Supplementary Table S4). Similarly, there was inconsistency in θ estimates observed among data types across triplets, with notably larger credible intervals for phased data in the *D. celsa* and *D. clintoniana* triplets. Estimates of φ were sensitive to data type across triplets, such that the mean φ suggested the major contribution to the focal allopolyploid from phased data was *D. goldiana* into *D. celsa*, *D. intermedia* into *D. campyloptera*, and *D. goldiana* into *D. clintoniana*, with other data types sometimes disagreeing (Fig. 5). Credible intervals around φ always spanned φ = 0.5, but estimates for genotype, consensus, and phased data suggested some biased gene retention for the *D. celsa* and *D. clintoniana* triplets (Supplementary Table S4).

**Figure 5.**
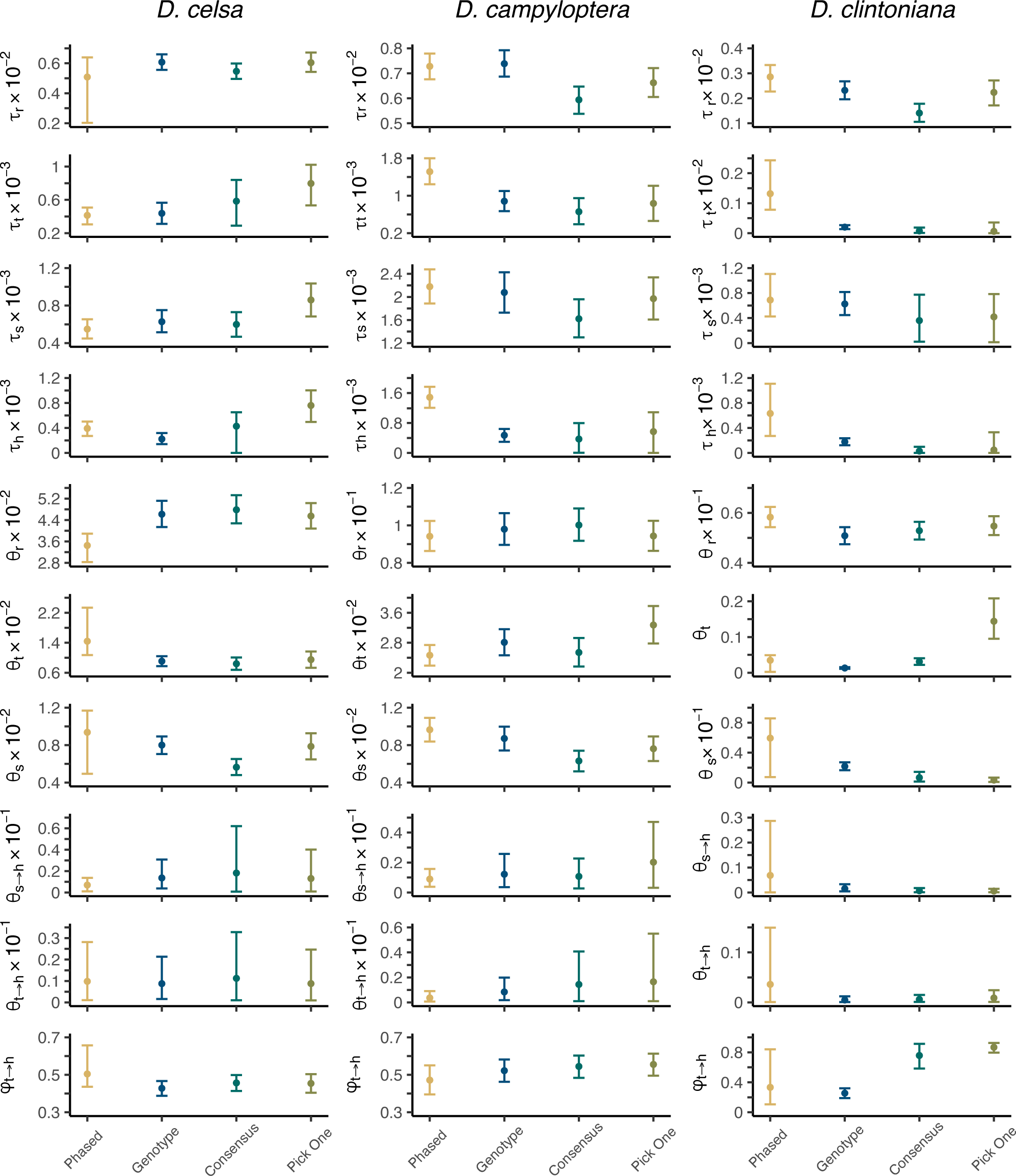
*Dryopteris* Divergence Times, Population Sizes, and Introgression Probabilities under the MSci Model. Points are posterior means and error bars show the 95% HPD intervals. Divergence times are measured as the expected number of substitutions per site.

When performing a network search based on gene tree distributions with SNaQ, all data types recovered the putatively correct relationship among *D. celsa* and its diploid parental lineages (Fig. 6a). For *D. campyloptera*, all data types correctly detected one hybrid edge (Supplementary Table S5), but they got the direction of gene flow wrong (Fig. 6b). All data types also detected one hybrid edge for the allohexaploid *D. clintoniana* (Supplementary Table S5), but there was disagreement over the underlying network. Both the phased and consensus data invoked a ghost lineage sister to the three sampled species, such that gene flow was from the ghost into the allotetraploid *D. cristata*. While not unreasonable given the putatively extinct *D. semiscristata* and missing *D. ludoviciana* from the sample, the relationship between *D. clintoniana* and its parental lineages was undetected (Fig. 6c). The genotype and pick one data recovered the correct network, but reversed the direction of gene flow from allohexaploid *D. clintoniana* into allotetraploid *D. cristata* (Fig 6c).

**Figure 6.**
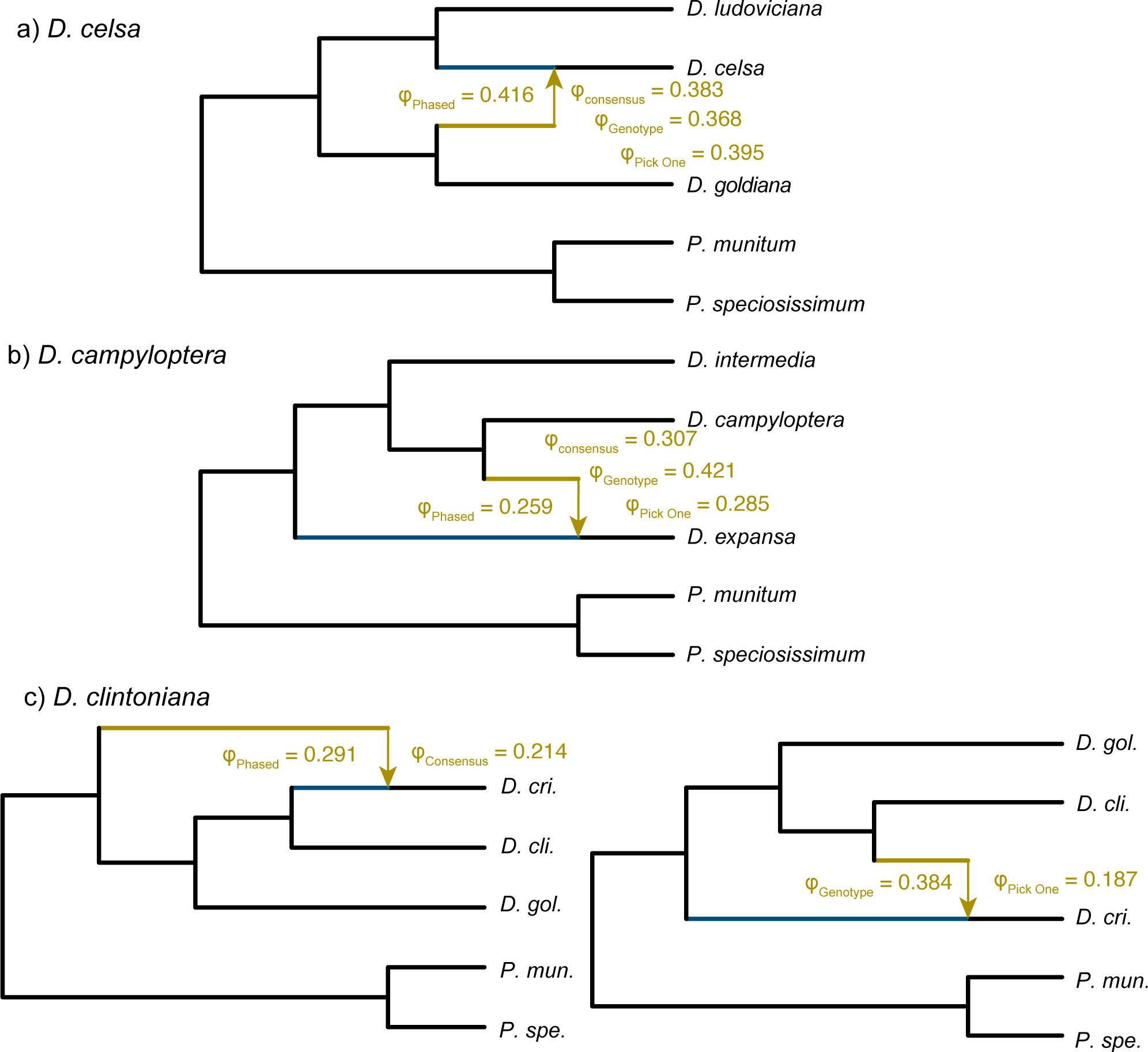
Network search results for three-taxon *Dryopteris* cases with SNaQ. Colors indicate the major and minor edge. The arrow from the minor edge shows the direction of introgression. The introgression probability (φ) is shown for each data type. In cases where a) *D. celsa* or b) *D. campyloptera* were the allotetraploids under investigation, all data types recovered the same species network. When c) *D. clintoniana* was the allohexaploid lineage from the allotetraploid *D. cristata* and diploid *D. goldiana* progenitors, different networks were recovered by the phased or consensus versus genotype or pick one data types.

#### Inferring relationships among a complex with multiple reticulation events

The nine *Dryopteris* species SNaQ network analysis with phased data identified two out of five hypothesized reticulation events correctly along with one incorrect reticulation edge (Fig. 7a; Supplementary Table S6). The analysis correctly identified the diploids *D. ludoviciana* and *D. goldiana* as parents to the allotetraploid *D. celsa*, as well as the diploids *D. expansa* and *D. intermedia* as parents to the allotetraploid *D. campyloptera*. A hybrid edge from the common ancestor of *D. celsa*, *D. ludoviciana*, and *D. goldiana* to the common ancestor of *Polystichum* was also recovered with an introgression probability of 0.335, although with low bootstrap support. The parents of *D. celsa* were also detected by the other data types, but all had some degree of error. The genotype data found a reticulation edge between the allotetraploid *D. carthusiana* and diploid parent *D. intermedia*, but reversed the direction of gene flow (Fig. 7b). A reticulation edge was also detected between *D. cristata* and the common ancestor of *D. clintoniana*, *D. goldiana*, *D. celsa*, and *D. ludoviciana*, which is not completely unreasonable given the allotetraploid *D. cristata* is believed to be a parent of *D. clintoniana*. The consensus data only detected two reticulation edges (Supplementary Table S6), with the second edge correctly showing gene flow from the diploid parent *D. intermedia* into the allotetraploid *D. carthusiana*, provided the other parent *D. semicristata* is presumed extinct obscuring the network interpretation (Fig. 7c). The pick one data, similar to the genotype data, inferred a reticulation edge between *D. carthusiana* and *D. intermedia* with the wrong direction of gene flow (Fig. 7d). The pick one data also detected a reticulation edge from *D. cristata* into the common ancestor of *D. carthusiana D. intermedia*, *D. campyloptera*, and *D. expansa*, which lacks a clear interpretation other than invoking missing lineages underlying complexity of the backbone topology (Fig. 1). In the one case where all data types detected the correct relationship regarding *D. celsa*, the introgression probability was approximately 0.5 with phased data while between 0.37 and 0.39 with others (Fig. 7; Supplementary Fig. S17).

**Figure 7.**
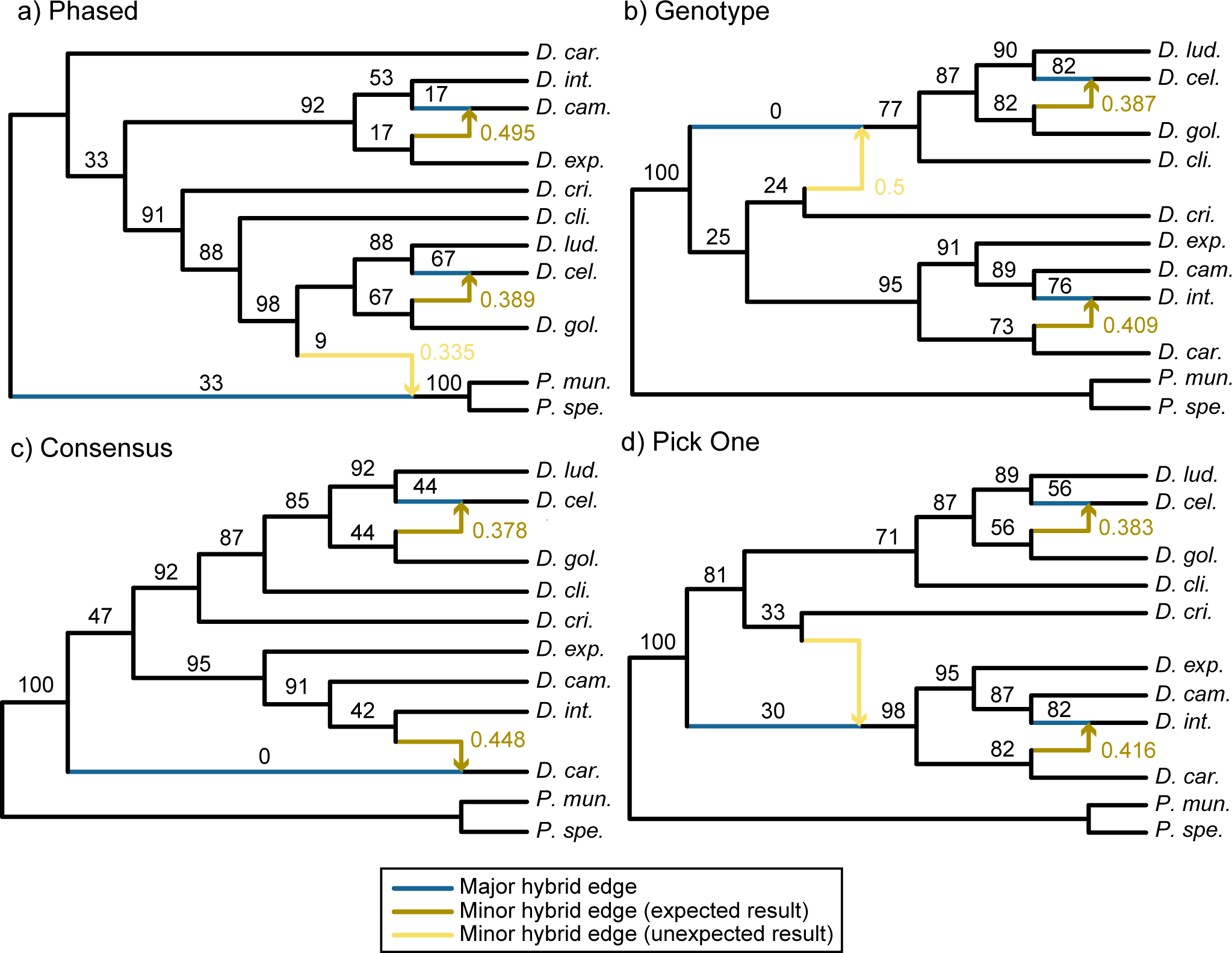
Networks for all North American *Dryopteris* inferred from SNaQ. Colors indicate major and minor (reticulation) edges, and which minor edges were hypothesized or an unexpected result. Introgression probabilities for each minor edge are provided. Numbers on branches subtending nodes are bootstrap support values. Results are shown separately for a) phased data where all haplotype sequences are included in gene trees, b) genotyped data with a single sequence for individual with IUPAC ambiguity codes, c) haplotype consensus sequences, and d) a single haplotype sequence representing each individual among gene trees.

Differences in the network estimates among data types may be explained by their underlying spectra of quartet concordance factors (Supplementary Fig. S18). While spectra were overall similar among data types, some deviations from expectations under ILS alone were stronger in the phased data versus others (Supplementary Fig. S18). Analyses of gene tree incongruence among the consensus (Supplementary Fig. S19), genotype (Supplementary Fig. S20), and pick one (Supplementary Fig. S21) data showed similar levels of conflict among the different data types with respect to a species tree estimated from the phased data (such analyses with multiple alleles per species is not possible). The frequency of conflicting splits with high bootstrap support was similar across datatypes for all nodes (Supplementary Fig. S22), and low-support conflicts represented on average 56% of splits within *Dryopteris*.

## Discussion

New phylogenetic network methods offer the promise of elucidating the often complex reticulate histories of polyploid lineages, even in the presence of ILS. Our results demonstrate that under some conditions phasing polyploid target enrichment data can improve the accuracy of such network inferences as well as divergence time estimates for networks, and we describe a novel pipeline (PATÉ) to address the difficult problem of phasing polyploid data. Although PATÉ could handle different types of genomic data, such as transcriptomes and whole genomes, target enrichment data are ideal for computational phasing because they can yield high (i.e. greater than 100x) and even coverage within a locus. This was demonstrated in our *Dryopteris* analyses when almost all loci were recovered as single haplotype blocks (Supplementary Table S2). Because MSC methods treat loci independently and assume free recombination between loci, it is not necessary to assign individual loci to parental subgenomes for network analyses. However, the allele sequences output by PATÉ can also be used as input for other tools such as *Homologizer* (Freyman et al. 2023) or *HybPhaser* (Nauheimer et al. 2020), which attempt to phase across loci and assign parental subgenomes. Our methods also enable population genomic studies of polyploids (Tiley et al. 2024) where accurate estimation of site frequency spectra can be used for demographic analyses otherwise complicated by polyploidy (e.g., Excoffier et al. 2013; Liu and Fu 2020) or SNP-based network inference in the absence of variation suitable for gene tree estimation (Blischak et al. 2018a; Olave and Meyer 2020).

### Promises and Limits of Phasing

The prospect of using alleles from phased genomic data presents an exciting step towards revealing the evolutionary history of polyploids, which remains a critical impediment within the plant evolution community (McKain et al. 2018). New strategies for explicitly addressing this challenge are emerging (Freyman et al. 2023; Nauheimer et al. 2020; Mendez-Reneau et al. 2023), and PATÉ can be a useful tool by phasing variants for many individuals while leveraging genotype quality information. Our simulations demonstrated that phasing can improve estimates of reticulate evolutionary relationships using network methods. In simulations, phased data with all alleles represented in gene trees could more efficiently recover the placement and directionality of hybrid edges than the genotyped, haplotype consensus, or single haplotype sequences (Fig. 3). The advantages of using phased data for network estimation diminished in simulations as the number of loci increased (Fig. 3), and this was evident from our empirical analyses of *Dryopteris* triplets, which produced similar results across data types with 330 or 328 loci (Table 1; Fig. 6), except for the correct inference of allopolyploid and parent relationships with genotyped data in the case of *D. clintoniana* (Table 1).

Both our simulations (Fig. 4; Supplementary Figs. S4-S15) and previous investigations (Anderman et al. 2019) suggested phased data can be important for unbiased divergence time estimation, especially estimating the origin of a hybrid lineage (Fig. 4). Our empirical analyses also demonstrated how the timing of introgression τ_*h*_and other parameters can be greatly affected by the data type used (Fig. 5). In our *Dryopteris* analyses, τ_*h*_ for *D. celsa* was at least in the same order-of-magnitude among data types, which had partially overlapping credible intervals, while phased τ_*h*_ estimate *for D. campyloptera* was nearly four times older than the other data types (Fig. 5; Supplementary Table S4). This was surprising given that simulations suggested haplotype consensus data should generate older estimates than phased data (Fig. 4). The *D. clintoniana* τ_*h*_estimate from phased data was also much older than other data types, albeit with a wide 95% HPD interval. There were noticeable mixing problems in our *D. clintoniana* full-likelihood analyses, which makes it difficult to evaluate whether the uncertainty was due to extreme differences in θ among phased alleles within *D. clintoniana* and/or the computational difficulty of the Bayesian methods with increasing sequence data. In all cases, the differences between τ_*h*_among data types is not consistent with results from the simulation experiments, which suggested the consensus data should overestimate age compared to phased or pick one data, with genotyped data having slightly younger estimates than phased data. This highlights the difficulties of simulating data that capture the complexities of empirical data and makes deciding which estimate is more reliable difficult. For example, the uncertainty of τ_*h*_ with phased data for *D. celsa* and *D. campyloptera* triplets was reduced compared to the consensus and pick one estimates, but greater uncertainty was observed for τ and θ at other nodes (Fig. 5). A potential compromise from simulations and empirical analyses is the use of analytical phasing of polyploid data, as done with our genotyped data for BPP analyses. Analytical phasing (Gronau et al. 2011; Flouri et al. 2020) has been shown to perform well in diploids (Huang et al. 2022), and any potential biases from ignoring ploidy seem outweighed by getting the *D. clintoniana* model correct (Table 1) as well as seemingly reasonable parameter estimates under the MSci model with tight credible intervals (Fig. 5).

Our analyses also suggest that phasing may be problematic when the parental lineages are deeply diverged. Although the phased data were able to accurately estimate the age of older speciation nodes in simulations (Supplementary Figs. S4-S15), as phylogenetic information is lost from multiple hits, the influence of the prior in full likelihood analyses should become stronger. Deeper divergences will likely generate more switch errors among phased haplotype sequences generated by PATÉ (Xie et al. 2016), as there will be more opportunities for paralogy, structural variants, and sequence divergence to confound the read mapping, genotyping, and phasing processes. When enough reads are available to call high-confidence variants, we suggest that phasing with PATÉ can improve network estimation based on the correct identification of the allotetraploids *D, celsa* and *D. campyloptera* and their parental lineages in the analyses of the North American *Dryopteris* complex (Fig. 7a); however, phasing alone cannot overcome some inherent challenges of network estimation.

### Challenges of Network Estimation

Our analyses highlight the difficulty of estimating the evolutionary histories of reticulate complexes, regardless of data type. The full-likelihood implementation of the MSci model appears to be useful when the number of tips and plausible models for investigation are low, but these methods may not be practical for generating hypotheses and exploring unknown relationships for large numbers of taxa (e.g., Zhang et al. 2018). Quartet-based methods are much faster and accurate when model assumptions are met (Solis-Lemus and Ané 2016) and show similar accuracy to full-likelihood methods for estimating introgression probabilities (Flouri et al. 2020). However, there are scenarios where the true networks are non-identifiable (Yu and Nakhleh 2015; Solís-Lemus and Ané 2016). For example, when multiple introgression events affect the same lineage, such that there are overlapping cycles in a network (Solís-Lemus and Ané 2016), the expected quartet distribution under the MSci model becomes a poor fit for the empirical data (Cai and Ané 2020), and the network estimated may be incorrect. These effects were evident in our empirical analyses where the relationship between *D. campyloptera*, *D. expansa*, and *D. intermedia* was missing in the nine-species analysis (Fig. 7), which we expect is due to a reticulation edge present between *D. intermedia* and *D. carthusiana*. Phasing sequence data adds information that can improve estimates (Supplementary Table S3), but unsampled or extinct lineages, such as the hypothesized *D. semicristata,* can create significant barriers to recovering the true evolutionary history of reticulate complexes, regardless of how many loci or individuals are available. Another identifiability problem is getting the direction of gene flow correct (Yu and Nakhleh 2015; Thawornwattana et al. 2023), and these scenarios were evident in our quartet-based analyses of a triplet that reversed gene flow from the allotetraploid *D. campyloptera* into the diploid *D. expansa* (Fig. 6d) or in the analyses of the full complex that reversed gene flow from the allotetraploid *D. carthusiana* into the diploid *D. intermedia* (Fig. 7b and 7d).

### Insights into Dryopteris Evolution

The North American *Dryopteris* complex has been well-characterized through the study of multi-locus nuclear and chloroplast phylogenies, morphology, cytological observations, chromatography, and isozyme analyses (reviewed in Sessa et al. 2012b). This makes it a useful system for testing our phasing pipeline, and our analyses add nuance to our understanding of some of the relationships among *Dryopteris* species. For example, our results with phased data indicated *D. campyloptera* has received more loci from *D. intermedia* than *D. expansa* (Fig. 7a), and *D. intermedia* is likely the maternal progenitor of *D. campyloptera* (Sessa et al. 2012b). Following the allopolyploidy event, the *D. intermedia* subgenome may have been dominant, providing a selective advantage for *D. campyloptera* in its distribution at the time (Bird et al. 2018). The phased nine-taxon analyses also supported the hypothesis of the unsampled diploid lineage *D. semicristata*, based on the placement of *D. carthusiana* as sister to the rest of *Dryopteris*, and the general sensitivity of *D. carthusiana* and *D. cristata* in the phylogenetic backbone across data types (Fig. 7). Additional insights may be gained by a more complete sampling of the genus, as the North American *Dryopteris* complex is not a clade (Sessa et al. 2012b), which should alleviate some of the identifiability problems. Analyses of the North American *Dryopteris* complex alone highlight the need for caution when interpreting networks, which are increasingly prominent plant evolutionary research.

### Does Phasing Matter?

Combining phased data with recent network methods can help confront a major challenge of plant phylogenetics: resolving the complex histories of polyploids. While our analyses showed haplotype consensus sequences can be adequate for resolving single reticulation events where both parental lineages are sampled, phased sequences may provide more information when only a few loci are available for cases where sequence divergence is not too high, while more generally improving divergence time estimates (see also Huang et al. 2022). Analytical phasing approaches (Gronau et al. 2011; Flouri et al. 2020) provide a way forward for full-likelihood methods while avoiding the potential for switch errors from read-based phasing as done in PATÉ, but these will become difficult with many tips. This highlights the need for continued development of models that explicitly consider polyploidization mechanisms (Jones et al. 2013) while addressing scalability and limiting assumptions for contemporary genomic data (Yan et al. 2022). Some reticulate complexes will be difficult to disentangle with any data when confronted with statistical identifiability problems, such as networks with multiple reticulations on a single edge (Solís-Lemus et al. 2016) when using quartet-based methods, and our *Dryopteris* analyses demonstrate that caution is needed when interpreting biological processes from phylogenetic networks in the absence of corroborating evidence. As haplotype-phased genomes from long-read data become increasingly available, the discussion of phase resolution will become trivial, but progress on models that leverage such data and improve interpretation of processes, especially in polyploids that maintain distinct subgenomes, from networks is still needed.

## Data Availability

PATÉ is available through GitHub (https://github.com/gtiley/Phasing) and can be run on any UNIX environment after installing basic genotyping software and H-PoPG. Simulated and empirical data supporting findings and files for replicating some analyses are available from the Dryad Digital Repository: https://doi.org/10.5061/dryad.5qfttdz53. Raw Fastq reads for *Dryopteris* individuals are available through the NCBI SRA database and are associated with BioProject PRJNA725004. Individual SRA Identifiers are available in Supplementary Table S1.

## Supplementary Material

Data and supplementary material is available from the Dryad Digital Repository: https://doi.org/10.5061/dryad.5qfttdz53

## Acknowledgements

The authors thank A.M. Duffy, K. Imwattana, M. Nieto-Lugilde, B.T. Piatkowski, and A.J. Shaw for helpful discussions and providing feedback on the manuscript. We also thank M.G Johnson, J. Mendez Reneau, L. Nauheimer, and C.J. Rothfels for discussions and sharing their strategies for phasing sequence data. We are grateful to the people who made data collection possible; we used *Dryopteris* samples collected by C.J. Rothfels, M.A. Sundue, and W. Testo, and DNA extractions were performed by S.B. Carey and E. Lockwood. The manuscript was improved through helpful comments from the associate editor S.D. Smith and three anonymous reviewers.

