## Supplementary Information for "Benefits and Limits of Phasing Alleles for Network Inference of Allopolyploid Complexes"

### Supplementary Figures

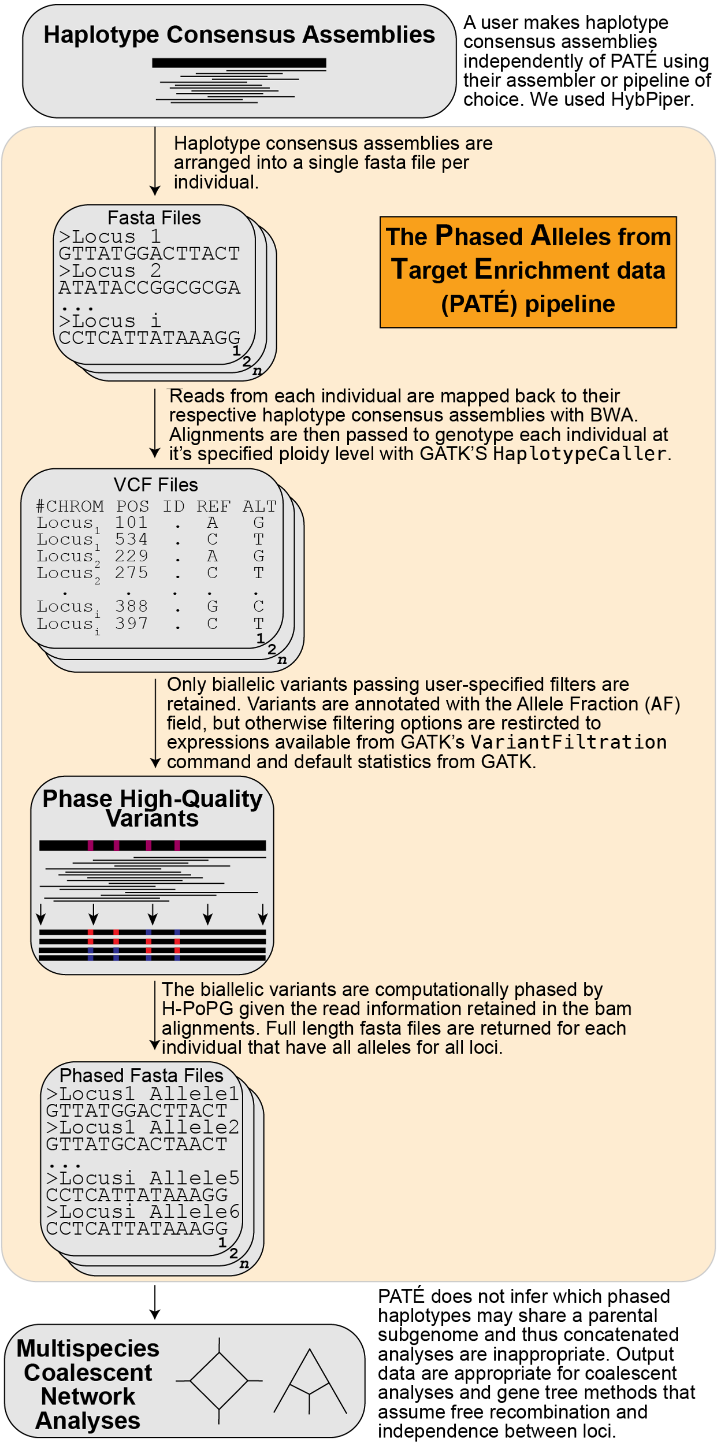


**Figure S1 — Generalized PATÉ workflow.** PATÉ is focused on calling variants and recovering phased haplotype sequences for arbitrary ploidy levels.

**Key to triples**

***D. campyloptera***
A = *D. campyloptera*
B = *D. expansa*
C = *D. intermedia*

***D. celsa***
A = *D. celsa*
B = *D. goldiana*
C = *D. ludoviciana*

***D. clintoniana***
A = *D. clintoniana*
B = *D. cristata*
C = *D. goldiana*

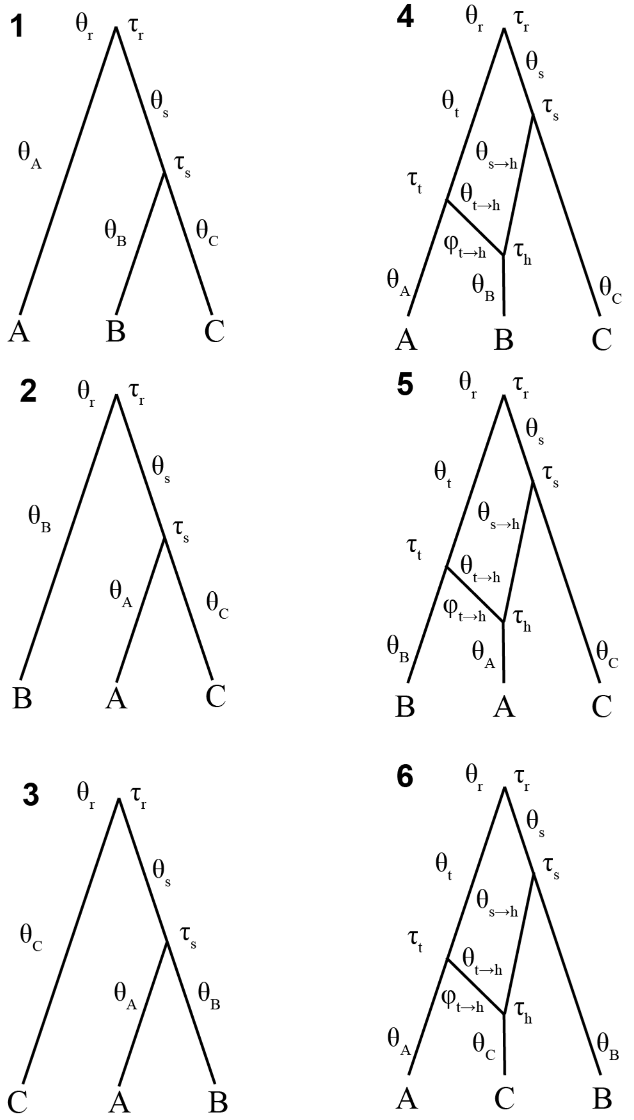


**Figure S2 — Network hypotheses for three-species analyses with marginal likelihoods.** Numbers indicate model numbers from main text figures and tables. Model five is the correct one based on available evidence in *Dryopteris*. Only one $\varphi$ parameter is estimated because the other branch is constrained to $1 - \varphi$.

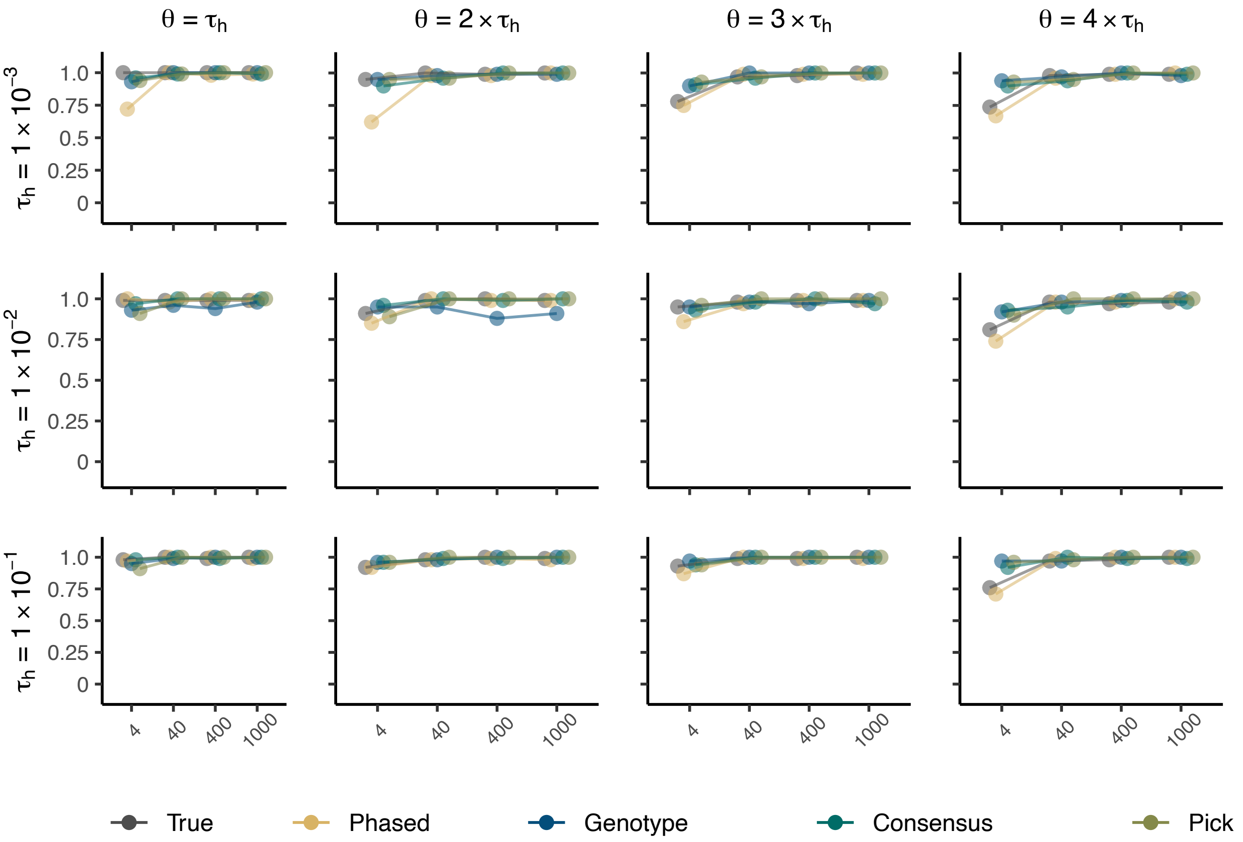


**Figure S3 — Proportion of simulations that correctly detect one reticulation.** The x-axis is the number of loci and the y-axis is the proportion of correct networks. Networks are scored as correct if the best network is determined to have one reticulation based on at least a two-unit pseudologlikelihood increase over no reticulations and no more than a two-unit pseudologlikelihood improvement for including two reticulations.

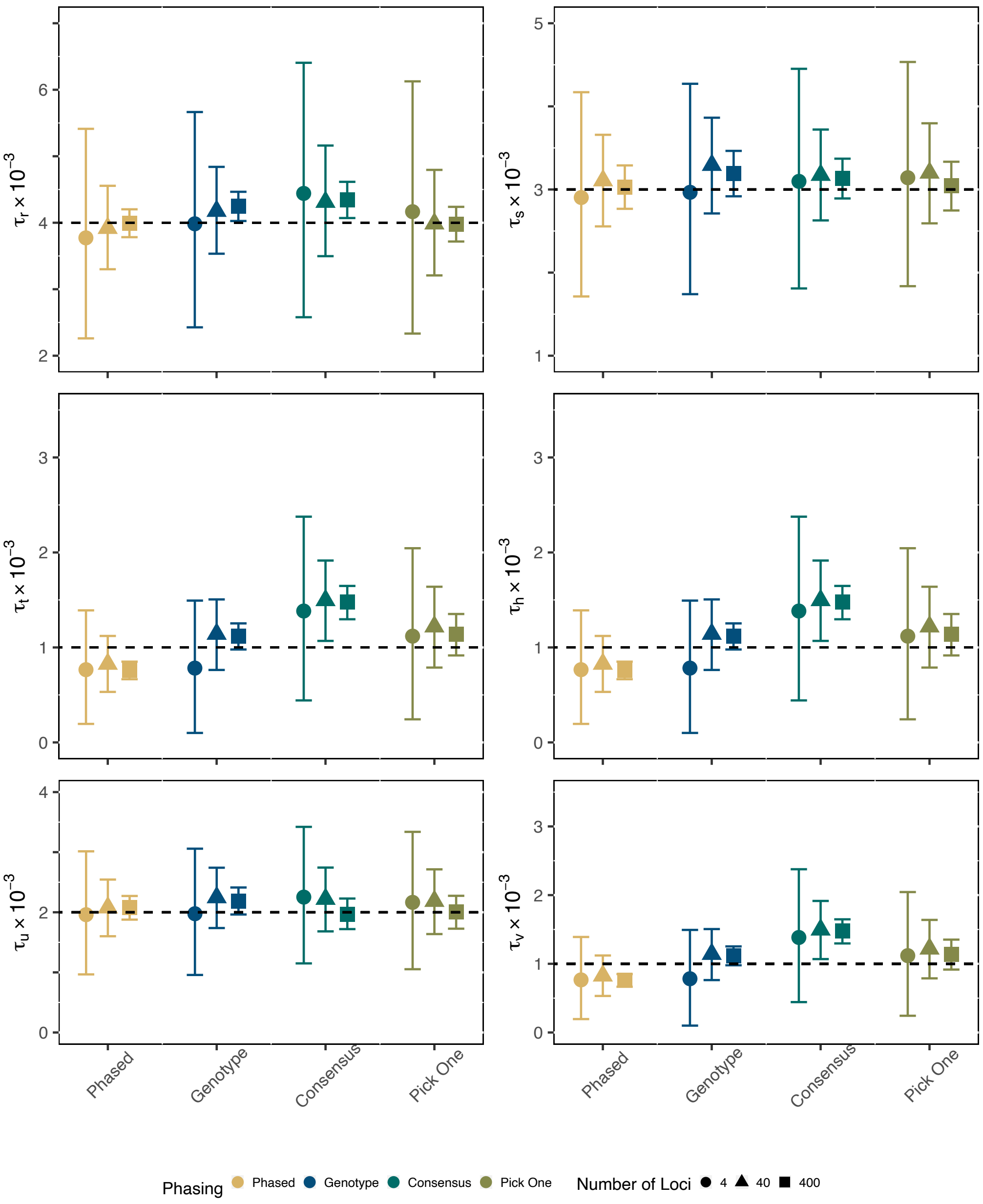


**Figure S4 — Divergence Time Estimation for Species Network with $\tau_h = 0.001$ and $\theta = \tau_h$.** The y-axis is divergence time for each individual parameter. Divergence times are measured in the expected number of substitutions per site. The dashed line represents the true simulated values. Points are posterior means and error bars are 95% HPD intervals, averaged across 100 replicates.

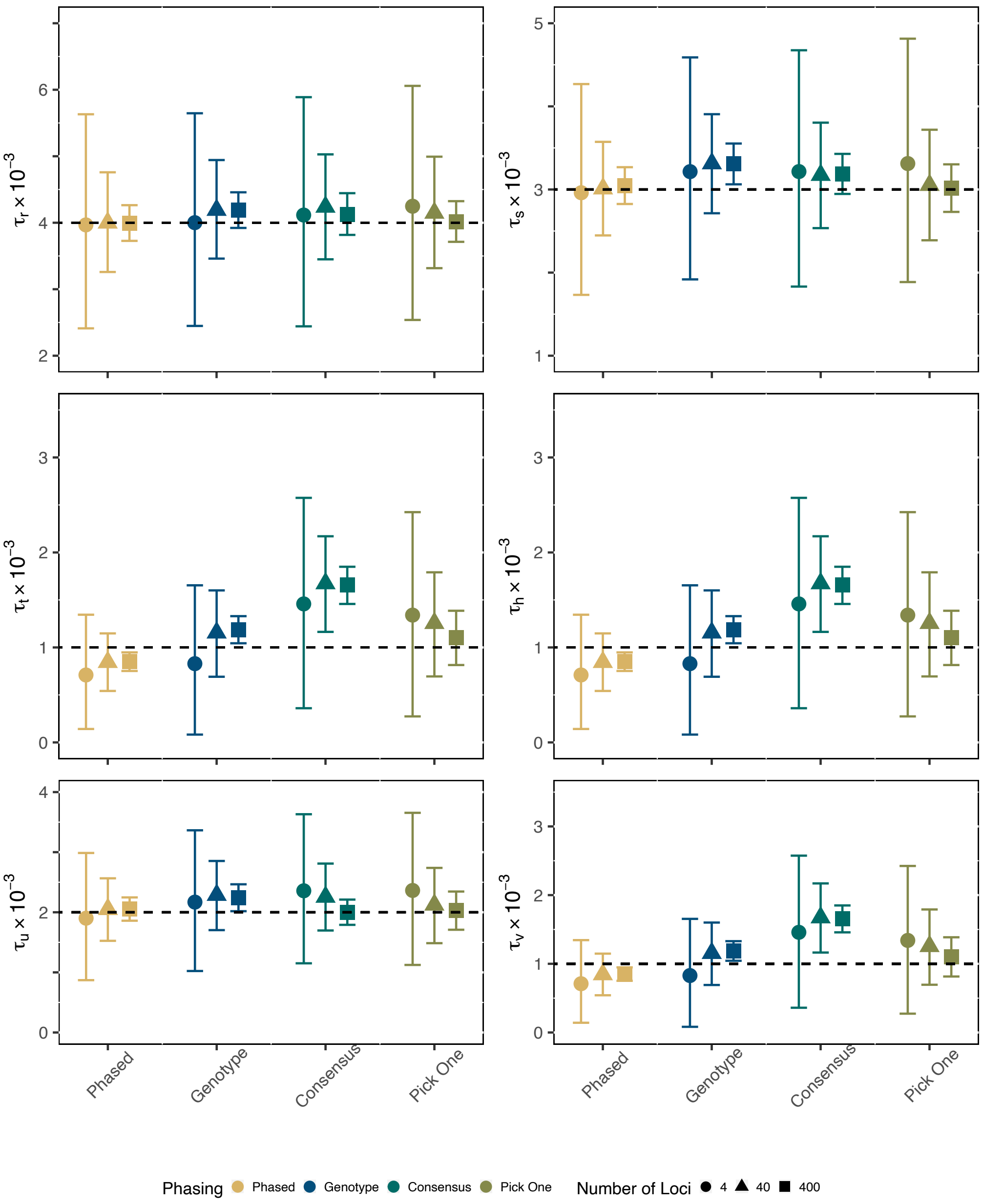


**Figure S5 — Divergence Time Estimation for Species Network with $\tau_h = 0.001$ and $\theta = 2\tau_h$.** The y-axis is divergence time for each individual parameter. Divergence times are measured in the expected number of substitutions per site. The dashed line represents the true simulated values. Points are posterior means and error bars are 95% HPD intervals, averaged across 100 replicates.

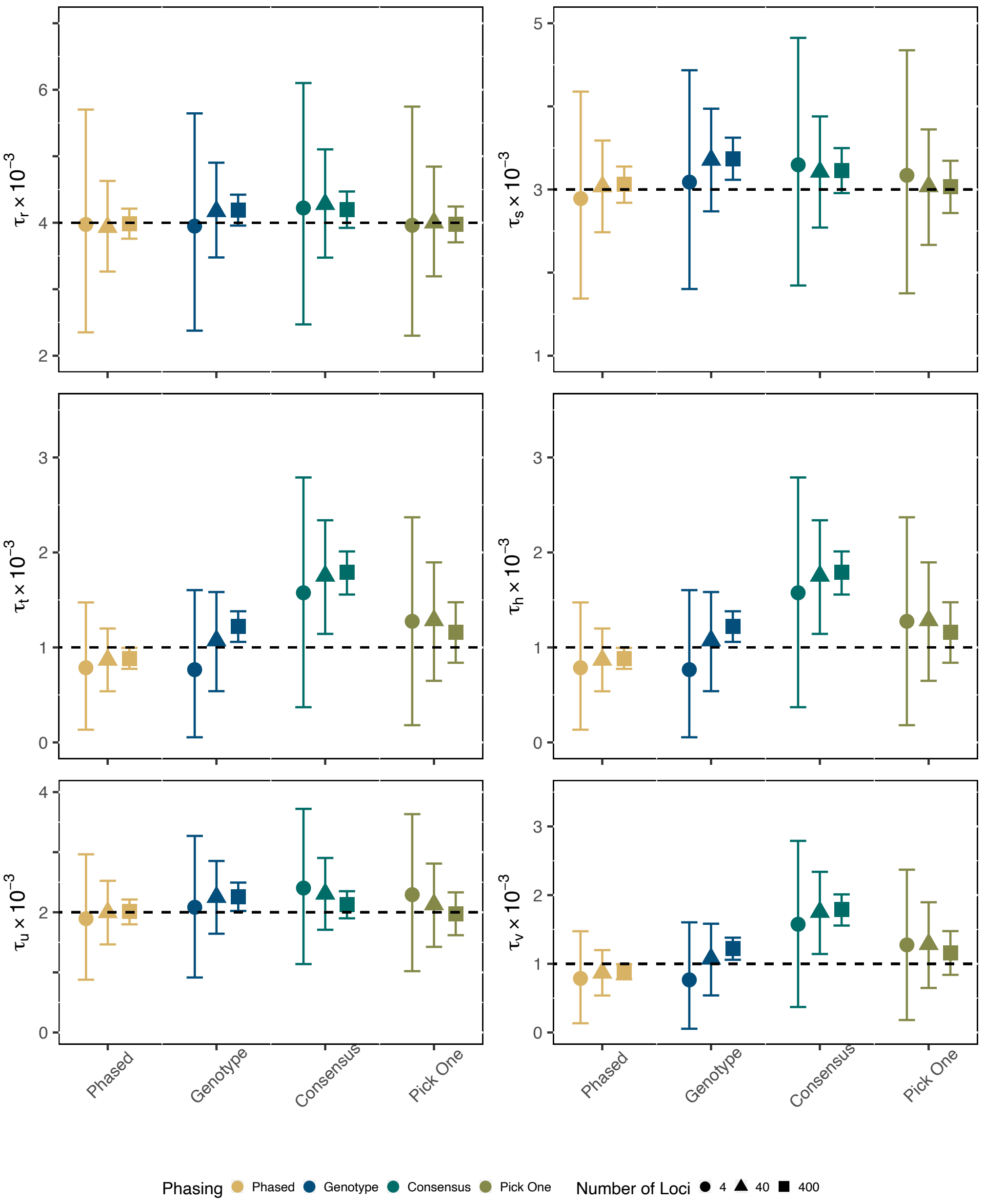


**Figure S6 — Divergence Time Estimation for Species Network with $\tau_h = 0.001$ and $\theta = 3\tau_h$.** The y-axis is divergence time for each individual parameter. Divergence times are measured in the expected number of substitutions per site. The dashed line represents the true simulated values. Points are posterior means and error bars are 95% HPD intervals, averaged across 100 replicates.

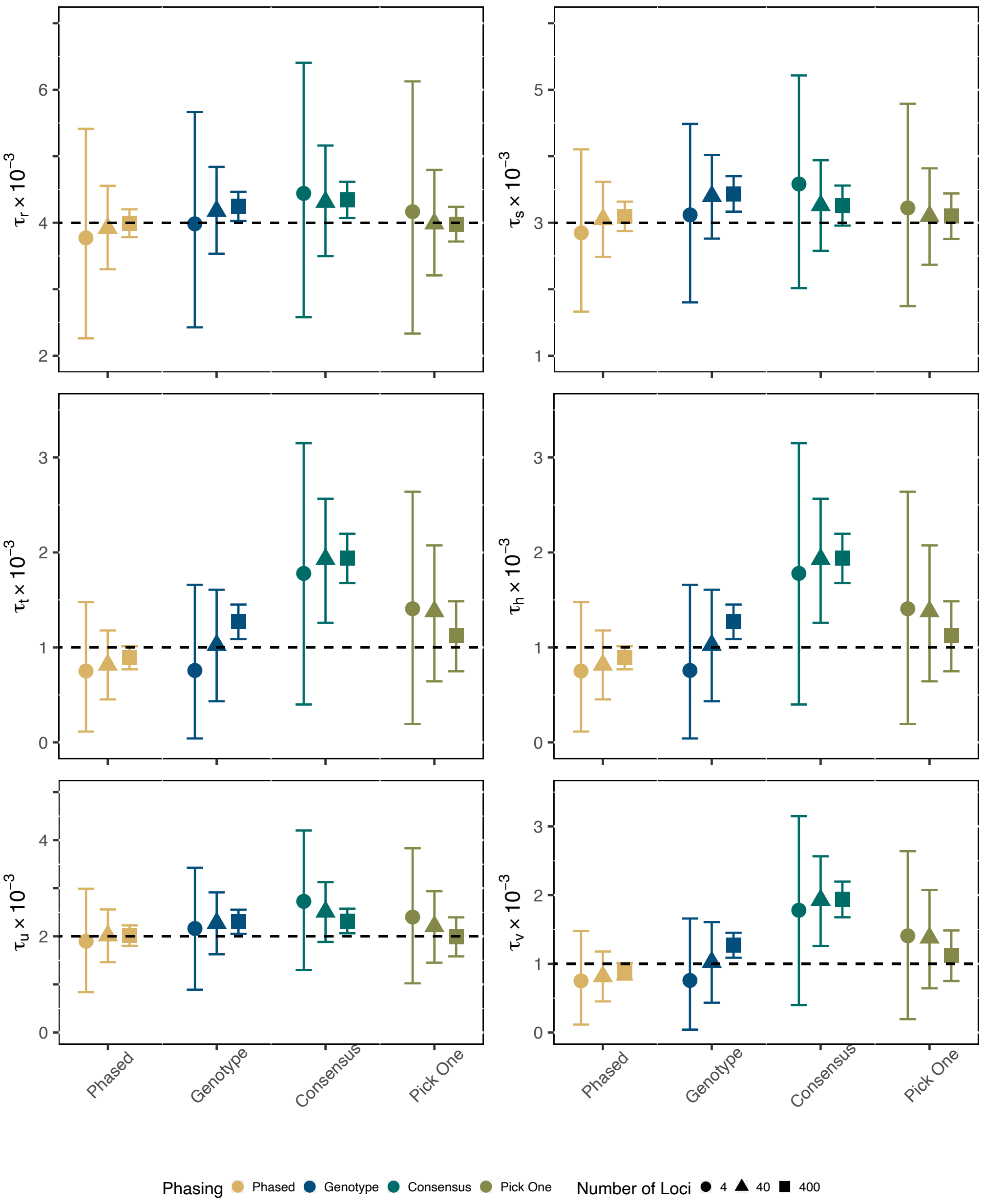


**Figure S7 — Divergence Time Estimation for Species Network with $\tau_h = 0.001$ and $\theta = 4\tau_h$.** The y-axis is divergence time for each individual parameter. Divergence times are measured in the expected number of substitutions per site. The dashed line represents the true simulated values. Points are posterior means and error bars are 95% HPD intervals, averaged across 100 replicates.

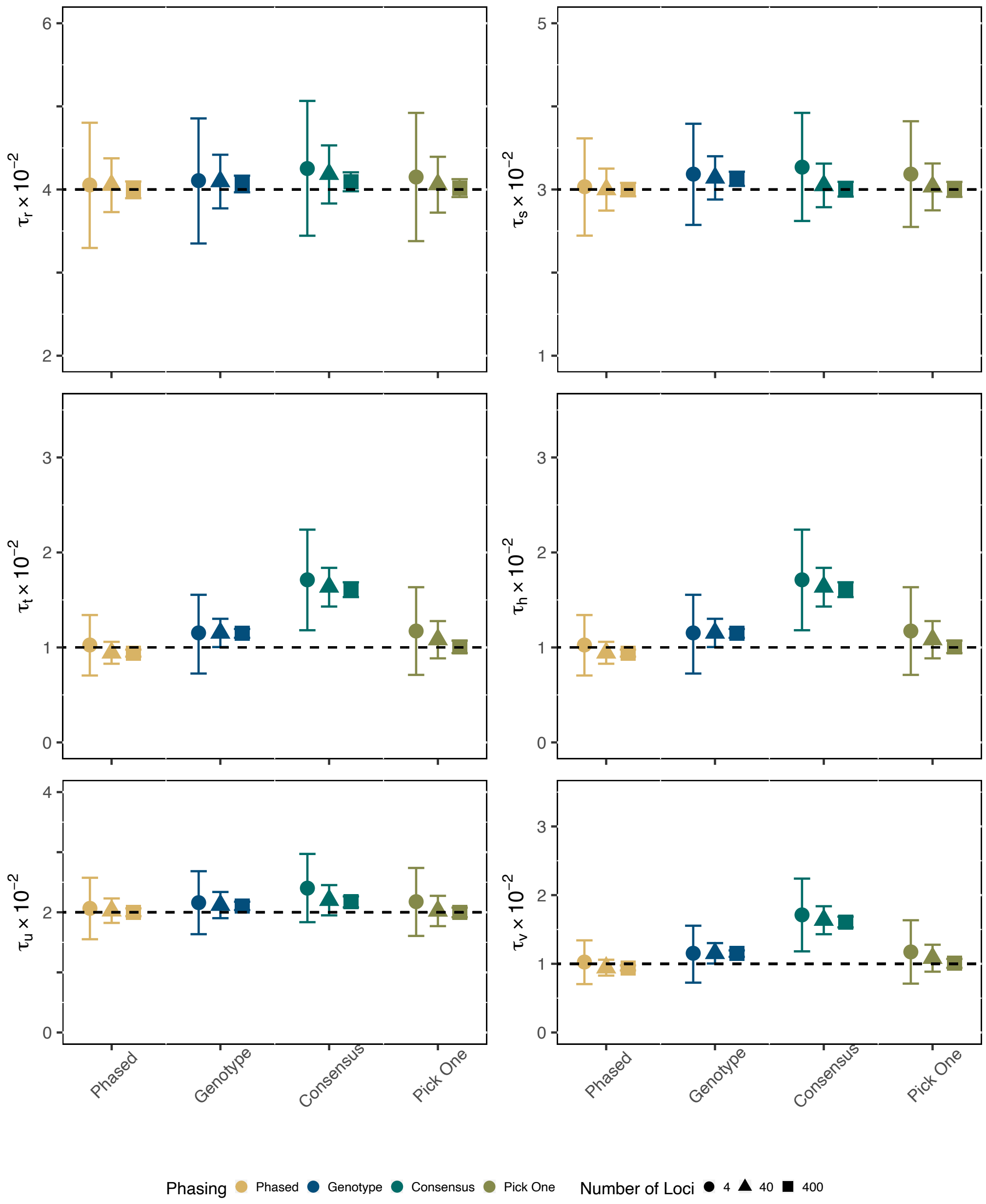


**Figure S8 — Divergence Time Estimation for Species Network with $\tau_h = 0.01$ and $\theta = \tau_h$.** The y-axis is divergence time for each individual parameter. Divergence times are measured in the expected number of substitutions per site. The dashed line represents the true simulated values. Points are posterior means and error bars are 95% HPD intervals, averaged across 100 replicates.

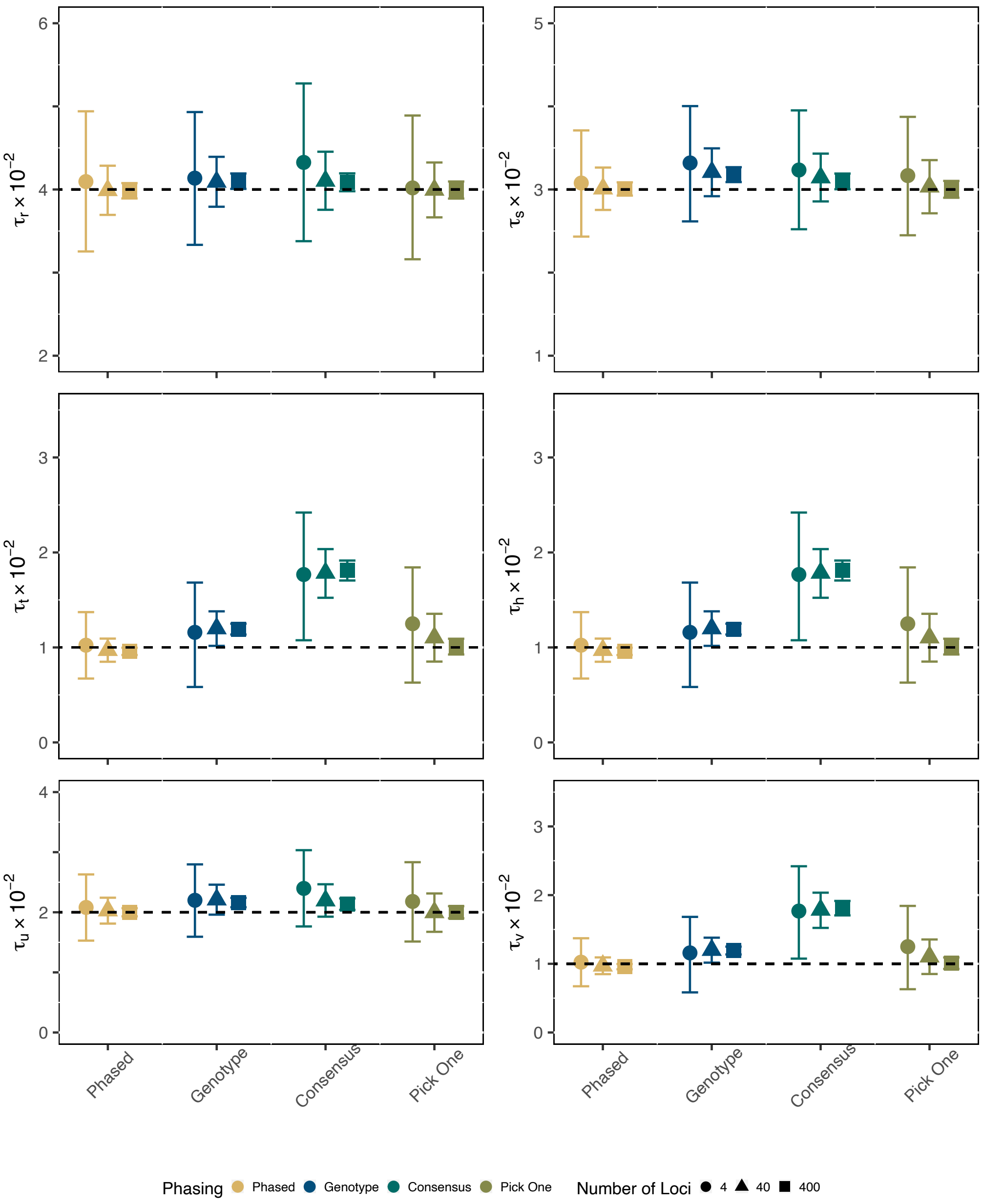


**Figure S9 — Divergence Time Estimation for Species Network with $\tau_h = 0.001$ and $\theta = 2\tau_h$.** The y-axis is divergence time for each individual parameter. Divergence times are measured in the expected number of substitutions per site. The dashed line represents the true simulated values. Points are posterior means and error bars are 95% HPD intervals, averaged across 100 replicates.

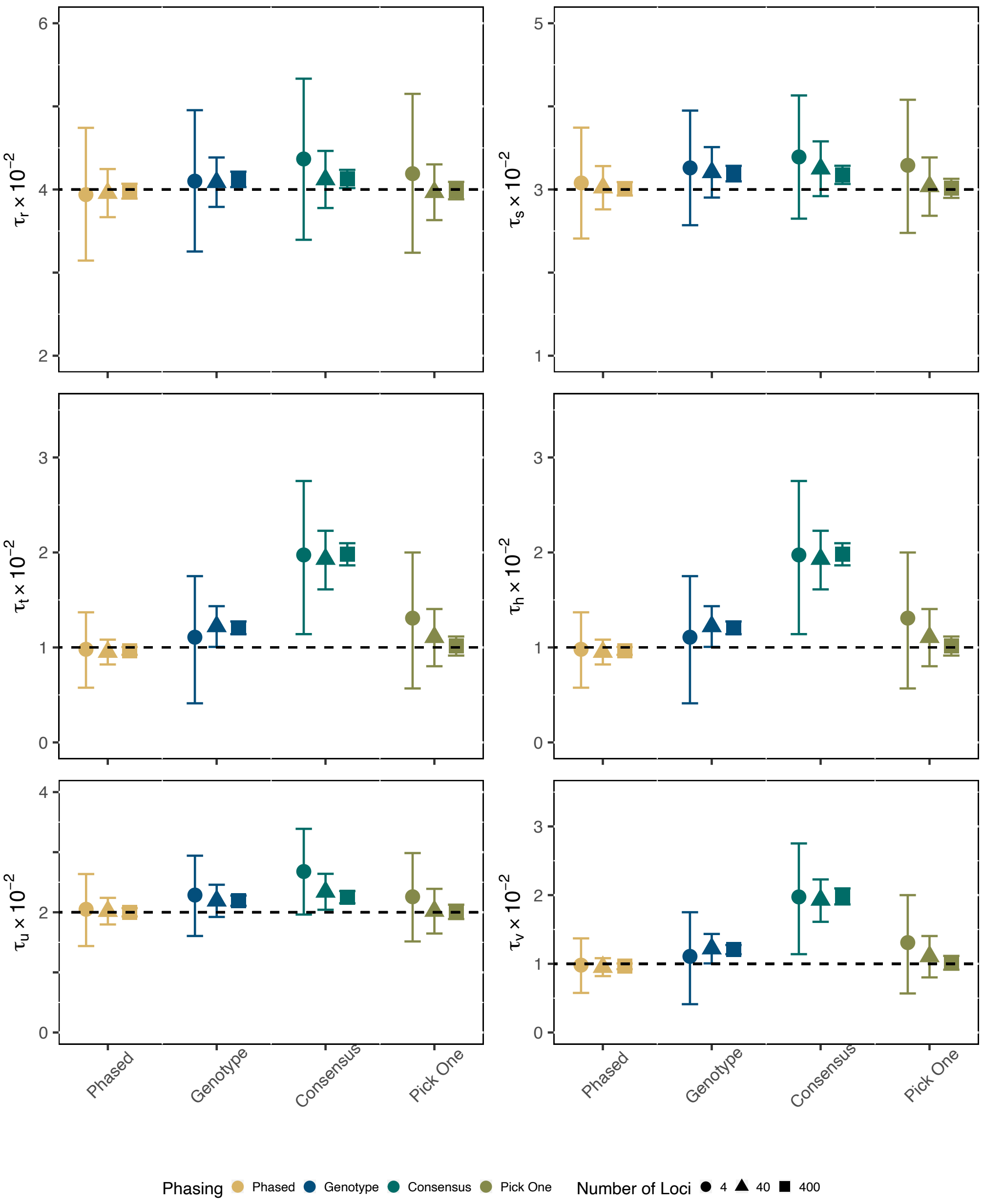


**Figure S10 — Divergence Time Estimation for Species Network with $\tau_h = 0.01$ and $\theta = 3\tau_h$.** The y-axis is divergence time for each individual parameter. Divergence times are measured in the expected number of substitutions per site. The dashed line represents the true simulated values. Points are posterior means and error bars are 95% HPD intervals, averaged across 100 replicates.

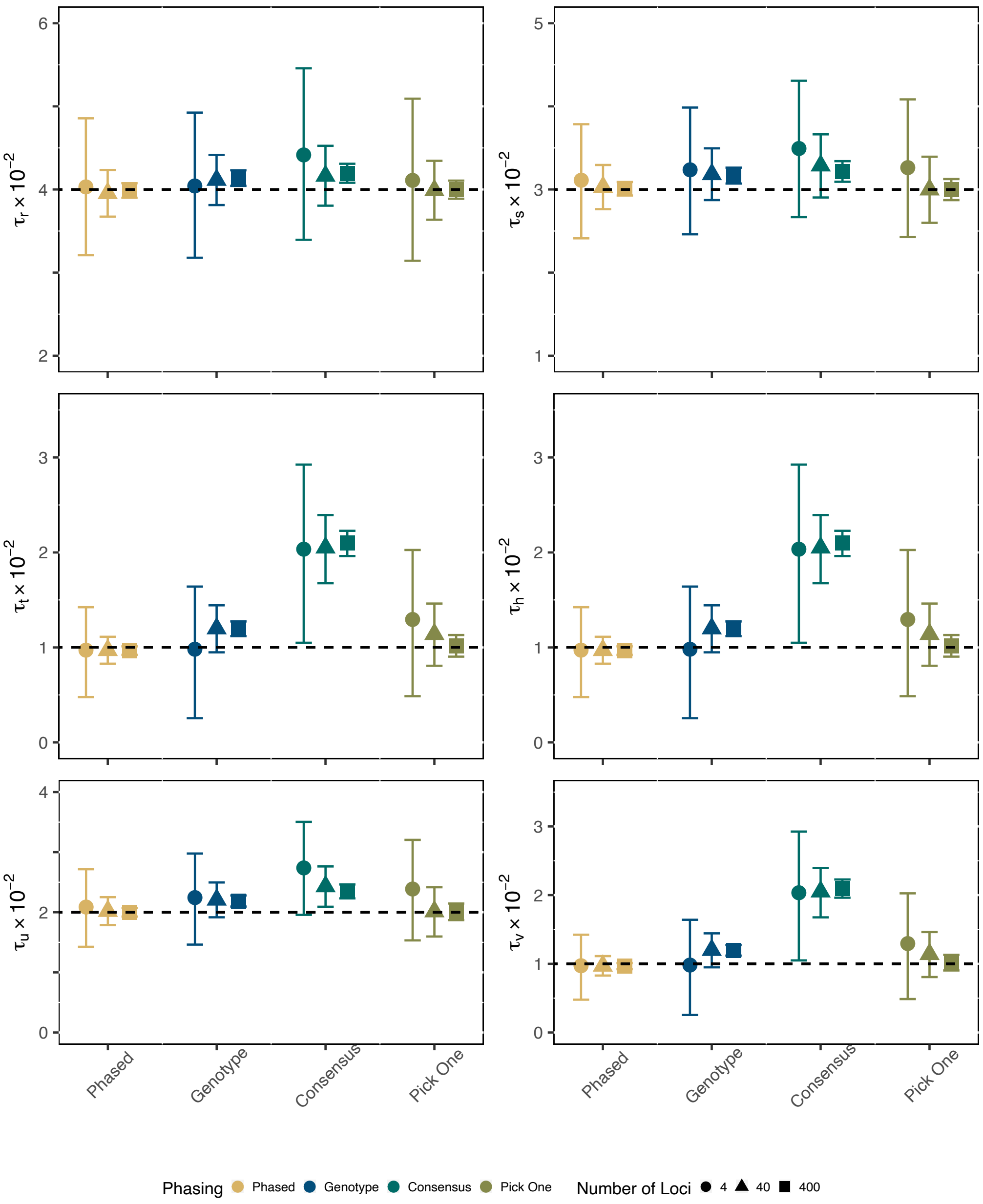


**Figure S11 — Divergence Time Estimation for Species Network with $\tau_h = 0.01$ and $\theta = 4\tau_h$.** The y-axis is divergence time for each individual parameter. Divergence times are measured in the expected number of substitutions per site. The dashed line represents the true simulated values. Points are posterior means and error bars are 95% HPD intervals, averaged across 100 replicates.

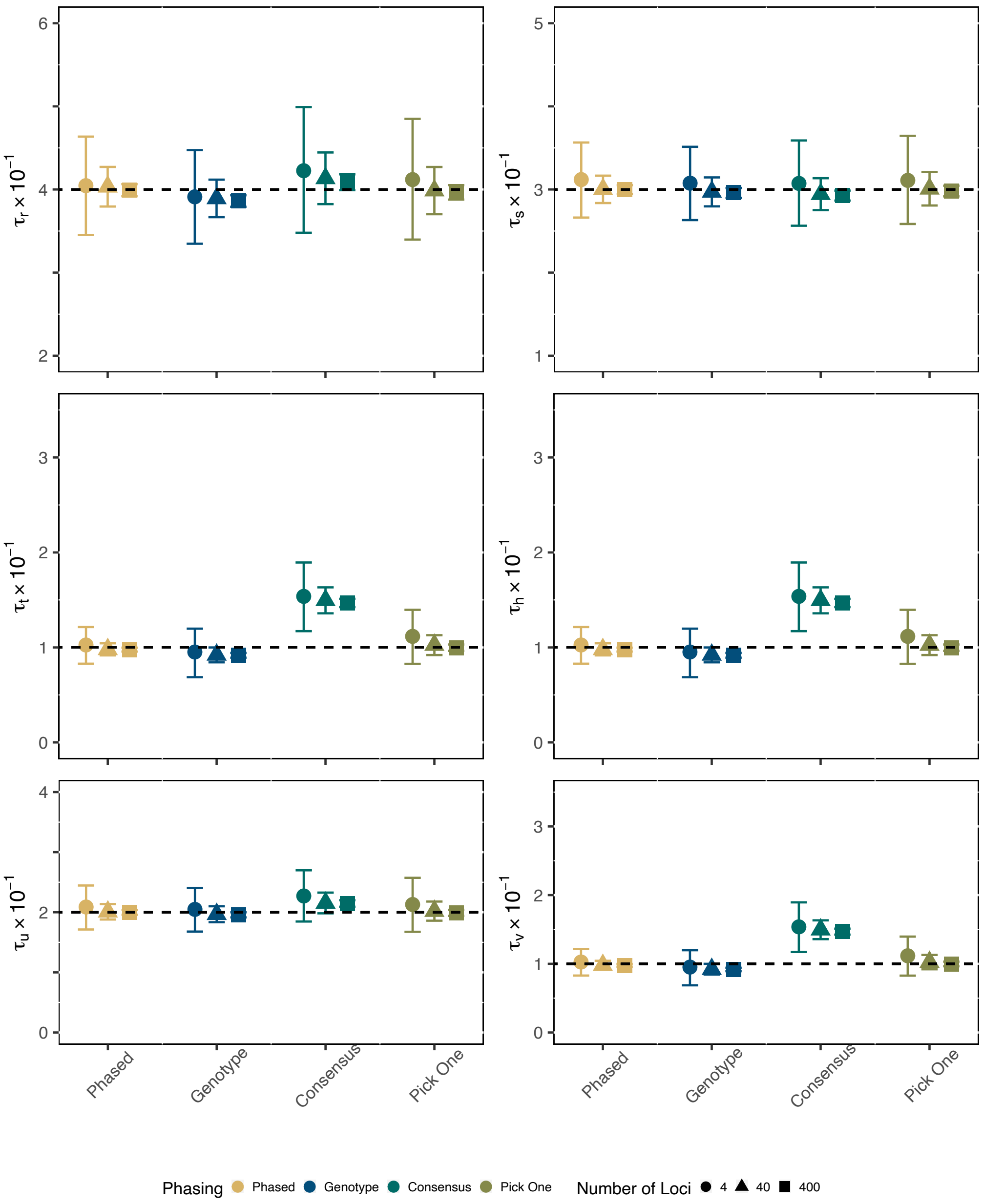


**Figure S12 — Divergence Time Estimation for Species Network with $\tau_h = 0.1$ and $\theta = \tau_h$.** The y-axis is divergence time for each individual parameter. Divergence times are measured in the expected number of substitutions per site. The dashed line represents the true simulated values. Points are posterior means and error bars are 95% HPD intervals, averaged across 100 replicates.

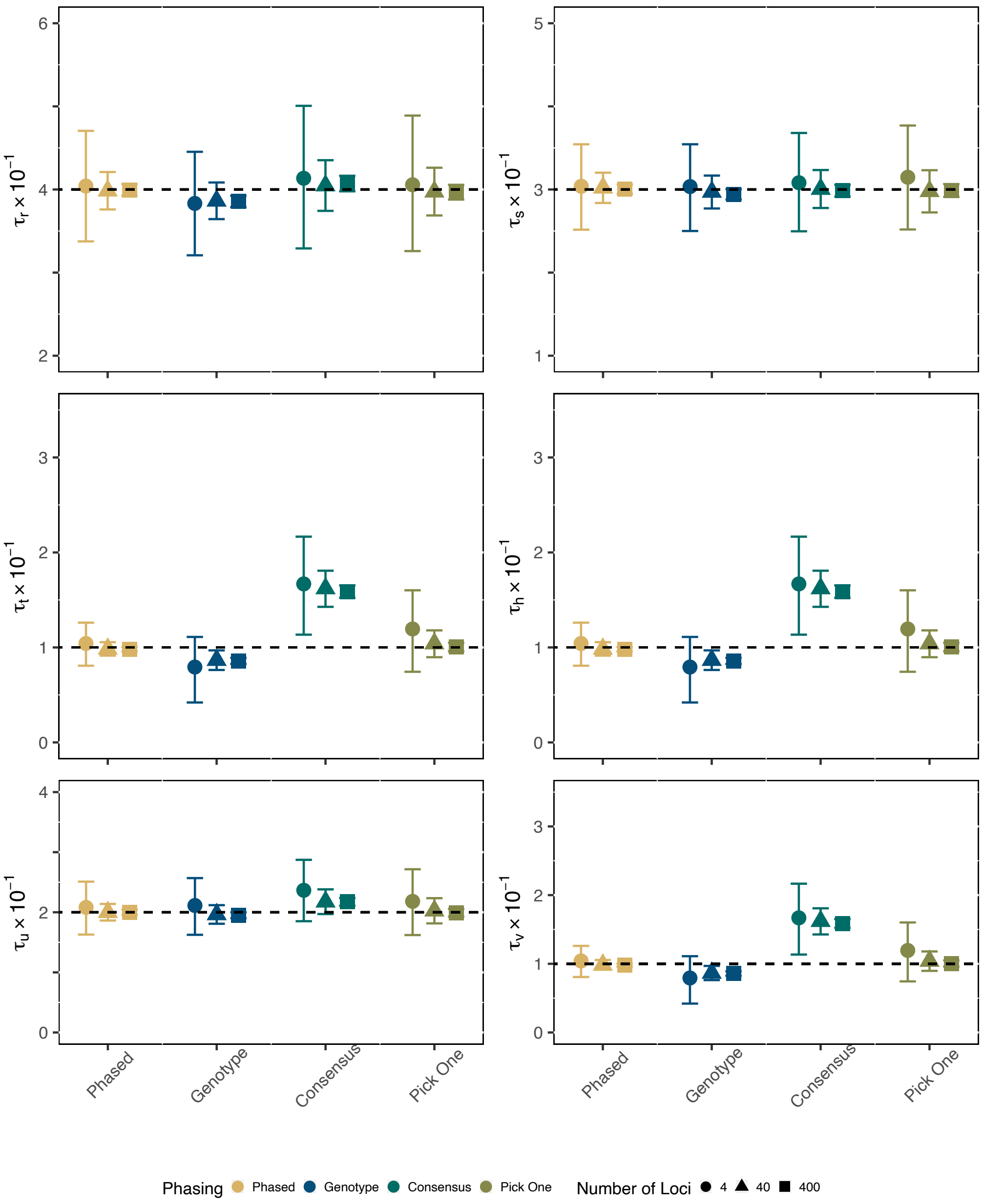


**Figure S13 — Divergence Time Estimation for Species Network with $\tau_h = 0.1$ and $\theta = 2\tau_h$.** The y-axis is divergence time for each individual parameter. Divergence times are measured in the expected number of substitutions per site. The dashed line represents the true simulated values. Points are posterior means and error bars are 95% HPD intervals, averaged across 100 replicates.

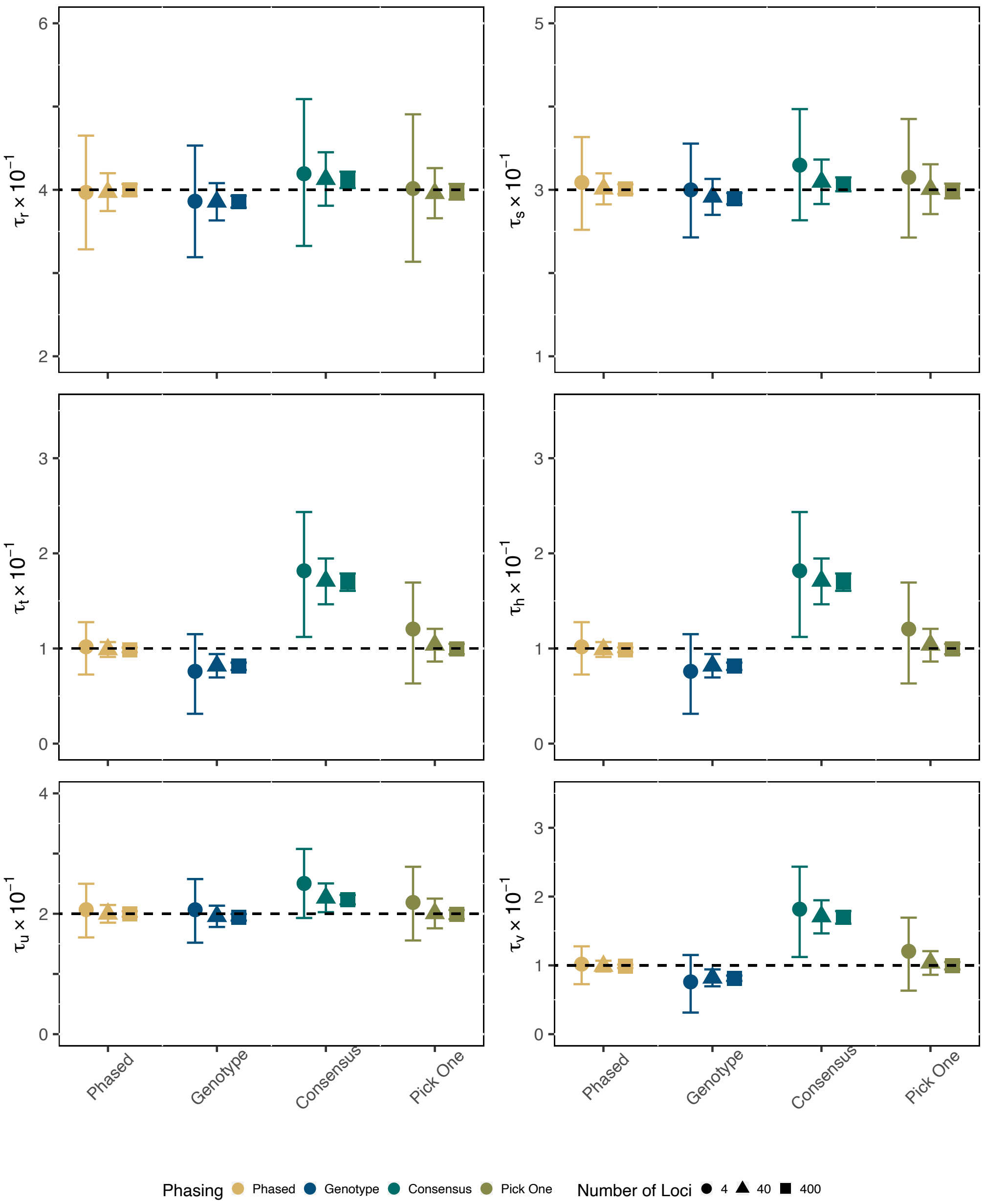


**Figure S14 — Divergence Time Estimation for Species Network with $\tau_h = 0.1$ and $\theta = 3\tau_h$.** The y-axis is divergence time for each individual parameter. Divergence times are measured in the expected number of substitutions per site. The dashed line represents the true simulated values. Points are posterior means and error bars are 95% HPD intervals, averaged across 100 replicates.

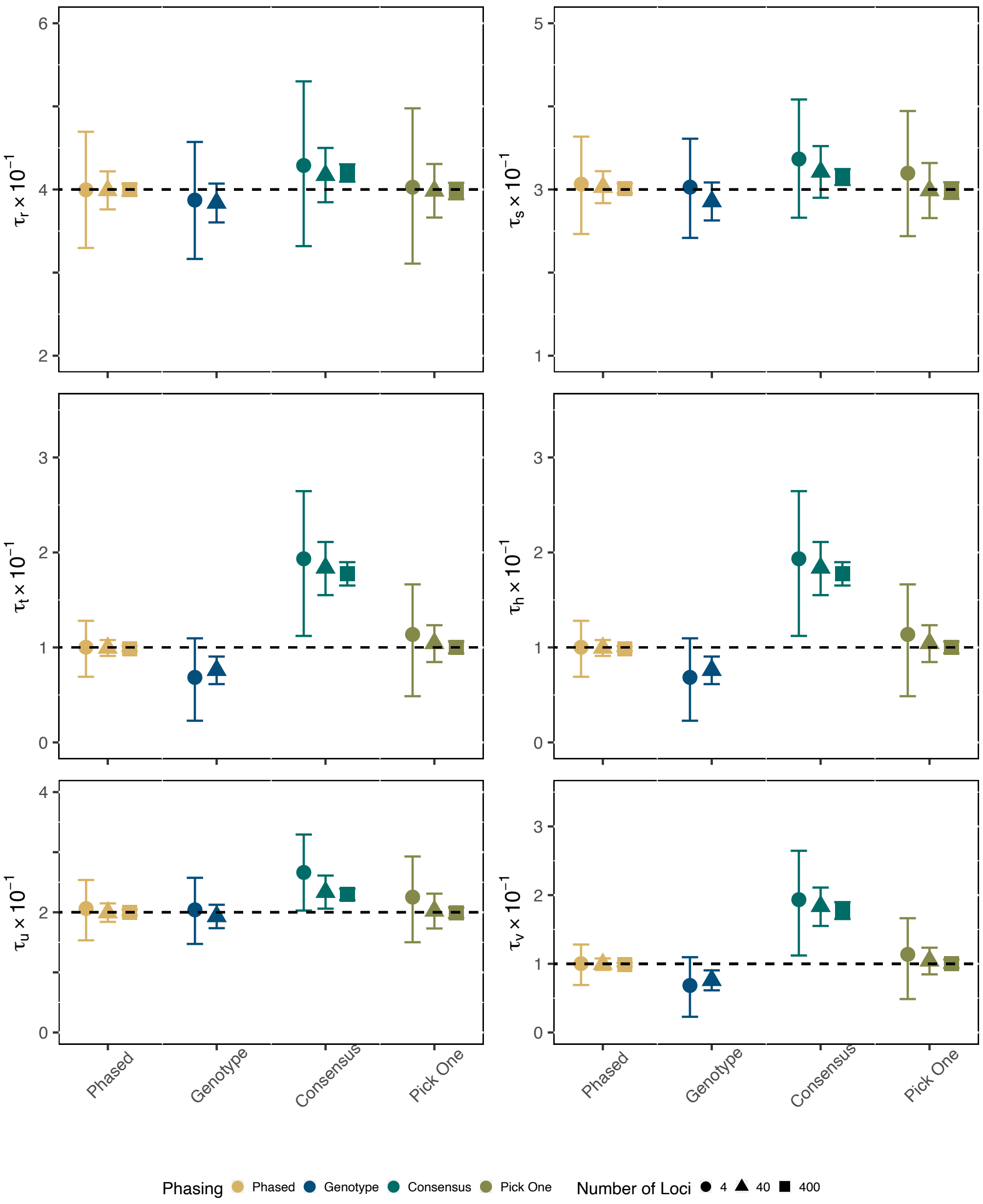


**Figure S15 — Divergence Time Estimation for Species Network with $\tau_h = 0.1$ and $\theta = 4\tau_h$.** The y-axis is divergence time for each individual parameter. Divergence times are measured in the expected number of substitutions per site. The dashed line represents the true simulated values. Points are posterior means and error bars are 95% HPD intervals, averaged across 100 replicates.

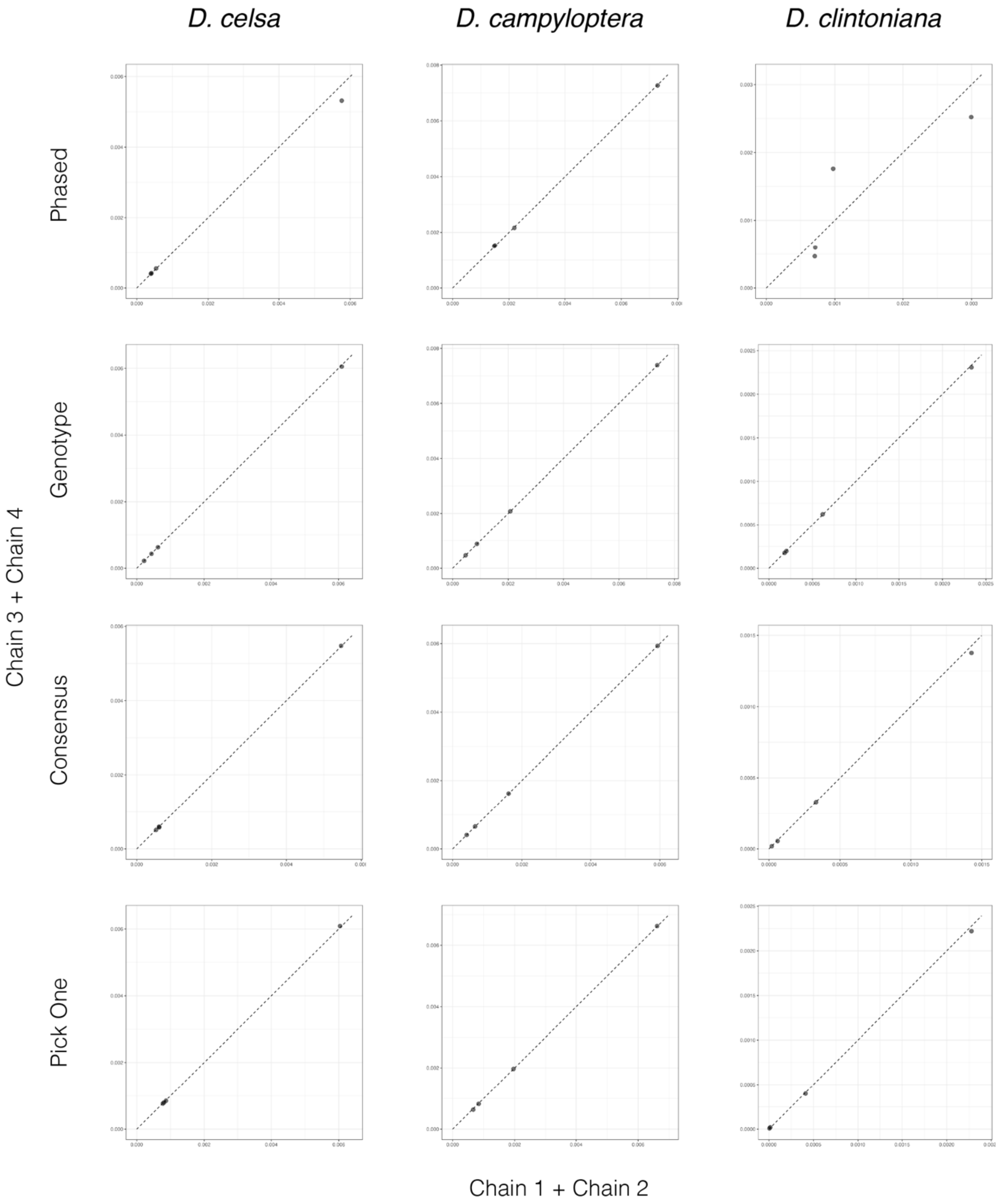


**Figure S16 — Median Node Heights between MCMC Chains.** Scatterplots show the median divergence time parameters in substitutions per site between MCMC chains one and two versus three and four for each data type and triplet. Adherence of data points to the dashed one-to-one line indicates convergence, at least for divergence times.

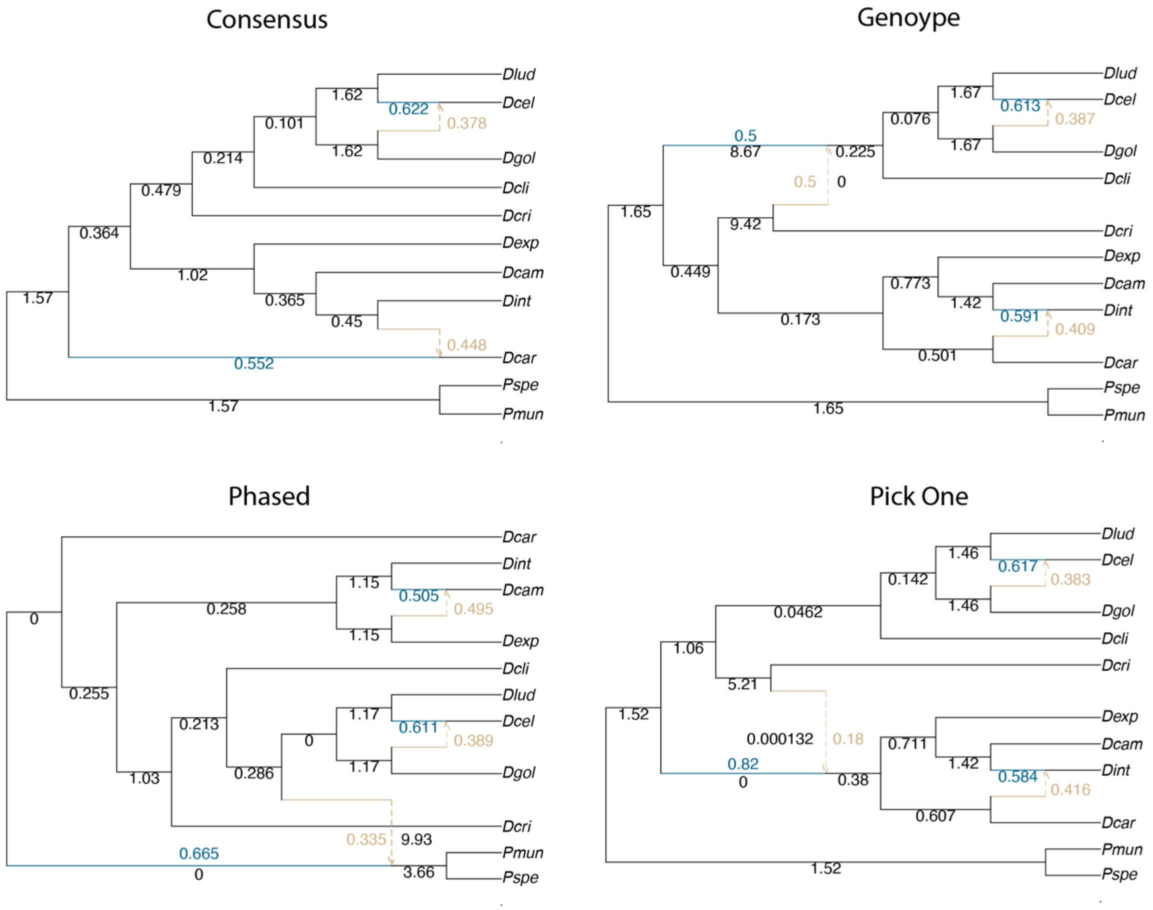


**Figure S17 — SNaQ Networks for *Dryopteris*.** Networks are the same as Figure 7, but identifiable branch lengths are shown in black next to edges. Note that terminal branch lengths are not identifiable. Introgression probabilities for the minor (hybrid) edges are shown in tan while the major edge is shown in blue.

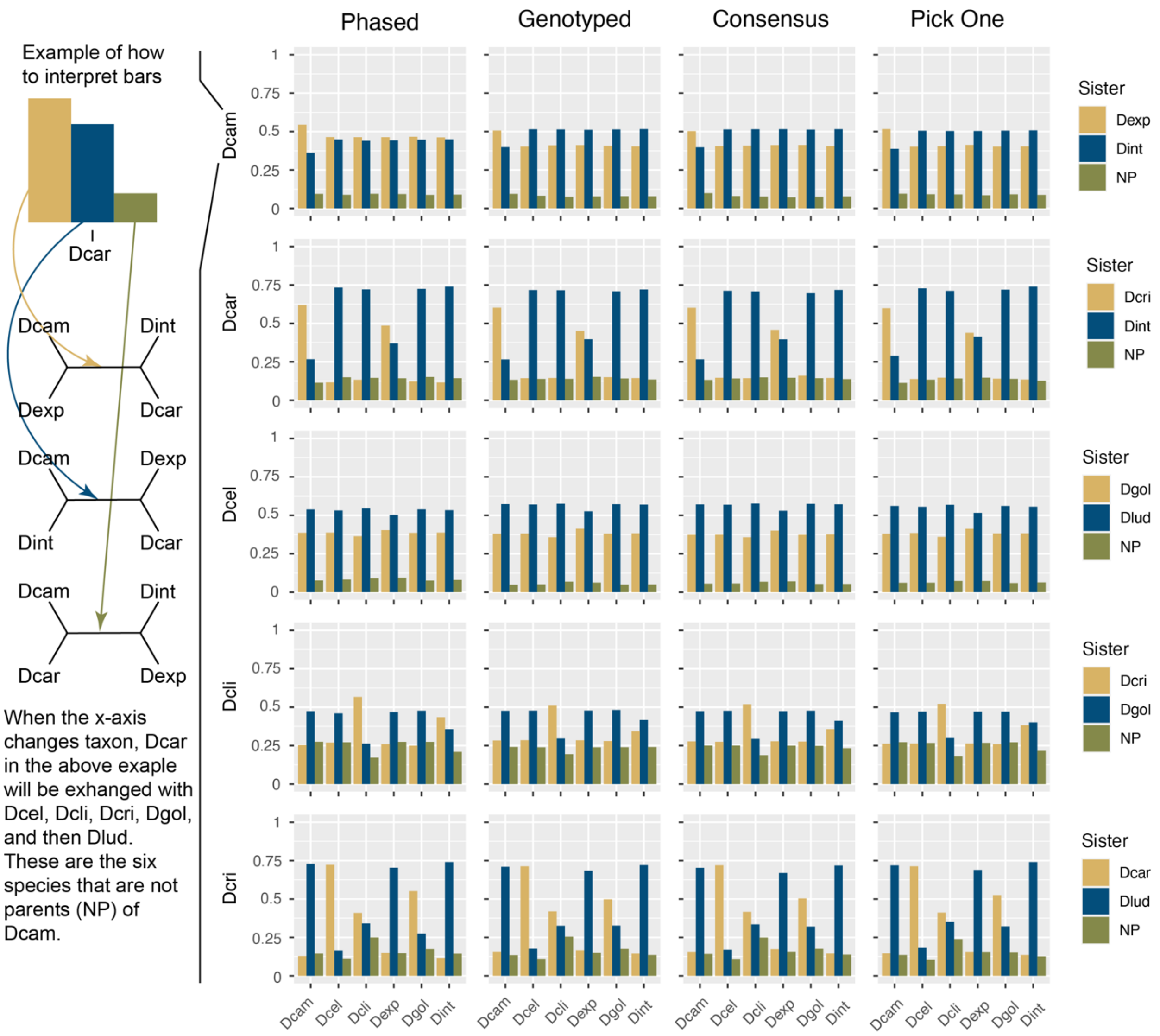


**Figure S18 — Quartet Concordance Factors for Allopolyploids and their Parents.** Taxa abbreviations on the y-axis are the focal allopolyploid in a quartet. Quartets are displayed here only if they include both parents. The x-axis is the fourth member of the quartet that is a non-parent (NP). Bars are concordance factors for the proportion of quartets across all genes that the focal allopolyploid is sister to one of its parents or the non-parent. Non-ILS patterns are observable when the two minor quartets are not approximately equal, but not all patterns are reconciled in the networks. Spectra are similar between phased and unphased data (consensus).

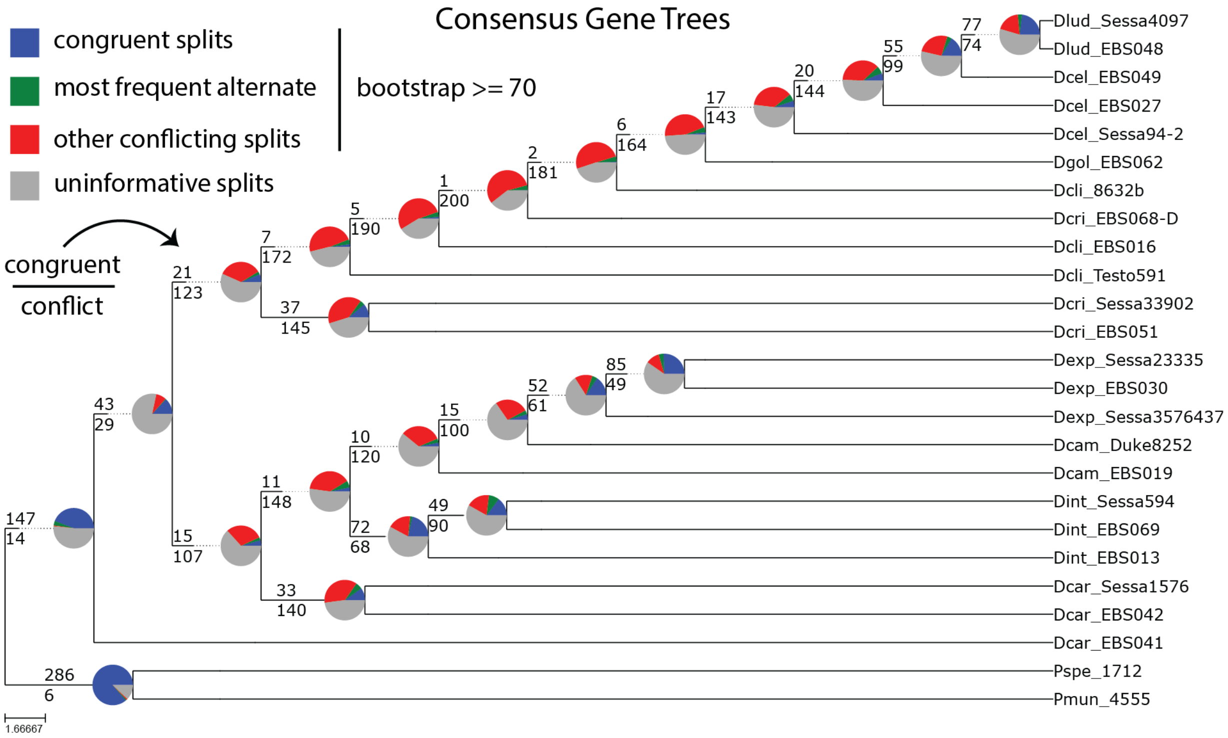


**Figure S19 — Gene Tree Discordance with Consensus Sequences.** Supported splits either in agreement or in conflict with the ASTRAL species tree from phased data was determined by a standard bootstrap proportion of 70% or greater.

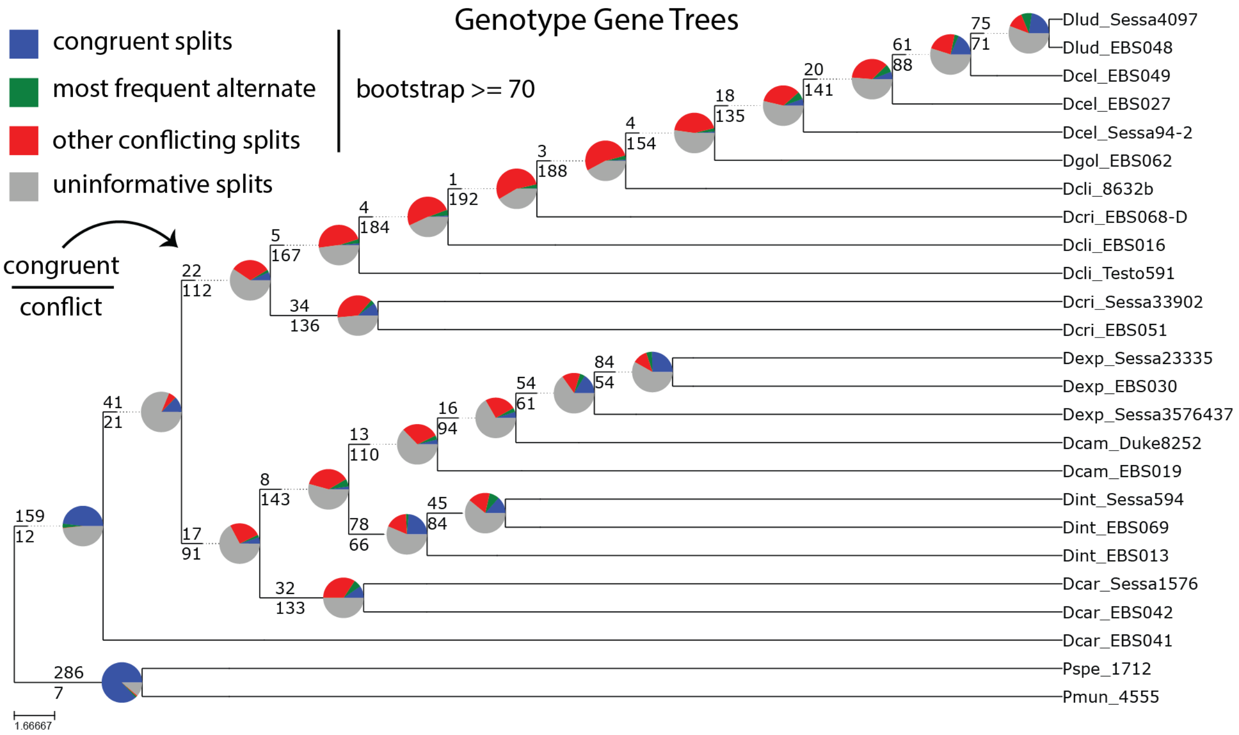


**Figure S20 — Gene Tree Discordance with Genotype Sequences.** Supported splits either in agreement or in conflict with the ASTRAL species tree from phased data was determined by a standard bootstrap proportion of 70% or greater.

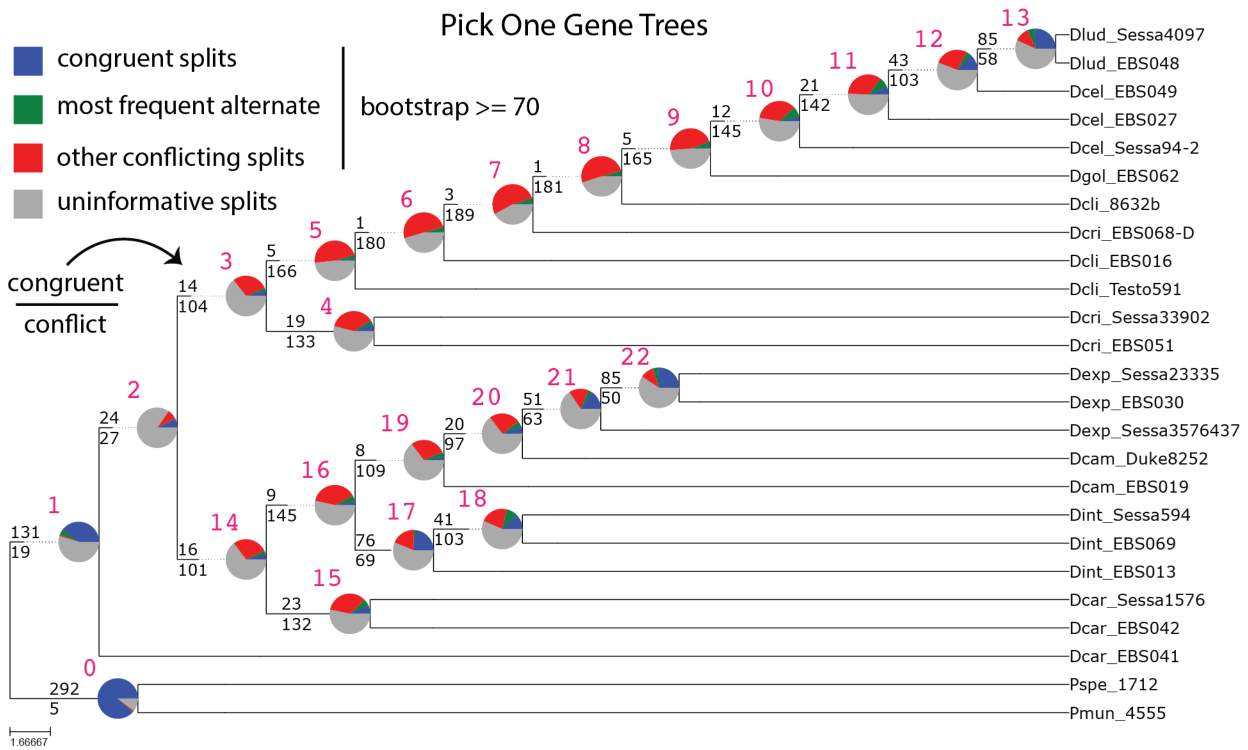


**Figure S21 — Gene Tree Discordance with Pick One Sequences.** Supported splits either in agreement or in conflict with the ASTRAL species tree from phased data was determined by a standard bootstrap proportion of 70% or greater. The hot-pink numbers next to nodes are node numbers that correspond to Supplementary Figure S22.

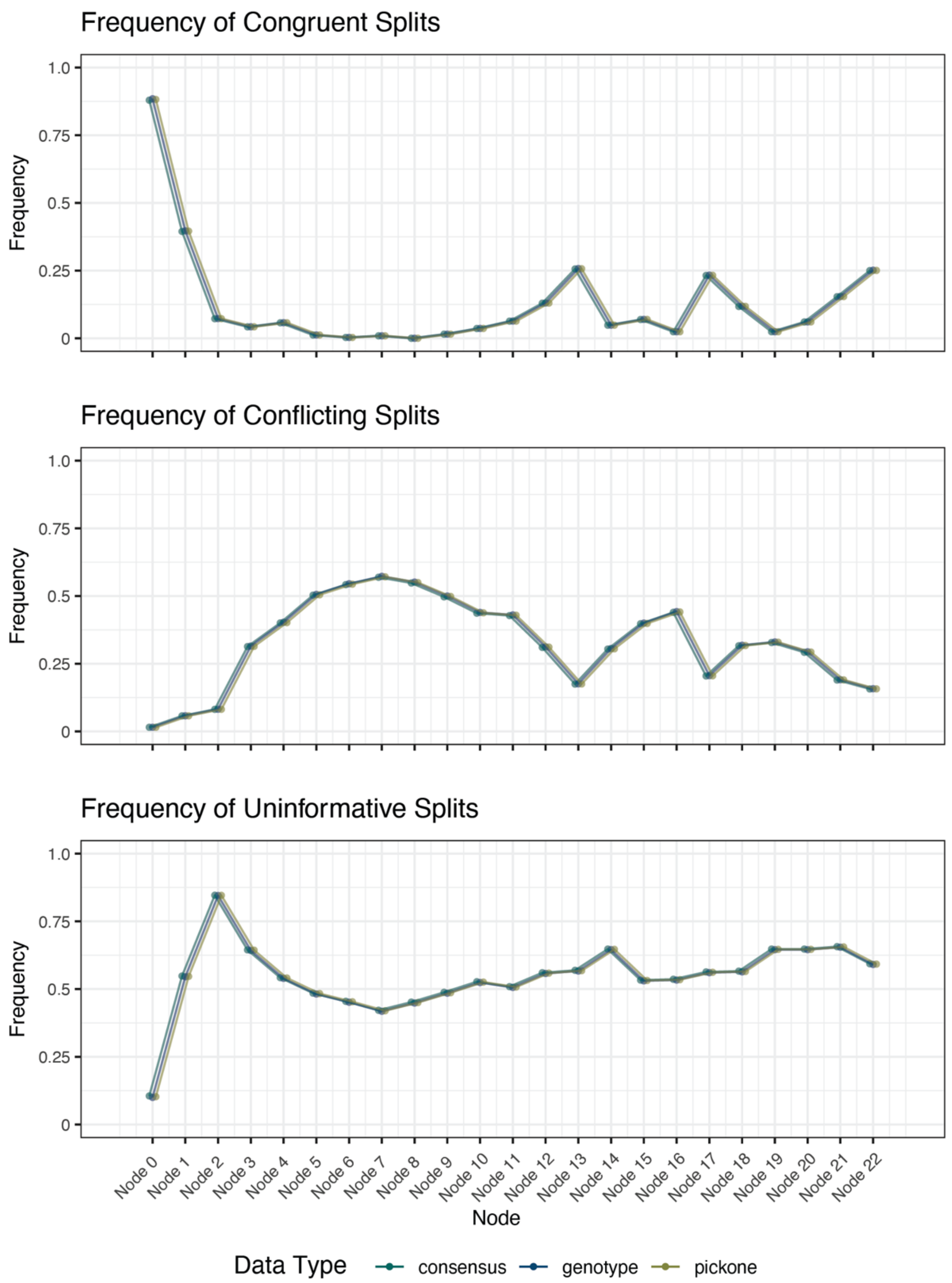


**Figure S22 — Frequency of Informative and Uninformative Splits across Nodes and Data Types.** Phased data are not shown because such conflict analyses are not possible with multiple alleles per species without genome-scale haplotype phasing.

### Supplementary Tables

| ID | Species | Collector | Voucher | Ploidy | SRA |
|---|---|---|---|---|---|
| B087-A08 | *Dryopteris campyloptera* | EB Sessa | EBS019 | 4 | SRS8770054 |
| B087-D08 | *Dryopteris campyloptera* | EB Sessa | EBS022 | 4 | SRS8770074 |
| B125-H06 | *Dryopteris campyloptera* | EB Sessa | Duke catalog 8252 | 4 | SRS8770073 |
| B087-A09 | *Dryopteris carthusiana* | EB Sessa | EBS042 | 4 | SRS8770079 |
| B087-C08 | *Dryopteris carthusiana* | EB Sessa | EBS041 | 4 | SRS8770066 |
| B124-B11 | *Dryopteris carthusiana* | EB Sessa | Sessa Catalog 1576 | 4 | SRS8770067 |
| B087-G08 | *Dryopteris celsa* | EB Sessa | EBS027 | 4 | SRS8770077 |
| B087-H08 | *Dryopteris celsa* | EB Sessa | EBS049 | 4 | SRS8770078 |
| B124-E12 | *Dryopteris celsa* | EB Sessa | Sessa Catalog 94-2 | 4 | SRS8770071 |
| B087-B09 | *Dryopteris clintoniana* | EB Sessa | 8632b | 6 | SRS8770080 |
| B087-C09 | *Dryopteris clintoniana* | EB Sessa | EBS016 | 6 | SRS8770056 |
| B095-B12 | *Dryopteris clintoniana* | W Testo | Testo 591 | 6 | SRS8770062 |
| B087-E09 | *Dryopteris cristata* | EB Sessa | EBS068 - D | 4 | SRS8770058 |
| B087-F09 | *Dryopteris cristata* | EB Sessa | EBS051 | 4 | SRS8770059 |
| B124-C12 | *Dryopteris cristata* | EB Sessa | Sessa Catalog 33902 | 4 | SRS8770069 |
| B087-D09 | *Dryopteris expansa* | EB Sessa | EBS030 | 2 | SRS8770057 |
| B124-B12 | *Dryopteris expansa* | EB Sessa | Sessa Catalog 23335 | 2 | SRS8770068 |
| B125-D06 | *Dryopteris expansa* | EB Sessa | Sessa catalog 3576437 | 2 | SRS8770072 |
| B087-G09 | *Dryopteris goldiana* | EB Sessa | EBS062 | 2 | SRS8770060 |
| B087-E08 | *Dryopteris intermedia* | EB Sessa | EBS069 | 2 | SRS8770075 |
| B087-F08 | *Dryopteris intermedia* | EB Sessa | EBS013 | 2 | SRS8770076 |
| B124-E11 | *Dryopteris intermedia* | EB Sessa | Sessa Catalog 594 | 2 | SRS8770070 |
| B087-H09 | *Dryopteris ludoviciana* | EB Sessa | EBS048 | 2 | SRS8770061 |
| B124-A12 | *Dryopteris ludoviciana* | EB Sessa | Sessa Catalog 4097 | 2 | SRS8770065 |
| B98-A1 | *Polystichum munitum* | CJ Rothfels | 4555 | 2 | SRS8770063 |
| B98-G1 | *Polystichum speciosissimum* | MA Sundue | 1712 | 2 | SRS8770064 |

**Table S1 – Individuals included in study.**

| | | | | | | | | | | | Number of Unique Phased Alleles | | | | | |
|---|---|---|---|---|---|---|---|---|---|---|---|---|---|---|---|---|
| ID | Species | # Loci | # Variable Loci | # Invariable Loci | # Phased Loci | Locus Length | # Variants | Het | # Phased Blocks† | Longest Block‡ | 1* | 2 | 3 | 4 | 5 | 6 |
| B087-A08 | *D. campyloptera* | 404 | 357 | 47 | 308 | 518.58 | 5.30 | 0.0104 | 1.03 | 6.69 | 96 | 51 | 109 | 148 | NA | NA |
| B087-B08 | *D. goldiana* | 403 | 258 | 145 | 199 | 1165.57 | 7.02 | 0.0064 | 1.02 | 13.81 | 204 | 199 | NA | NA | NA | NA |
| B087-C08 | *D. carthusiana* | 408 | 337 | 71 | 289 | 1249.54 | 9.92 | 0.0086 | 1.02 | 13.71 | 119 | 43 | 88 | 158 | NA | NA |
| B087-D08 | *D. campyloptera* | 121 | 62 | 59 | 50 | 564.39 | 7.34 | 0.0133 | 1.00 | 17.50 | 71 | 8 | 8 | 34 | NA | NA |
| B087-E08 | *D. intermedia* | 407 | 200 | 207 | 120 | 622.50 | 2.52 | 0.0037 | 1.01 | 7.74 | 287 | 120 | NA | NA | NA | NA |
| B087-F08 | *D. intermedia* | 403 | 178 | 225 | 124 | 609.41 | 2.36 | 0.0036 | 1.04 | 7.02 | 279 | 124 | NA | NA | NA | NA |
| B087-G08 | *D. celsa* | 409 | 376 | 33 | 366 | 802.90 | 10.61 | 0.0138 | 1.01 | 11.77 | 43 | 40 | 112 | 214 | NA | NA |
| B087-H08 | *D. celsa* | 409 | 375 | 34 | 365 | 944.28 | 11.95 | 0.0133 | 1.01 | 13.33 | 44 | 42 | 100 | 223 | NA | NA |
| B087-A09 | *D. carthusiana* | 408 | 379 | 29 | 370 | 845.50 | 13.15 | 0.0163 | 1.01 | 14.45 | 38 | 34 | 106 | 230 | NA | NA |
| B087-B09 | *D. clintoniana* | 409 | 402 | 7 | 396 | 569.66 | 12.83 | 0.0230 | 1.01 | 13.21 | 13 | 7 | 20 | 38 | 116 | 215 |
| B087-C09 | *D. clintoniana* | 410 | 390 | 20 | 380 | 927.59 | 21.64 | 0.0237 | 1.01 | 23.31 | 30 | 5 | 10 | 26 | 68 | 271 |
| B087-D09 | *D. expansa* | 399 | 121 | 278 | 77 | 501.36 | 1.65 | 0.0032 | 1.01 | 7.90 | 322 | 77 | NA | NA | NA | NA |
| B087-E09 | *D. cristata* | 409 | 394 | 15 | 389 | 728.63 | 16.39 | 0.0236 | 1.01 | 17.20 | 20 | 9 | 42 | 338 | NA | NA |
| B087-F09 | *D. cristata* | 409 | 374 | 35 | 366 | 813.84 | 15.67 | 0.0201 | 1.01 | 17.45 | 43 | 30 | 96 | 240 | NA | NA |
| B087-G09 | *D. goldiana* | 410 | 108 | 302 | 79 | 961.39 | 2.29 | 0.0025 | 1.01 | 11.37 | 331 | 79 | NA | NA | NA | NA |
| B087-H09 | *D. ludoviciana* | 410 | 184 | 226 | 105 | 1010.95 | 2.93 | 0.0030 | 1.02 | 10.57 | 305 | 105 | NA | NA | NA | NA |
| B095-B12 | *D. clintoniana* | 412 | 395 | 17 | 384 | 839.66 | 19.22 | 0.0233 | 1.01 | 20.55 | 28 | 6 | 14 | 26 | 72 | 266 |
| B98-A1 | *P. munitum* | 409 | 320 | 89 | 251 | 951.47 | 6.01 | 0.0065 | 1.01 | 9.46 | 158 | 251 | NA | NA | NA | NA |
| B98-G1 | *P. speciosissimum* | 406 | 187 | 219 | 108 | 785.40 | 3.37 | 0.0041 | 1.02 | 11.81 | 298 | 108 | NA | NA | NA | NA |
| B124-A12 | *D. ludoviciana* | 405 | 139 | 266 | 91 | 795.13 | 2.51 | 0.0034 | 1.00 | 10.63 | 314 | 91 | NA | NA | NA | NA |
| B124-B11 | *D. carthusiana* | 405 | 399 | 6 | 390 | 597.90 | 10.36 | 0.0177 | 1.00 | 10.72 | 15 | 19 | 75 | 296 | NA | NA |
| B124-B12 | *D. expansa* | 404 | 153 | 251 | 97 | 623.04 | 2.71 | 0.0042 | 1.00 | 10.67 | 307 | 97 | NA | NA | NA | NA |
| B124-C12 | *D. cristata* | 405 | 388 | 17 | 377 | 624.96 | 11.61 | 0.0190 | 1.00 | 12.43 | 28 | 33 | 107 | 237 | NA | NA |
| B124-E11 | *D. intermedia* | 405 | 245 | 160 | 179 | 856.78 | 4.28 | 0.0049 | 1.01 | 9.21 | 226 | 179 | NA | NA | NA | NA |
| B124-E12 | *D. celsa* | 407 | 391 | 16 | 373 | 709.76 | 9.53 | 0.0138 | 1.01 | 10.32 | 34 | 35 | 102 | 236 | NA | NA |
| B125-D06 | *D. expansa* | 395 | 190 | 205 | 117 | 381.70 | 2.19 | 0.0053 | 1.00 | 6.72 | 278 | 117 | NA | NA | NA | NA |
| B125-H06 | *D. campyloptera* | 390 | 364 | 26 | 344 | 709.58 | 7.90 | 0.0113 | 1.01 | 8.88 | 46 | 45 | 120 | 179 | NA | NA |

† The average number of haplotype blocks recoverd for each locus where a haplotype block exists - not all loci have phased haplotype blocks

‡ The average number of variants per haplotype block. When more than one block per locus, only the phased variants of the longest block were retained

* One allele may not be stricktly homozygous and thus the counts are higher than the number of invariable loci

**Table S2 – Per-individual Phasing Statistics.**

| Locus | Phased Sequences # Seqs | Phased Sequences # Sites | Phased Sequences # PI Sites | Consensus Sequences # Seqs | Consensus Sequences # Sites | Consensus Sequences # PI Sites | Genotype Sequences # Seqs | Genotype Sequences # Sites | Genotype Sequences # PI Sites | Pick One Sequences # Seqs | Pick One Sequences # Sites | Pick One Sequences # PI Sites |
|---|---|---|---|---|---|---|---|---|---|---|---|---|
| L1 | 84 | 1469 | 324 | 25 | 1456 | 61 | 25 | 1442 | 58 | 25 | 1469 | 65 |
| L100 | 84 | 1280 | 543 | 25 | 1303 | 232 | 25 | 1300 | 221 | 25 | 1276 | 220 |
| L101 | 84 | 1301 | 457 | 25 | 1292 | 243 | 25 | 1304 | 247 | 25 | 1350 | 222 |
| L102 | 80 | 1397 | 476 | 24 | 1330 | 253 | 24 | 1332 | 245 | 24 | 1335 | 253 |
| L103 | 84 | 1659 | 575 | 25 | 1659 | 101 | 25 | 1659 | 100 | 25 | 1683 | 103 |
| L105 | 84 | 1251 | 274 | 25 | 1240 | 84 | 25 | 1247 | 81 | 25 | 1247 | 85 |
| L107 | 84 | 1997 | 568 | 25 | 2007 | 184 | 25 | 2007 | 183 | 25 | 2018 | 178 |
| L11 | 84 | 1692 | 666 | 25 | 1722 | 182 | 25 | 1726 | 157 | 25 | 1760 | 166 |
| L112 | 84 | 1957 | 525 | 25 | 1885 | 123 | 25 | 1885 | 153 | 25 | 1949 | 101 |
| L113 | 84 | 1995 | 450 | 25 | 1992 | 138 | 25 | 1992 | 135 | 25 | 1993 | 151 |
| L114 | 84 | 1559 | 792 | 25 | 1659 | 385 | 25 | 1536 | 379 | 25 | 1454 | 407 |
| L115 | 84 | 1674 | 604 | 25 | 1703 | 165 | 25 | 1652 | 165 | 25 | 1705 | 183 |
| L116 | 84 | 1794 | 382 | 25 | 1777 | 84 | 25 | 1764 | 95 | 25 | 1764 | 98 |
| L117 | 84 | 1294 | 363 | 25 | 1294 | 62 | 25 | 1294 | 61 | 25 | 1294 | 68 |
| L118 | 84 | 3398 | 894 | 25 | 3333 | 370 | 25 | 3332 | 362 | 25 | 3285 | 375 |
| L12 | 82 | 1696 | 639 | 24 | 1682 | 242 | 24 | 1714 | 234 | 24 | 1727 | 256 |
| L121 | 84 | 1460 | 455 | 25 | 1462 | 98 | 25 | 1462 | 96 | 25 | 1458 | 98 |
| L123 | 84 | 1491 | 554 | 25 | 1496 | 214 | 25 | 1496 | 199 | 25 | 1486 | 246 |
| L125 | 84 | 2009 | 611 | 25 | 1884 | 192 | 25 | 2026 | 181 | 25 | 2005 | 195 |
| L127 | 84 | 1630 | 469 | 25 | 1645 | 77 | 25 | 1713 | 70 | 25 | 1709 | 70 |
| L129 | 84 | 1663 | 782 | 25 | 1630 | 320 | 25 | 1729 | 264 | 25 | 1700 | 285 |
| L130 | 84 | 1584 | 493 | 25 | 1583 | 117 | 25 | 1583 | 107 | 25 | 1583 | 108 |
| L131 | 84 | 1589 | 423 | 25 | 1589 | 79 | 25 | 1589 | 78 | 25 | 1589 | 78 |
| L132 | 84 | 1398 | 460 | 25 | 1398 | 93 | 25 | 1398 | 90 | 25 | 1393 | 90 |
| L133 | 84 | 1323 | 478 | 25 | 1322 | 117 | 25 | 1322 | 113 | 25 | 1322 | 113 |
| L135 | 84 | 1643 | 492 | 25 | 1639 | 80 | 25 | 1639 | 82 | 25 | 1644 | 77 |
| L137 | 84 | 1214 | 463 | 25 | 1223 | 118 | 25 | 1214 | 141 | 25 | 1216 | 131 |
| L138 | 84 | 1912 | 295 | 25 | 1911 | 82 | 25 | 1911 | 82 | 25 | 1910 | 84 |
| L139 | 84 | 1421 | 637 | 25 | 1328 | 174 | 25 | 1338 | 167 | 25 | 1436 | 166 |
| L14 | 84 | 1435 | 391 | 25 | 1431 | 128 | 25 | 1431 | 128 | 25 | 1429 | 111 |
| L140 | 82 | 1375 | 655 | 24 | 1414 | 231 | 24 | 1316 | 261 | 24 | 1408 | 219 |
| L141 | 84 | 1272 | 307 | 25 | 1270 | 119 | 25 | 1273 | 112 | 25 | 1274 | 72 |
| L142 | 84 | 1531 | 618 | 25 | 1543 | 115 | 25 | 1543 | 119 | 25 | 1520 | 115 |
| L143 | 84 | 1701 | 615 | 25 | 1716 | 159 | 25 | 1716 | 174 | 25 | 1767 | 219 |
| L144 | 84 | 1364 | 715 | 25 | 1425 | 261 | 25 | 1426 | 244 | 25 | 1452 | 218 |
| L145 | 84 | 1424 | 562 | 25 | 1410 | 125 | 25 | 1365 | 138 | 25 | 1419 | 152 |
| L146 | 84 | 1727 | 441 | 25 | 1731 | 94 | 25 | 1726 | 92 | 25 | 1722 | 107 |
| L147 | 84 | 1368 | 506 | 25 | 1384 | 81 | 25 | 1376 | 78 | 25 | 1385 | 83 |
| L148 | 84 | 1378 | 522 | 25 | 1353 | 141 | 25 | 1311 | 131 | 25 | 1359 | 143 |
| L149 | 84 | 1240 | 594 | 25 | 1289 | 140 | 25 | 1289 | 140 | 25 | 1289 | 117 |
| L15 | 84 | 1524 | 254 | 25 | 1496 | 85 | 25 | 1527 | 86 | 25 | 1509 | 95 |
| L151 | 84 | 1450 | 581 | 25 | 1211 | 294 | 25 | 1320 | 277 | 25 | 1346 | 305 |
| L152 | 84 | 1745 | 581 | 25 | 1740 | 158 | 25 | 1736 | 158 | 25 | 1746 | 128 |
| L154 | 84 | 1384 | 416 | 25 | 1374 | 117 | 25 | 1375 | 124 | 25 | 1374 | 127 |
| L155 | 84 | 1690 | 458 | 25 | 1702 | 134 | 25 | 1702 | 133 | 25 | 1687 | 140 |
| L157 | 84 | 1608 | 395 | 25 | 1613 | 84 | 25 | 1605 | 71 | 25 | 1605 | 82 |
| L159 | 80 | 1569 | 541 | 24 | 1569 | 111 | 24 | 1569 | 111 | 24 | 1569 | 90 |
| L161 | 84 | 1471 | 444 | 25 | 1471 | 140 | 25 | 1499 | 138 | 25 | 1471 | 153 |
| L163 | 84 | 1919 | 708 | 25 | 1882 | 413 | 25 | 1888 | 385 | 25 | 1889 | 363 |
| L164 | 84 | 1183 | 390 | 25 | 1183 | 60 | 25 | 1183 | 60 | 25 | 1183 | 62 |
| L165 | 70 | 1177 | 370 | 20 | 1130 | 132 | 20 | 1065 | 187 | 20 | 1098 | 179 |
| L166 | 84 | 1112 | 431 | 25 | 1113 | 93 | 25 | 1113 | 92 | 25 | 1113 | 100 |
| L167 | 82 | 1398 | 481 | 24 | 1413 | 104 | 24 | 1413 | 102 | 24 | 1413 | 108 |
| L168 | 84 | 1462 | 353 | 25 | 1458 | 96 | 25 | 1464 | 98 | 25 | 1464 | 99 |
| L169 | 84 | 1576 | 719 | 25 | 1581 | 257 | 25 | 1653 | 234 | 25 | 1654 | 251 |
| L17 | 84 | 1369 | 442 | 25 | 1369 | 69 | 25 | 1369 | 69 | 25 | 1368 | 67 |
| L170 | 84 | 1541 | 833 | 25 | 1513 | 264 | 25 | 1563 | 237 | 25 | 1504 | 263 |
| L171 | 84 | 1352 | 612 | 25 | 1338 | 180 | 25 | 1398 | 171 | 25 | 1287 | 206 |
| L172 | 84 | 1642 | 509 | 25 | 1595 | 130 | 25 | 1594 | 132 | 25 | 1590 | 129 |
| L173 | 84 | 1416 | 512 | 25 | 1423 | 123 | 25 | 1416 | 115 | 25 | 1423 | 138 |
| L174 | 80 | 1329 | 341 | 24 | 1329 | 83 | 24 | 1329 | 82 | 24 | 1329 | 81 |

| | | | | | | | | | | | | |
|---|---|---|---|---|---|---|---|---|---|---|---|---|
| L175 | 84 | 1537 | 452 | 25 | 1527 | 182 | 25 | 1523 | 173 | 25 | 1534 | 186 |
| L177 | 84 | 1562 | 753 | 25 | 1566 | 312 | 25 | 1584 | 320 | 25 | 1585 | 289 |
| L179 | 84 | 1602 | 583 | 25 | 1568 | 153 | 25 | 1561 | 142 | 25 | 1581 | 160 |
| L18 | 84 | 1988 | 738 | 25 | 1942 | 208 | 25 | 1953 | 218 | 25 | 1899 | 256 |
| L181 | 82 | 1758 | 569 | 24 | 1718 | 144 | 24 | 1726 | 159 | 24 | 1719 | 149 |
| L182 | 30 | 1238 | 299 | 10 | 1238 | 48 | 10 | 1238 | 49 | 10 | 1238 | 54 |
| L183 | 84 | 1634 | 549 | 25 | 1645 | 162 | 25 | 1647 | 156 | 25 | 1657 | 164 |
| L184 | 84 | 1153 | 291 | 25 | 1157 | 132 | 25 | 1153 | 131 | 25 | 1153 | 128 |
| L185 | 84 | 1353 | 528 | 25 | 1412 | 199 | 25 | 1420 | 192 | 25 | 1231 | 127 |
| L186 | 84 | 1500 | 529 | 25 | 1410 | 221 | 25 | 1411 | 213 | 25 | 1410 | 244 |
| L187 | 84 | 1338 | 685 | 25 | 1332 | 210 | 25 | 1281 | 225 | 25 | 1240 | 217 |
| L188 | 84 | 1013 | 282 | 25 | 1028 | 83 | 25 | 1026 | 87 | 25 | 1029 | 81 |
| L189 | 84 | 1295 | 365 | 25 | 1288 | 149 | 25 | 1240 | 152 | 25 | 1277 | 153 |
| L19 | 84 | 1876 | 1024 | 25 | 2038 | 516 | 25 | 1916 | 462 | 25 | 2001 | 426 |
| L191 | 84 | 1550 | 349 | 25 | 1520 | 91 | 25 | 1522 | 88 | 25 | 1513 | 90 |
| L193 | 84 | 1704 | 566 | 25 | 1706 | 131 | 25 | 1673 | 116 | 25 | 1681 | 134 |
| L194 | 84 | 1727 | 683 | 25 | 1735 | 166 | 25 | 1758 | 160 | 25 | 1838 | 155 |
| L196 | 84 | 1535 | 575 | 25 | 1489 | 124 | 25 | 1553 | 107 | 25 | 1494 | 127 |
| L197 | 84 | 1693 | 758 | 25 | 1642 | 460 | 25 | 1729 | 411 | 25 | 1678 | 420 |
| L198 | 84 | 1392 | 494 | 25 | 1393 | 88 | 25 | 1404 | 85 | 25 | 1402 | 93 |
| L199 | 82 | 1408 | 571 | 24 | 1426 | 182 | 24 | 1407 | 194 | 24 | 1408 | 201 |
| L2 | 80 | 1473 | 389 | 24 | 1461 | 95 | 24 | 1463 | 92 | 24 | 1453 | 88 |
| L20 | 84 | 1544 | 702 | 25 | 1541 | 263 | 25 | 1504 | 306 | 25 | 1476 | 281 |
| L200 | 84 | 1411 | 485 | 25 | 1472 | 86 | 25 | 1432 | 78 | 25 | 1471 | 77 |
| L201 | 84 | 1715 | 726 | 25 | 1682 | 257 | 25 | 1620 | 258 | 25 | 1668 | 253 |
| L202 | 84 | 3334 | 686 | 25 | 3315 | 281 | 25 | 3322 | 303 | 25 | 3240 | 291 |
| L203 | 80 | 1242 | 284 | 24 | 1245 | 56 | 24 | 1242 | 63 | 24 | 1242 | 57 |
| L204 | 82 | 1339 | 506 | 24 | 1342 | 113 | 24 | 1342 | 109 | 24 | 1339 | 97 |
| L207 | 84 | 1870 | 480 | 25 | 1983 | 98 | 25 | 1984 | 95 | 25 | 1976 | 102 |
| L21 | 84 | 1381 | 589 | 25 | 1321 | 228 | 25 | 1300 | 227 | 25 | 1311 | 200 |
| L210 | 84 | 1547 | 445 | 25 | 1519 | 138 | 25 | 1474 | 156 | 25 | 1528 | 97 |
| L212 | 82 | 1760 | 470 | 24 | 1834 | 83 | 24 | 1834 | 83 | 24 | 1834 | 87 |
| L213 | 32 | 1304 | 160 | 9 | 1304 | 17 | 9 | 1304 | 17 | 9 | 1304 | 17 |
| L217 | 84 | 1308 | 316 | 25 | 1308 | 74 | 25 | 1306 | 74 | 25 | 1306 | 72 |
| L219 | 84 | 1720 | 614 | 25 | 1721 | 184 | 25 | 1793 | 134 | 25 | 1784 | 135 |
| L223 | 84 | 1725 | 513 | 25 | 1722 | 213 | 25 | 1745 | 185 | 25 | 1722 | 211 |
| L226 | 84 | 1672 | 744 | 25 | 1662 | 358 | 25 | 1673 | 361 | 25 | 1752 | 352 |
| L228 | 84 | 1323 | 691 | 25 | 1446 | 264 | 25 | 1441 | 245 | 25 | 1377 | 297 |
| L229 | 76 | 1413 | 492 | 22 | 1413 | 97 | 22 | 1414 | 97 | 22 | 1493 | 124 |
| L23 | 84 | 1664 | 740 | 25 | 1562 | 458 | 25 | 1656 | 477 | 25 | 1679 | 426 |
| L230 | 84 | 1556 | 592 | 25 | 1584 | 149 | 25 | 1527 | 154 | 25 | 1697 | 156 |
| L232 | 84 | 1596 | 587 | 25 | 1603 | 132 | 25 | 1600 | 155 | 25 | 1665 | 154 |
| L234 | 84 | 1848 | 747 | 25 | 1743 | 324 | 25 | 1861 | 265 | 25 | 1820 | 275 |
| L235 | 84 | 1653 | 419 | 25 | 1657 | 103 | 25 | 1657 | 106 | 25 | 1657 | 113 |
| L236 | 84 | 1587 | 325 | 25 | 1558 | 100 | 25 | 1587 | 108 | 25 | 1568 | 95 |
| L238 | 84 | 1210 | 424 | 25 | 1202 | 191 | 25 | 1201 | 189 | 25 | 1201 | 188 |
| L24 | 84 | 1521 | 463 | 25 | 1555 | 147 | 25 | 1555 | 145 | 25 | 1523 | 143 |
| L240 | 84 | 2004 | 812 | 25 | 1991 | 230 | 25 | 2113 | 217 | 25 | 2022 | 316 |
| L241 | 84 | 1790 | 999 | 25 | 1817 | 511 | 25 | 1827 | 503 | 25 | 1825 | 537 |
| L242 | 84 | 1144 | 219 | 25 | 1144 | 60 | 25 | 1144 | 60 | 25 | 1144 | 59 |
| L243 | 84 | 1521 | 479 | 25 | 1636 | 78 | 25 | 1611 | 75 | 25 | 1521 | 74 |
| L244 | 84 | 1566 | 394 | 25 | 1564 | 82 | 25 | 1529 | 82 | 25 | 1564 | 85 |
| L245 | 84 | 1659 | 589 | 25 | 1660 | 111 | 25 | 1657 | 111 | 25 | 1657 | 127 |
| L246 | 84 | 1276 | 484 | 25 | 1245 | 116 | 25 | 1245 | 115 | 25 | 1293 | 120 |
| L248 | 84 | 2146 | 618 | 25 | 2152 | 104 | 25 | 2144 | 107 | 25 | 2257 | 122 |
| L249 | 84 | 1255 | 473 | 25 | 1281 | 92 | 25 | 1258 | 89 | 25 | 1281 | 97 |
| L25 | 84 | 1627 | 638 | 25 | 1675 | 258 | 25 | 1539 | 231 | 25 | 1544 | 251 |
| L250 | 84 | 1607 | 408 | 25 | 1607 | 108 | 25 | 1606 | 96 | 25 | 1607 | 106 |
| L251 | 18 | 447 | 71 | 4 | 447 | 2 | 4 | 447 | 2 | 4 | 447 | 2 |
| L252 | 84 | 2198 | 657 | 25 | 2209 | 125 | 25 | 2203 | 117 | 25 | 2198 | 129 |
| L253 | 84 | 1670 | 593 | 25 | 1682 | 196 | 25 | 1682 | 195 | 25 | 1670 | 206 |
| L254 | 84 | 1553 | 590 | 25 | 1551 | 160 | 25 | 1549 | 157 | 25 | 1551 | 167 |
| L255 | 84 | 1428 | 566 | 25 | 1406 | 124 | 25 | 1425 | 127 | 25 | 1485 | 114 |
| L256 | 84 | 1786 | 561 | 25 | 1704 | 242 | 25 | 1705 | 245 | 25 | 1715 | 208 |

| | | | | | | | | | | | | |
|---|---|---|---|---|---|---|---|---|---|---|---|---|
| L257 | 84 | 1709 | 595 | 25 | 1734 | 180 | 25 | 1642 | 174 | 25 | 1649 | 163 |
| L26 | 84 | 1506 | 413 | 25 | 1494 | 93 | 25 | 1494 | 88 | 25 | 1494 | 101 |
| L260 | 84 | 1487 | 342 | 25 | 1455 | 105 | 25 | 1455 | 88 | 25 | 1455 | 87 |
| L261 | 84 | 1365 | 446 | 25 | 1306 | 150 | 25 | 1306 | 148 | 25 | 1302 | 155 |
| L262 | 84 | 1313 | 262 | 25 | 1308 | 62 | 25 | 1308 | 61 | 25 | 1310 | 64 |
| L263 | 84 | 2905 | 1084 | 25 | 3102 | 405 | 25 | 3097 | 441 | 25 | 2968 | 400 |
| L265 | 82 | 1219 | 372 | 24 | 1219 | 70 | 24 | 1219 | 67 | 24 | 1218 | 92 |
| L266 | 84 | 1433 | 644 | 25 | 1413 | 260 | 25 | 1420 | 253 | 25 | 1384 | 278 |
| L268 | 84 | 2061 | 549 | 25 | 1983 | 90 | 25 | 1983 | 90 | 25 | 2057 | 112 |
| L270 | 84 | 2272 | 329 | 25 | 2285 | 145 | 25 | 2285 | 140 | 25 | 2271 | 177 |
| L273 | 80 | 1259 | 342 | 24 | 1259 | 85 | 24 | 1259 | 82 | 24 | 1259 | 95 |
| L274 | 84 | 1713 | 426 | 25 | 1713 | 81 | 25 | 1649 | 76 | 25 | 1653 | 98 |
| L275 | 84 | 1790 | 746 | 25 | 1777 | 120 | 25 | 1897 | 110 | 25 | 1790 | 142 |
| L277 | 84 | 1312 | 399 | 25 | 1362 | 76 | 25 | 1362 | 72 | 25 | 1361 | 71 |
| L278 | 84 | 1553 | 453 | 25 | 1551 | 84 | 25 | 1551 | 101 | 25 | 1606 | 82 |
| L279 | 84 | 1635 | 684 | 25 | 1614 | 320 | 25 | 1662 | 302 | 25 | 1628 | 291 |
| L28 | 84 | 1425 | 920 | 25 | 1417 | 491 | 25 | 1486 | 477 | 25 | 1434 | 516 |
| L280 | 84 | 1305 | 614 | 25 | 1281 | 219 | 25 | 1341 | 272 | 25 | 1297 | 276 |
| L281 | 84 | 1294 | 601 | 25 | 1234 | 331 | 25 | 1325 | 299 | 25 | 1313 | 307 |
| L282 | 84 | 1198 | 318 | 25 | 1211 | 61 | 25 | 1211 | 60 | 25 | 1211 | 62 |
| L283 | 84 | 1786 | 574 | 25 | 1773 | 144 | 25 | 1771 | 148 | 25 | 1746 | 141 |
| L284 | 84 | 1221 | 487 | 25 | 1168 | 112 | 25 | 1224 | 131 | 25 | 1217 | 155 |
| L285 | 78 | 1619 | 406 | 24 | 1600 | 125 | 24 | 1600 | 123 | 24 | 1601 | 127 |
| L286 | 84 | 1506 | 431 | 25 | 1510 | 84 | 25 | 1509 | 81 | 25 | 1505 | 76 |
| L287 | 84 | 1111 | 506 | 25 | 1078 | 308 | 25 | 1088 | 309 | 25 | 1233 | 202 |
| L288 | 84 | 1613 | 566 | 25 | 1604 | 237 | 25 | 1602 | 241 | 25 | 1596 | 220 |
| L289 | 84 | 1420 | 419 | 25 | 1411 | 106 | 25 | 1429 | 94 | 25 | 1420 | 84 |
| L29 | 84 | 1525 | 527 | 25 | 1490 | 142 | 25 | 1513 | 134 | 25 | 1533 | 114 |
| L291 | 84 | 1373 | 484 | 25 | 1356 | 170 | 25 | 1386 | 166 | 25 | 1386 | 186 |
| L292 | 84 | 1316 | 621 | 25 | 1322 | 233 | 25 | 1322 | 232 | 25 | 1299 | 213 |
| L293 | 84 | 1194 | 345 | 25 | 1155 | 169 | 25 | 1142 | 168 | 25 | 1218 | 158 |
| L294 | 84 | 2149 | 914 | 25 | 2042 | 523 | 25 | 2095 | 543 | 25 | 2039 | 544 |
| L295 | 84 | 1502 | 402 | 25 | 1508 | 92 | 25 | 1508 | 89 | 25 | 1508 | 116 |
| L296 | 84 | 1250 | 505 | 25 | 1256 | 76 | 25 | 1242 | 75 | 25 | 1274 | 86 |
| L297 | 82 | 1033 | 436 | 24 | 1138 | 104 | 24 | 1061 | 120 | 24 | 1033 | 124 |
| L298 | 84 | 1413 | 502 | 25 | 1322 | 121 | 25 | 1416 | 83 | 25 | 1408 | 112 |
| L299 | 84 | 1403 | 525 | 25 | 1468 | 152 | 25 | 1490 | 148 | 25 | 1395 | 144 |
| L3 | 84 | 1263 | 499 | 25 | 1256 | 177 | 25 | 1255 | 173 | 25 | 1263 | 172 |
| L300 | 84 | 1352 | 417 | 25 | 1326 | 97 | 25 | 1340 | 97 | 25 | 1334 | 97 |
| L302 | 84 | 1522 | 540 | 25 | 1582 | 141 | 25 | 1584 | 130 | 25 | 1556 | 147 |
| L305 | 84 | 1659 | 443 | 25 | 1658 | 81 | 25 | 1658 | 81 | 25 | 1659 | 82 |
| L307 | 80 | 1104 | 444 | 24 | 1091 | 118 | 24 | 1091 | 108 | 24 | 1091 | 119 |
| L308 | 84 | 1792 | 648 | 25 | 1838 | 220 | 25 | 1835 | 218 | 25 | 1745 | 243 |
| L309 | 84 | 1499 | 451 | 25 | 1443 | 213 | 25 | 1433 | 189 | 25 | 1444 | 208 |
| L31 | 84 | 1635 | 536 | 25 | 1738 | 105 | 25 | 1645 | 107 | 25 | 1628 | 111 |
| L310 | 84 | 1499 | 339 | 25 | 1389 | 110 | 25 | 1389 | 104 | 25 | 1389 | 114 |
| L311 | 80 | 1427 | 492 | 24 | 1508 | 118 | 24 | 1508 | 118 | 24 | 1415 | 99 |
| L312 | 84 | 1451 | 425 | 25 | 1444 | 127 | 25 | 1444 | 131 | 25 | 1444 | 132 |
| L313 | 84 | 1240 | 310 | 25 | 1236 | 89 | 25 | 1235 | 121 | 25 | 1238 | 92 |
| L314 | 84 | 1250 | 489 | 25 | 1235 | 124 | 25 | 1236 | 122 | 25 | 1226 | 136 |
| L315 | 84 | 1534 | 461 | 25 | 1563 | 164 | 25 | 1477 | 154 | 25 | 1546 | 140 |
| L316 | 84 | 1720 | 693 | 25 | 1720 | 193 | 25 | 1734 | 176 | 25 | 1726 | 205 |
| L318 | 84 | 1364 | 442 | 25 | 1411 | 111 | 25 | 1411 | 112 | 25 | 1363 | 114 |
| L319 | 84 | 1275 | 472 | 25 | 1291 | 90 | 25 | 1291 | 90 | 25 | 1234 | 114 |
| L320 | 84 | 1583 | 816 | 25 | 1610 | 271 | 25 | 1633 | 263 | 25 | 1636 | 304 |
| L321 | 84 | 1518 | 635 | 25 | 1606 | 188 | 25 | 1519 | 148 | 25 | 1598 | 162 |
| L322 | 84 | 1593 | 682 | 25 | 1702 | 231 | 25 | 1585 | 193 | 25 | 1572 | 177 |
| L323 | 80 | 1501 | 528 | 24 | 1496 | 130 | 24 | 1432 | 126 | 24 | 1432 | 124 |
| L324 | 84 | 1418 | 550 | 25 | 1338 | 236 | 25 | 1451 | 177 | 25 | 1396 | 238 |
| L325 | 84 | 1633 | 494 | 25 | 1616 | 85 | 25 | 1616 | 89 | 25 | 1629 | 88 |
| L326 | 84 | 1588 | 421 | 25 | 1560 | 188 | 25 | 1560 | 195 | 25 | 1556 | 153 |
| L327 | 80 | 1310 | 357 | 24 | 1179 | 125 | 24 | 1225 | 81 | 24 | 1189 | 105 |
| L328 | 84 | 1524 | 592 | 25 | 1486 | 139 | 25 | 1481 | 117 | 25 | 1486 | 121 |
| L329 | 84 | 1720 | 806 | 25 | 1826 | 225 | 25 | 1662 | 229 | 25 | 1866 | 222 |

| | | | | | | | | | | | | |
|---|---|---|---|---|---|---|---|---|---|---|---|---|
| L33 | 84 | 1734 | 695 | 25 | 1606 | 269 | 25 | 1882 | 205 | 25 | 1725 | 245 |
| L332 | 84 | 1265 | 569 | 25 | 1260 | 220 | 25 | 1281 | 196 | 25 | 1270 | 255 |
| L333 | 84 | 2601 | 1155 | 25 | 2492 | 395 | 25 | 2671 | 375 | 25 | 2654 | 376 |
| L334 | 84 | 1702 | 668 | 25 | 1721 | 211 | 25 | 1705 | 190 | 25 | 1839 | 186 |
| L336 | 84 | 1233 | 368 | 25 | 1285 | 120 | 25 | 1256 | 137 | 25 | 1254 | 138 |
| L337 | 84 | 1123 | 306 | 25 | 1123 | 77 | 25 | 1123 | 77 | 25 | 1123 | 66 |
| L338 | 84 | 1512 | 629 | 25 | 1457 | 218 | 25 | 1460 | 227 | 25 | 1503 | 205 |
| L339 | 84 | 1440 | 538 | 25 | 1411 | 161 | 25 | 1410 | 169 | 25 | 1350 | 167 |
| L34 | 84 | 1551 | 472 | 25 | 1559 | 65 | 25 | 1551 | 60 | 25 | 1459 | 71 |
| L340 | 84 | 2088 | 1012 | 25 | 2102 | 350 | 25 | 2053 | 368 | 25 | 2202 | 320 |
| L341 | 84 | 1303 | 757 | 25 | 1325 | 461 | 25 | 1349 | 425 | 25 | 1334 | 461 |
| L344 | 84 | 2210 | 748 | 25 | 2433 | 231 | 25 | 2261 | 196 | 25 | 2358 | 208 |
| L346 | 84 | 1772 | 472 | 25 | 1726 | 175 | 25 | 1714 | 175 | 25 | 1711 | 204 |
| L348 | 84 | 1760 | 594 | 25 | 1915 | 113 | 25 | 1898 | 98 | 25 | 1729 | 177 |
| L349 | 84 | 1499 | 563 | 25 | 1498 | 88 | 25 | 1498 | 83 | 25 | 1498 | 85 |
| L35 | 84 | 1797 | 866 | 25 | 1891 | 409 | 25 | 1901 | 410 | 25 | 1859 | 428 |
| L350 | 84 | 1668 | 455 | 25 | 1665 | 115 | 25 | 1665 | 101 | 25 | 1666 | 115 |
| L352 | 84 | 1485 | 444 | 25 | 1503 | 201 | 25 | 1607 | 181 | 25 | 1484 | 202 |
| L353 | 84 | 2468 | 705 | 25 | 2434 | 169 | 25 | 2465 | 152 | 25 | 2431 | 181 |
| L354 | 84 | 1683 | 736 | 25 | 1679 | 298 | 25 | 1726 | 301 | 25 | 1679 | 280 |
| L356 | 84 | 1509 | 660 | 25 | 1532 | 138 | 25 | 1531 | 145 | 25 | 1535 | 159 |
| L358 | 84 | 1901 | 459 | 25 | 1924 | 148 | 25 | 1883 | 72 | 25 | 1864 | 150 |
| L365 | 84 | 1361 | 743 | 25 | 1289 | 377 | 25 | 1364 | 343 | 25 | 1271 | 388 |
| L366 | 84 | 1613 | 341 | 25 | 1620 | 121 | 25 | 1620 | 127 | 25 | 1612 | 127 |
| L367 | 84 | 1548 | 637 | 25 | 1506 | 227 | 25 | 1525 | 213 | 25 | 1567 | 204 |
| L368 | 84 | 1320 | 447 | 25 | 1320 | 72 | 25 | 1315 | 72 | 25 | 1320 | 73 |
| L369 | 84 | 1448 | 599 | 25 | 1440 | 198 | 25 | 1440 | 154 | 25 | 1448 | 199 |
| L37 | 84 | 960 | 314 | 25 | 971 | 77 | 25 | 969 | 70 | 25 | 971 | 72 |
| L370 | 84 | 1382 | 519 | 25 | 1361 | 120 | 25 | 1384 | 86 | 25 | 1382 | 124 |
| L372 | 84 | 1594 | 500 | 25 | 1714 | 127 | 25 | 1620 | 120 | 25 | 1685 | 102 |
| L373 | 84 | 1776 | 594 | 25 | 1867 | 109 | 25 | 1866 | 108 | 25 | 1866 | 110 |
| L374 | 84 | 1497 | 755 | 25 | 1481 | 425 | 25 | 1530 | 399 | 25 | 1475 | 396 |
| L375 | 84 | 1580 | 648 | 25 | 1573 | 134 | 25 | 1576 | 127 | 25 | 1579 | 154 |
| L376 | 84 | 1256 | 286 | 25 | 1253 | 58 | 25 | 1255 | 58 | 25 | 1252 | 60 |
| L379 | 84 | 2156 | 709 | 25 | 2004 | 314 | 25 | 1969 | 318 | 25 | 1979 | 317 |
| L38 | 84 | 1333 | 364 | 25 | 1332 | 116 | 25 | 1332 | 106 | 25 | 1333 | 120 |
| L380 | 84 | 1532 | 649 | 25 | 1605 | 196 | 25 | 1605 | 193 | 25 | 1605 | 201 |
| L381 | 84 | 1373 | 444 | 25 | 1377 | 104 | 25 | 1377 | 103 | 25 | 1377 | 103 |
| L382 | 84 | 1853 | 668 | 25 | 1867 | 201 | 25 | 1850 | 236 | 25 | 1832 | 172 |
| L384 | 84 | 1422 | 302 | 25 | 1410 | 82 | 25 | 1407 | 80 | 25 | 1410 | 81 |
| L386 | 84 | 1810 | 723 | 25 | 1895 | 333 | 25 | 1820 | 322 | 25 | 1865 | 330 |
| L387 | 84 | 1814 | 237 | 25 | 1811 | 51 | 25 | 1814 | 54 | 25 | 1814 | 61 |
| L39 | 84 | 1439 | 473 | 25 | 1387 | 107 | 25 | 1387 | 100 | 25 | 1389 | 117 |
| L390 | 84 | 1247 | 522 | 25 | 1281 | 138 | 25 | 1217 | 157 | 25 | 1219 | 148 |
| L391 | 84 | 2025 | 658 | 25 | 2291 | 103 | 25 | 2291 | 101 | 25 | 1999 | 207 |
| L392 | 84 | 1431 | 537 | 25 | 1434 | 141 | 25 | 1418 | 141 | 25 | 1417 | 200 |
| L394 | 84 | 1640 | 646 | 25 | 1630 | 272 | 25 | 1630 | 264 | 25 | 1643 | 270 |
| L395 | 84 | 1912 | 480 | 25 | 1842 | 121 | 25 | 1877 | 114 | 25 | 1857 | 132 |
| L396 | 84 | 1487 | 619 | 25 | 1519 | 192 | 25 | 1487 | 203 | 25 | 1598 | 176 |
| L397 | 84 | 2080 | 616 | 25 | 1949 | 189 | 25 | 1949 | 188 | 25 | 1877 | 224 |
| L398 | 84 | 1280 | 437 | 25 | 1258 | 128 | 25 | 1258 | 122 | 25 | 1256 | 146 |
| L4 | 84 | 1467 | 753 | 25 | 1520 | 202 | 25 | 1580 | 196 | 25 | 1495 | 184 |
| L400 | 84 | 1644 | 591 | 25 | 1566 | 280 | 25 | 1584 | 282 | 25 | 1618 | 283 |
| L401 | 80 | 1423 | 413 | 24 | 1431 | 94 | 24 | 1430 | 94 | 24 | 1394 | 103 |
| L402 | 84 | 1340 | 598 | 25 | 1324 | 168 | 25 | 1304 | 189 | 25 | 1340 | 152 |
| L403 | 84 | 1193 | 350 | 25 | 1193 | 69 | 25 | 1206 | 70 | 25 | 1236 | 71 |
| L404 | 84 | 1755 | 565 | 25 | 1572 | 162 | 25 | 1589 | 156 | 25 | 1572 | 148 |
| L405 | 84 | 1380 | 408 | 25 | 1289 | 108 | 25 | 1363 | 103 | 25 | 1289 | 106 |
| L406 | 84 | 1605 | 654 | 25 | 1606 | 229 | 25 | 1617 | 234 | 25 | 1524 | 262 |
| L407 | 84 | 2242 | 956 | 25 | 2090 | 492 | 25 | 2012 | 462 | 25 | 2013 | 492 |
| L408 | 84 | 1767 | 463 | 25 | 1772 | 109 | 25 | 1650 | 121 | 25 | 1762 | 108 |
| L41 | 84 | 1304 | 245 | 25 | 1304 | 51 | 25 | 1304 | 51 | 25 | 1304 | 54 |
| L410 | 84 | 1426 | 728 | 25 | 1468 | 375 | 25 | 1434 | 372 | 25 | 1435 | 361 |
| L411 | 78 | 1537 | 327 | 23 | 1485 | 61 | 23 | 1485 | 59 | 23 | 1485 | 65 |

| | | | | | | | | | | | | |
|---|---|---|---|---|---|---|---|---|---|---|---|---|
| L412 | 84 | 1755 | 615 | 25 | 1672 | 200 | 25 | 1727 | 210 | 25 | 1690 | 186 |
| L413 | 70 | 1294 | 225 | 20 | 1294 | 40 | 20 | 1294 | 36 | 20 | 1293 | 51 |
| L414 | 84 | 1318 | 311 | 25 | 1261 | 89 | 25 | 1276 | 72 | 25 | 1313 | 114 |
| L415 | 84 | 1484 | 338 | 25 | 1482 | 76 | 25 | 1484 | 66 | 25 | 1484 | 80 |
| L416 | 84 | 1567 | 754 | 25 | 1570 | 155 | 25 | 1570 | 153 | 25 | 1571 | 157 |
| L418 | 84 | 2022 | 752 | 25 | 1927 | 288 | 25 | 1983 | 258 | 25 | 1966 | 290 |
| L419 | 84 | 1373 | 310 | 25 | 1366 | 46 | 25 | 1366 | 44 | 25 | 1365 | 58 |
| L420 | 84 | 1583 | 547 | 25 | 1575 | 105 | 25 | 1539 | 115 | 25 | 1517 | 128 |
| L421 | 84 | 1704 | 710 | 25 | 1707 | 187 | 25 | 1644 | 207 | 25 | 1660 | 225 |
| L422 | 84 | 1355 | 243 | 25 | 1356 | 74 | 25 | 1349 | 71 | 25 | 1355 | 74 |
| L423 | 80 | 1255 | 467 | 24 | 1255 | 105 | 24 | 1255 | 104 | 24 | 1255 | 106 |
| L424 | 84 | 1692 | 468 | 25 | 1692 | 118 | 25 | 1687 | 111 | 25 | 1692 | 123 |
| L425 | 84 | 1380 | 366 | 25 | 1387 | 87 | 25 | 1387 | 86 | 25 | 1388 | 88 |
| L426 | 84 | 1887 | 435 | 25 | 1850 | 131 | 25 | 1876 | 110 | 25 | 1848 | 119 |
| L428 | 80 | 1241 | 434 | 24 | 1199 | 185 | 24 | 1155 | 176 | 24 | 1208 | 171 |
| L429 | 84 | 1413 | 519 | 25 | 1475 | 189 | 25 | 1456 | 203 | 25 | 1468 | 194 |
| L430 | 84 | 1466 | 441 | 25 | 1466 | 83 | 25 | 1455 | 86 | 25 | 1455 | 85 |
| L431 | 76 | 1372 | 385 | 22 | 1409 | 113 | 22 | 1427 | 124 | 22 | 1410 | 120 |
| L432 | 84 | 1543 | 454 | 25 | 1571 | 84 | 25 | 1707 | 83 | 25 | 1704 | 75 |
| L433 | 84 | 1451 | 501 | 25 | 1446 | 124 | 25 | 1445 | 118 | 25 | 1465 | 108 |
| L434 | 84 | 1519 | 472 | 25 | 1451 | 93 | 25 | 1505 | 91 | 25 | 1507 | 95 |
| L435 | 84 | 3085 | 907 | 25 | 3068 | 336 | 25 | 3090 | 312 | 25 | 3116 | 324 |
| L436 | 76 | 1569 | 475 | 21 | 1621 | 81 | 21 | 1646 | 93 | 21 | 1635 | 93 |
| L437 | 84 | 1366 | 375 | 25 | 1347 | 112 | 25 | 1347 | 112 | 25 | 1348 | 110 |
| L438 | 84 | 1696 | 666 | 25 | 1619 | 359 | 25 | 1512 | 334 | 25 | 1598 | 355 |
| L439 | 84 | 1358 | 400 | 25 | 1372 | 87 | 25 | 1371 | 87 | 25 | 1389 | 85 |
| L440 | 84 | 1369 | 477 | 25 | 1368 | 125 | 25 | 1368 | 120 | 25 | 1380 | 132 |
| L441 | 84 | 1454 | 528 | 25 | 1427 | 132 | 25 | 1444 | 136 | 25 | 1433 | 140 |
| L442 | 82 | 1595 | 498 | 24 | 1584 | 122 | 24 | 1584 | 124 | 24 | 1584 | 128 |
| L443 | 84 | 1413 | 514 | 25 | 1407 | 194 | 25 | 1359 | 191 | 25 | 1363 | 180 |
| L447 | 84 | 2310 | 787 | 25 | 2527 | 197 | 25 | 2527 | 197 | 25 | 2578 | 176 |
| L448 | 84 | 1709 | 621 | 25 | 1694 | 199 | 25 | 1610 | 198 | 25 | 1743 | 198 |
| L449 | 74 | 932 | 142 | 22 | 932 | 78 | 22 | 932 | 79 | 22 | 927 | 98 |
| L450 | 84 | 1490 | 453 | 25 | 1490 | 153 | 25 | 1550 | 126 | 25 | 1492 | 151 |
| L46 | 84 | 1900 | 274 | 25 | 1900 | 55 | 25 | 1900 | 55 | 25 | 1900 | 53 |
| L48 | 84 | 1567 | 568 | 25 | 1496 | 158 | 25 | 1499 | 175 | 25 | 1579 | 152 |
| L49 | 84 | 1917 | 539 | 25 | 1823 | 255 | 25 | 1667 | 237 | 25 | 1879 | 233 |
| L5 | 84 | 1058 | 417 | 25 | 1045 | 107 | 25 | 1045 | 106 | 25 | 1058 | 105 |
| L50 | 84 | 1526 | 600 | 25 | 1733 | 136 | 25 | 1692 | 133 | 25 | 1692 | 121 |
| L51 | 84 | 1224 | 338 | 25 | 1221 | 49 | 25 | 1226 | 50 | 25 | 1229 | 56 |
| L53 | 84 | 1643 | 348 | 25 | 1643 | 69 | 25 | 1643 | 69 | 25 | 1643 | 75 |
| L54 | 84 | 1754 | 463 | 25 | 1711 | 79 | 25 | 1711 | 80 | 25 | 1723 | 96 |
| L55 | 84 | 1959 | 331 | 25 | 1959 | 47 | 25 | 1959 | 43 | 25 | 1959 | 50 |
| L56 | 84 | 1519 | 578 | 25 | 1533 | 178 | 25 | 1533 | 173 | 25 | 1545 | 227 |
| L57 | 84 | 1901 | 694 | 25 | 1948 | 215 | 25 | 1916 | 214 | 25 | 1843 | 222 |
| L58 | 84 | 1796 | 511 | 25 | 1732 | 140 | 25 | 1798 | 144 | 25 | 1734 | 125 |
| L59 | 84 | 2069 | 772 | 25 | 1984 | 176 | 25 | 1966 | 168 | 25 | 2071 | 149 |
| L6 | 84 | 1684 | 435 | 25 | 1723 | 97 | 25 | 1724 | 97 | 25 | 1724 | 108 |
| L60 | 84 | 1682 | 494 | 25 | 1681 | 89 | 25 | 1681 | 86 | 25 | 1669 | 113 |
| L61 | 84 | 1667 | 536 | 25 | 1802 | 145 | 25 | 1803 | 144 | 25 | 1802 | 147 |
| L64 | 84 | 1239 | 397 | 25 | 1239 | 101 | 25 | 1239 | 97 | 25 | 1237 | 93 |
| L67 | 84 | 1564 | 425 | 25 | 1570 | 172 | 25 | 1566 | 166 | 25 | 1566 | 183 |
| L69 | 84 | 1536 | 406 | 25 | 1589 | 120 | 25 | 1535 | 113 | 25 | 1591 | 120 |
| L7 | 84 | 2262 | 727 | 25 | 2333 | 156 | 25 | 2126 | 146 | 25 | 2323 | 133 |
| L70 | 84 | 1644 | 639 | 25 | 1681 | 283 | 25 | 1592 | 259 | 25 | 1627 | 253 |
| L71 | 84 | 1464 | 694 | 25 | 1439 | 415 | 25 | 1432 | 405 | 25 | 1464 | 377 |
| L72 | 84 | 1714 | 458 | 25 | 1594 | 307 | 25 | 1575 | 307 | 25 | 1545 | 305 |
| L73 | 84 | 1402 | 469 | 25 | 1441 | 185 | 25 | 1437 | 172 | 25 | 1435 | 199 |
| L74 | 84 | 1383 | 716 | 25 | 1239 | 447 | 25 | 1293 | 435 | 25 | 1315 | 434 |
| L75 | 84 | 1713 | 597 | 25 | 1713 | 118 | 25 | 1714 | 134 | 25 | 1651 | 142 |
| L77 | 84 | 1469 | 595 | 25 | 1345 | 194 | 25 | 1345 | 186 | 25 | 1481 | 185 |
| L78 | 84 | 1690 | 994 | 25 | 1828 | 462 | 25 | 2038 | 469 | 25 | 1743 | 499 |
| L79 | 84 | 1785 | 726 | 25 | 1879 | 295 | 25 | 1757 | 340 | 25 | 1790 | 361 |
| L8 | 84 | 2576 | 645 | 25 | 2566 | 234 | 25 | 2566 | 218 | 25 | 2670 | 257 |

| | | | | | | | | | | | | |
|---|---|---|---|---|---|---|---|---|---|---|---|---|
| L80 | 84 | 1492 | 538 | 25 | 1500 | 245 | 25 | 1589 | 200 | 25 | 1532 | 202 |
| L83 | 84 | 2100 | 692 | 25 | 2123 | 262 | 25 | 2125 | 254 | 25 | 2068 | 284 |
| L84 | 84 | 1432 | 600 | 25 | 1441 | 109 | 25 | 1441 | 113 | 25 | 1582 | 120 |
| L85 | 84 | 1124 | 415 | 25 | 1146 | 134 | 25 | 1149 | 150 | 25 | 1148 | 137 |
| L86 | 84 | 1268 | 383 | 25 | 1278 | 81 | 25 | 1278 | 80 | 25 | 1281 | 79 |
| L87 | 84 | 1614 | 660 | 25 | 1615 | 190 | 25 | 1635 | 171 | 25 | 1596 | 200 |
| L89 | 84 | 1226 | 264 | 25 | 1220 | 61 | 25 | 1220 | 60 | 25 | 1226 | 67 |
| L9 | 84 | 1823 | 776 | 25 | 1849 | 187 | 25 | 1849 | 176 | 25 | 1884 | 197 |
| L91 | 84 | 1571 | 283 | 25 | 1561 | 54 | 25 | 1561 | 54 | 25 | 1561 | 54 |
| L93 | 84 | 1593 | 366 | 25 | 1628 | 169 | 25 | 1628 | 168 | 25 | 1532 | 207 |
| L94 | 78 | 982 | 379 | 23 | 1007 | 207 | 23 | 971 | 226 | 23 | 1033 | 222 |
| L95 | 84 | 1395 | 298 | 25 | 1312 | 164 | 25 | 1312 | 165 | 25 | 1343 | 147 |
| L97 | 84 | 1119 | 486 | 25 | 1124 | 161 | 25 | 1124 | 140 | 25 | 1123 | 174 |
| L98 | 84 | 2130 | 954 | 25 | 2131 | 557 | 25 | 2160 | 499 | 25 | 2201 | 526 |
| L99 | 84 | 1684 | 721 | 25 | 1646 | 245 | 25 | 1618 | 237 | 25 | 1707 | 246 |
| **Average** | **83.02** | **1564.24** | **530.65** | **24.69** | **1561.96** | **168.08** | **24.69** | **1562.20** | **163.76** | **24.69** | **1563.16** | **168.92** |

**Table S3 – Alignment Statistics across Data Types.** PI sites are the number of Parsimony Informative sites.

| Allopolyploid | Data Type | Parameter | Mean | Lower 95% HPD | Upper 95% HPD |
|---|---|---|---|---|---|
| *D. celsa* | phased | lnL | -679943.918 | -680077.843 | -679805.337 |
| *D. celsa* | genotype | lnL | -643104.864 | -643205.417 | -643004.169 |
| *D. celsa* | consensus | lnL | -627201.4897 | -627284.005 | -627119.895 |
| *D. celsa* | pickone | lnL | -634366.8596 | -634447.562 | -634285.619 |
| *D. celsa* | phased | phi_h<-t | 0.504627 | 0.436357 | 0.6569 |
| *D. celsa* | genotype | phi_h<-t | 0.428202 | 0.38779 | 0.466869 |
| *D. celsa* | consensus | phi_h<-t | 0.45668 | 0.413564 | 0.498812 |
| *D. celsa* | pickone | phi_h<-t | 0.454166 | 0.404159 | 0.503595 |
| *D. celsa* | phased | tau_4r | 0.005085 | 0.002027 | 0.006392 |
| *D. celsa* | genotype | tau_4r | 0.006077 | 0.005563 | 0.006601 |
| *D. celsa* | consensus | tau_4r | 0.005468 | 0.004957 | 0.005985 |
| *D. celsa* | pickone | tau_4r | 0.006045 | 0.005425 | 0.006716 |
| *D. celsa* | phased | tau_5t | 0.000414 | 0.000306 | 0.000508 |
| *D. celsa* | genotype | tau_5t | 0.000438 | 0.000311 | 0.000565 |
| *D. celsa* | consensus | tau_5t | 0.000584 | 0.00029 | 0.000839 |
| *D. celsa* | pickone | tau_5t | 0.000797 | 0.000533 | 0.001021 |
| *D. celsa* | phased | tau_6s | 0.000549 | 0.000449 | 0.000654 |
| *D. celsa* | genotype | tau_6s | 0.000628 | 0.000515 | 0.000752 |
| *D. celsa* | consensus | tau_6s | 0.000599 | 0.000467 | 0.00073 |
| *D. celsa* | pickone | tau_6s | 0.00086 | 0.000684 | 0.001035 |
| *D. celsa* | phased | tau_7h | 0.000393 | 0.000272 | 0.000502 |
| *D. celsa* | genotype | tau_7h | 0.000223 | 0.000141 | 0.000318 |
| *D. celsa* | consensus | tau_7h | 0.000428 | 0.000001 | 0.000652 |
| *D. celsa* | pickone | tau_7h | 0.00076 | 0.000496 | 0.001001 |
| *D. celsa* | phased | theta_1Dgol | 0.000457 | 0.000349 | 0.000557 |
| *D. celsa* | genotype | theta_1Dgol | 0.000523 | 0.000398 | 0.00065 |
| *D. celsa* | consensus | theta_1Dgol | NA | NA | NA |
| *D. celsa* | pickone | theta_1Dgol | NA | NA | NA |
| *D. celsa* | phased | theta_2Dlud | 0.001426 | 0.001173 | 0.001692 |
| *D. celsa* | genotype | theta_2Dlud | 0.001778 | 0.001502 | 0.002049 |
| *D. celsa* | consensus | theta_2Dlud | 0.002253 | 0.001694 | 0.002852 |
| *D. celsa* | pickone | theta_2Dlud | 0.002607 | 0.002004 | 0.003215 |
| *D. celsa* | phased | theta_3Dcel | 0.06869 | 0.039693 | 0.110449 |
| *D. celsa* | genotype | theta_3Dcel | 0.001671 | 0.00115 | 0.002215 |
| *D. celsa* | consensus | theta_3Dcel | 0.082217 | 0.000998 | 0.238998 |
| *D. celsa* | pickone | theta_3Dcel | 0.280123 | 0.059426 | 0.655403 |
| *D. celsa* | phased | theta_4r | 0.034482 | 0.02827 | 0.038914 |
| *D. celsa* | genotype | theta_4r | 0.046163 | 0.041346 | 0.051188 |
| *D. celsa* | consensus | theta_4r | 0.047823 | 0.042752 | 0.053267 |
| *D. celsa* | pickone | theta_4r | 0.045486 | 0.040775 | 0.050331 |
| *D. celsa* | phased | theta_5t | 0.014404 | 0.010708 | 0.023365 |
| *D. celsa* | genotype | theta_5t | 0.009078 | 0.007762 | 0.010407 |
| *D. celsa* | consensus | theta_5t | 0.008391 | 0.006775 | 0.010079 |
| *D. celsa* | pickone | theta_5t | 0.009492 | 0.007309 | 0.011678 |
| *D. celsa* | phased | theta_6s | 0.00937 | 0.004944 | 0.011689 |
| *D. celsa* | genotype | theta_6s | 0.008004 | 0.007046 | 0.008927 |
| *D. celsa* | consensus | theta_6s | 0.005645 | 0.004806 | 0.006514 |
| *D. celsa* | pickone | theta_6s | 0.007863 | 0.006482 | 0.009268 |

| | | | | | |
|---|---|---|---|---|---|
| *D. celsa* | phased | theta_7h | 0.006979 | 0.00102 | 0.01363 |
| *D. celsa* | genotype | theta_7h | 0.013604 | 0.003842 | 0.030768 |
| *D. celsa* | consensus | theta_7h | 0.018192 | 0.000882 | 0.061998 |
| *D. celsa* | pickone | theta_7h | 0.01308 | 0.00098 | 0.040084 |
| *D. celsa* | phased | theta_8h | 0.009827 | 0.00103 | 0.028194 |
| *D. celsa* | genotype | theta_8h | 0.008762 | 0.001611 | 0.021348 |
| *D. celsa* | consensus | theta_8h | 0.011238 | 0.001026 | 0.032819 |
| *D. celsa* | pickone | theta_8h | 0.008789 | 0.000953 | 0.024666 |
| *D. campyloptera* | phased | lnL | -585944.0596 | -586075.806 | -585813.001 |
| *D. campyloptera* | genotype | lnL | -569990.0378 | -570099.939 | -569876.18 |
| *D. campyloptera* | consensus | lnL | -545586.866 | -545676.096 | -545500.869 |
| *D. campyloptera* | pickone | lnL | -551530.8014 | -551618.271 | -551439.823 |
| *D. campyloptera* | phased | phi_h<-t | 0.471758 | 0.395516 | 0.550255 |
| *D. campyloptera* | genotype | phi_h<-t | 0.522176 | 0.463008 | 0.582006 |
| *D. campyloptera* | consensus | phi_h<-t | 0.544687 | 0.483768 | 0.603035 |
| *D. campyloptera* | pickone | phi_h<-t | 0.555426 | 0.495655 | 0.612902 |
| *D. campyloptera* | phased | tau_4r | 0.007279 | 0.006759 | 0.007789 |
| *D. campyloptera* | genotype | tau_4r | 0.007384 | 0.006867 | 0.007921 |
| *D. campyloptera* | consensus | tau_4r | 0.005939 | 0.005381 | 0.00647 |
| *D. campyloptera* | pickone | tau_4r | 0.006621 | 0.006051 | 0.007211 |
| *D. campyloptera* | phased | tau_5t | 0.001513 | 0.001243 | 0.0018 |
| *D. campyloptera* | genotype | tau_5t | 0.000882 | 0.000671 | 0.001099 |
| *D. campyloptera* | consensus | tau_5t | 0.000658 | 0.000389 | 0.000948 |
| *D. campyloptera* | pickone | tau_5t | 0.000835 | 0.000461 | 0.00121 |
| *D. campyloptera* | phased | tau_6s | 0.002178 | 0.001886 | 0.002479 |
| *D. campyloptera* | genotype | tau_6s | 0.002077 | 0.001726 | 0.002428 |
| *D. campyloptera* | consensus | tau_6s | 0.001621 | 0.001298 | 0.001959 |
| *D. campyloptera* | pickone | tau_6s | 0.00197 | 0.001608 | 0.002339 |
| *D. campyloptera* | phased | tau_7h | 0.001485 | 0.001204 | 0.001766 |
| *D. campyloptera* | genotype | tau_7h | 0.000472 | 0.000297 | 0.000641 |
| *D. campyloptera* | consensus | tau_7h | 0.000371 | 0.000004 | 0.000795 |
| *D. campyloptera* | pickone | tau_7h | 0.000569 | 0.000002 | 0.001084 |
| *D. campyloptera* | phased | theta_1Dexp | 0.005722 | 0.00492 | 0.006574 |
| *D. campyloptera* | genotype | theta_1Dexp | 0.004458 | 0.003668 | 0.005297 |
| *D. campyloptera* | consensus | theta_1Dexp | 0.004865 | 0.003196 | 0.006632 |
| *D. campyloptera* | pickone | theta_1Dexp | 0.006904 | 0.004535 | 0.00949 |
| *D. campyloptera* | phased | theta_2Dint | 0.005479 | 0.004995 | 0.006006 |
| *D. campyloptera* | genotype | theta_2Dint | 0.006216 | 0.005602 | 0.00682 |
| *D. campyloptera* | consensus | theta_2Dint | 0.005006 | 0.004248 | 0.005769 |
| *D. campyloptera* | pickone | theta_2Dint | 0.006481 | 0.005484 | 0.007474 |
| *D. campyloptera* | phased | theta_3Dcam | 0.014223 | 0.012595 | 0.015867 |
| *D. campyloptera* | genotype | theta_3Dcam | 0.00149 | 0.001085 | 0.001913 |
| *D. campyloptera* | consensus | theta_3Dcam | 0.02878 | 0.001191 | 0.077795 |
| *D. campyloptera* | pickone | theta_3Dcam | 0.064045 | 0.001128 | 0.18099 |
| *D. campyloptera* | phased | theta_4r | 0.094151 | 0.086343 | 0.102337 |
| *D. campyloptera* | genotype | theta_4r | 0.097957 | 0.089609 | 0.106504 |
| *D. campyloptera* | consensus | theta_4r | 0.100146 | 0.091742 | 0.109043 |
| *D. campyloptera* | pickone | theta_4r | 0.094327 | 0.08638 | 0.102448 |
| *D. campyloptera* | phased | theta_5t | 0.024663 | 0.021883 | 0.027395 |

| | | | | | |
|---|---|---|---|---|---|
| *D. campyloptera* | genotype | theta_5t | 0.028104 | 0.024631 | 0.031627 |
| *D. campyloptera* | consensus | theta_5t | 0.025404 | 0.021655 | 0.029276 |
| *D. campyloptera* | pickone | theta_5t | 0.032728 | 0.027807 | 0.037801 |
| *D. campyloptera* | phased | theta_6s | 0.009647 | 0.008377 | 0.01091 |
| *D. campyloptera* | genotype | theta_6s | 0.00871 | 0.007435 | 0.009976 |
| *D. campyloptera* | consensus | theta_6s | 0.006315 | 0.005204 | 0.007401 |
| *D. campyloptera* | pickone | theta_6s | 0.007617 | 0.006301 | 0.008929 |
| *D. campyloptera* | phased | theta_7h | 0.008955 | 0.003843 | 0.015733 |
| *D. campyloptera* | genotype | theta_7h | 0.01218 | 0.003548 | 0.025771 |
| *D. campyloptera* | consensus | theta_7h | 0.010725 | 0.002701 | 0.02266 |
| *D. campyloptera* | pickone | theta_7h | 0.020217 | 0.003158 | 0.047095 |
| *D. campyloptera* | phased | theta_8h | 0.003633 | 0.000684 | 0.009084 |
| *D. campyloptera* | genotype | theta_8h | 0.008383 | 0.001784 | 0.019821 |
| *D. campyloptera* | consensus | theta_8h | 0.014371 | 0.001025 | 0.040727 |
| *D. campyloptera* | pickone | theta_8h | 0.016487 | 0.000951 | 0.055001 |
| *D. clintoniana* | phased | lnL | -844606.6632 | -844796.987 | -844414.86 |
| *D. clintoniana* | genotype | lnL | -691785.3795 | -691893.399 | -691677.283 |
| *D. clintoniana* | consensus | lnL | -678628.4082 | -678713.278 | -678537.146 |
| *D. clintoniana* | pickone | lnL | -692474.636 | -692564.019 | -692383.155 |
| *D. clintoniana* | phased | phi_h<-t | 0.332922 | 0.106005 | 0.840279 |
| *D. clintoniana* | genotype | phi_h<-t | 0.254936 | 0.189996 | 0.320312 |
| *D. clintoniana* | consensus | phi_h<-t | 0.757824 | 0.584269 | 0.912797 |
| *D. clintoniana* | pickone | phi_h<-t | 0.86745 | 0.795465 | 0.925204 |
| *D. clintoniana* | phased | tau_4r | 0.002857 | 0.002272 | 0.003331 |
| *D. clintoniana* | genotype | tau_4r | 0.002318 | 0.001958 | 0.002679 |
| *D. clintoniana* | consensus | tau_4r | 0.001409 | 0.001057 | 0.001781 |
| *D. clintoniana* | pickone | tau_4r | 0.002236 | 0.001715 | 0.002714 |
| *D. clintoniana* | phased | tau_5t | 0.001317 | 0.000783 | 0.002431 |
| *D. clintoniana* | genotype | tau_5t | 0.000201 | 0.000134 | 0.000267 |
| *D. clintoniana* | consensus | tau_5t | 0.000074 | 0.000006 | 0.000188 |
| *D. clintoniana* | pickone | tau_5t | 0.00006 | 0.000001 | 0.000355 |
| *D. clintoniana* | phased | tau_6s | 0.000689 | 0.000425 | 0.001107 |
| *D. clintoniana* | genotype | tau_6s | 0.000626 | 0.000448 | 0.000818 |
| *D. clintoniana* | consensus | tau_6s | 0.00036 | 0.000021 | 0.000775 |
| *D. clintoniana* | pickone | tau_6s | 0.000419 | 0.000013 | 0.000784 |
| *D. clintoniana* | phased | tau_7h | 0.000632 | 0.000272 | 0.001107 |
| *D. clintoniana* | genotype | tau_7h | 0.000178 | 0.000123 | 0.000234 |
| *D. clintoniana* | consensus | tau_7h | 0.00003 | 0 | 0.000099 |
| *D. clintoniana* | pickone | tau_7h | 0.000047 | 0 | 0.000332 |
| *D. clintoniana* | phased | theta_1Dgol | 0.000654 | 0.000481 | 0.000895 |
| *D. clintoniana* | genotype | theta_1Dgol | 0.000647 | 0.00049 | 0.000801 |
| *D. clintoniana* | consensus | theta_1Dgol | NA | NA | NA |
| *D. clintoniana* | pickone | theta_1Dgol | NA | NA | NA |
| *D. clintoniana* | phased | theta_2Dcri | 0.089117 | 0.056512 | 0.118635 |
| *D. clintoniana* | genotype | theta_2Dcri | 0.002887 | 0.002158 | 0.003631 |
| *D. clintoniana* | consensus | theta_2Dcri | 0.010955 | 0.001595 | 0.027525 |
| *D. clintoniana* | pickone | theta_2Dcri | 0.008274 | 0.000919 | 0.033657 |
| *D. clintoniana* | phased | theta_3Dcli | 0.759117 | 0.216175 | 1.676369 |
| *D. clintoniana* | genotype | theta_3Dcli | 0.001077 | 0.000795 | 0.001357 |

| | | | | | |
|---|---|---|---|---|---|
| *D. clintoniana* | consensus | theta_3Dcli | 0.006674 | 0.001076 | 0.017165 |
| *D. clintoniana* | pickone | theta_3Dcli | 0.021785 | 0.00076 | 0.103124 |
| *D. clintoniana* | phased | theta_4r | 0.058278 | 0.054253 | 0.062397 |
| *D. clintoniana* | genotype | theta_4r | 0.050855 | 0.047451 | 0.054283 |
| *D. clintoniana* | consensus | theta_4r | 0.052871 | 0.049326 | 0.056406 |
| *D. clintoniana* | pickone | theta_4r | 0.054709 | 0.051096 | 0.05864 |
| *D. clintoniana* | phased | theta_5t | 0.034865 | 0.00224 | 0.049053 |
| *D. clintoniana* | genotype | theta_5t | 0.013239 | 0.010949 | 0.015634 |
| *D. clintoniana* | consensus | theta_5t | 0.031073 | 0.022006 | 0.040398 |
| *D. clintoniana* | pickone | theta_5t | 0.144179 | 0.095216 | 0.208439 |
| *D. clintoniana* | phased | theta_6s | 0.059369 | 0.007337 | 0.085735 |
| *D. clintoniana* | genotype | theta_6s | 0.021697 | 0.016597 | 0.027023 |
| *D. clintoniana* | consensus | theta_6s | 0.006642 | 0.001357 | 0.014338 |
| *D. clintoniana* | pickone | theta_6s | 0.003367 | 0.00098 | 0.006741 |
| *D. clintoniana* | phased | theta_7h | 0.06851 | 0.000624 | 0.286324 |
| *D. clintoniana* | genotype | theta_7h | 0.015401 | 0.004426 | 0.033165 |
| *D. clintoniana* | consensus | theta_7h | 0.006885 | 0.000938 | 0.017984 |
| *D. clintoniana* | pickone | theta_7h | 0.005872 | 0.000996 | 0.014861 |
| *D. clintoniana* | phased | theta_8h | 0.036053 | 0.000575 | 0.14961 |
| *D. clintoniana* | genotype | theta_8h | 0.004783 | 0.000799 | 0.012033 |
| *D. clintoniana* | consensus | theta_8h | 0.006054 | 0.001033 | 0.015025 |
| *D. clintoniana* | pickone | theta_8h | 0.00891 | 0.00097 | 0.024465 |

**Table S4 – Posterior Means and HPDs for Parameters from MSci Model 5.** Summary statistics are based on combined posteriors of four independent chains for each data type for each triplet. Parameters correspond to subscripts shown in Supplementary Figure S2.

| | | ***D. campyloptera*** | | ***D. celsa*** | | ***D. clintoniana*** | |
|---|---|---|---|---|---|---|---|
| **Phasing** | ***h*†** | **-Pseudo lnL** | **ΔPlnL** | **-Pseudo lnL** | **ΔPlnL** | **-Pseudo lnL** | **ΔPlnL** |
| Consensus | 0 | 21.10 | NA | 28.16 | NA | 6.42 | NA |
| | 1 | 0.10 | 21.00 | 0.06 | 28.10 | 0.16 | 6.26 |
| | 2 | 0.10 | 0.00 | 0.06 | 0.00 | 0.16 | 0.00 |
| Genotype | 0 | 19.71 | NA | 24.43 | NA | 5.67 | NA |
| | 1 | 0.52 | 19.19 | 0.13 | 24.30 | 0.23 | 5.44 |
| | 2 | 0.52 | 0.00 | 0.13 | 0.00 | 0.23 | 0.00 |
| Phased | 0 | 14.15 | NA | 16.15 | NA | 4.39 | NA |
| | 1 | 0.07 | 14.08 | 0.18 | 15.97 | 0.15 | 4.24 |
| | 2 | 0.07 | 0.00 | 0.18 | 0.00 | 0.15 | 0.00 |
| Pick One | 0 | 18.16 | NA | 27.17 | NA | 5.79 | NA |
| | 1 | 0.21 | 17.94 | 0.55 | 26.61 | 0.15 | 5.64 |
| | 2 | 0.21 | 0.00 | 0.55 | 0.00 | 0.15 | 0.00 |

†Maximum number of reticulation events allowed. Does not require that *h* reticulations be found. Species at the top are the allopolyploid involved in the tests with three taxa.

**Table S5 – PhyloNetworks Results for Analyses of *Dryopteris* Data.** Results are for analyses of all target-enrichment loci. Improvement to the negative Pseudo-loglikelihood score are the previous *h* minus the current *h*. The choice of *h* that reflects the last improvement greater than 2 was used for networks in Figure 6.

| Phasing | *h*† | -Pseudo lnL | Improvement |
|---|---|---|---|
| Phased | 0 | 720.90201 | NA |
| | 1 | 498.069854 | 222.83 |
| | 2 | 362.691802 | 135.38 |
| | **3** | **262.496633** | **100.20** |
| | 4 | 262.46153 | 0.04 |
| | 5 | 262.116879 | 0.34 |
| | 6 | 262.116875 | 0.00 |
| Consensus | 0 | 866.100725 | NA |
| | 1 | 624.106594 | 241.99 |
| | **2** | **485.718856** | **138.39** |
| | 3 | 0 | 485.72 |
| | 4 | 0 | 0.00 |
| | 5 | 0 | 0.00 |
| | 6 | 0 | 0.00 |
| Genotype | 0 | 896.671312 | NA |
| | 1 | 657.892974 | 238.78 |
| | 2 | 511.265498 | 146.63 |
| | **3** | **408.720801** | **102.54** |
| | 4 | 408.715613 | 0.01 |
| | 5 | 402.685359 | 6.03 |
| | 6 | 399.109218 | 3.58 |
| Pick One | 0 | 892.403129 | NA |
| | 1 | 678.22724 | 214.18 |
| | 2 | 555.232715 | 122.99 |
| | **3** | **455.461673** | **99.77** |
| | 4 | 406.99917 | 48.46 |
| | 5 | 396.792253 | 10.21 |
| | 6 | 396.766129 | 0.03 |

†Maximum number of reticulation events allowed. Does not require that *h* reticulations be found. Although the pseudo-loglikelihood improved for some models as h increased, the maximum number of reticulations recovered by any network for that data type is indicated in bold.

**Table S6 – PhyloNetworks Results for Analyses of all North American *Dryopteris*.** Results are for analyses of all target-enrichment loci. Improvement to the negative Pseudo-loglikelihood score are the previous *h* minus the current *h*. The number of reticulations indicated in bold were used for bootstrapping.
